# Inhibitor fluorination pattern tunes chemically induced protein dimerization

**DOI:** 10.1101/2025.07.23.666362

**Authors:** Eric Schwegler, Jean-Martin Harder, Marco D. Preuss, Charlotte Guhl, Shuaibing Zhang, Annika Wagner, Nicole Bader, Pierre Stallforth, Markus Lakemeyer, Hermann Schindelin, Till Opatz, Ute A. Hellmich

## Abstract

Chemically induced dimerization of proteins is a powerful approach to regulate biomolecular functions through small molecule ligands acting as “molecular glues”. Here, we demonstrate that simple, thienopyrimidinone scaffold-based inhibitors efficiently promote homodimerization of an essential oxidoreductase from the human pathogenic parasite *Trypanosoma brucei* through selective covalent attachment and self-assembly. A fluorine walk strategy, commonly used to optimize small molecule properties, resulted in tuning induced dimer affinity across two orders of magnitude. NMR spectroscopy, MD simulations, chromatography, multi-angle light scattering, mass spectrometry, calorimetry, and functional assays reveal how the inhibitor fluorination pattern alters the dynamics and interactions of the enzyme-bound inhibitor and surface-exposed aromatic protein side chains, affecting both enzyme inhibition kinetics and induced dimerization. This work highlights how site-specific fluorination can modulate protein interactions and offers a framework for the design of novel molecular glues with broad applications in chemical biology and drug development.

## Introduction

Chemically induced dimerization (CID) is a powerful tool to modulate the lifetime, localization and biological function of biomacromolecules in vitro and in cellulo.^1–6^ For instance, molecular glues and PROTACs (proteolysis targeting chimeras) for targeted protein degradation have greatly expanded the range of druggable disease targets.^5,7–9^ This strategy is currently applied to cancer, as well as neurodegenerative, inflammatory and infectious diseases, mainly those caused by bacteria or viruses.^5,7–12^ Eukaryotic parasites have traditionally received less attention in drug development, although these organisms are responsible for the majority of the so-called neglected tropical diseases (NTDs).^13^ The World Health Organization (WHO) classifies NTDs as a diverse group of severe medical conditions that affect over a billion people worldwide. This includes Sleeping Sickness in humans (Human African Trypanosomiasis, HAT) and the animal plagues Nagana, Surra and Dourine which are all caused by the unicellular protozoan parasite *Trypanosoma brucei* (*T. brucei*).^14–16^ Therefore, infections with *T. brucei* present a substantial burden not only on healthcare systems, but also on livestock farming, creating considerable economic loss which disproportionally affects marginalized communities in endemic areas.^14–17^

The central enzyme in the redox metabolism of *T. brucei* parasites is tryparedoxin (Tpx), an essential monomeric 16 kDa oxidoreductase with a highly conserved “WCPPC” active site motif that undergoes thiol-exchange reactions.^18,19^ Mutation or covalent modification of the active site nucleophilic cysteine 40 inhibits the enzyme’s redox activity and thus disables the parasite’s defense against oxidative stress, rendering Tpx a potential drug target.^20,21^ The fluorinated, thienopyrimidinone scaffold-based inhibitor 2-(chloromethyl)-5-(4-fluorophenyl)-thieno[2,3-*d*]pyrimidin-4(3*H*)-one (para-CFT, **1**) was previously shown to inactivate the enzyme via covalent modification of C40.^21^ We recently described the para-CFT induced homodimerization of Tpx mediated through an intricate network of intermolecular protein-protein and inhibitor-inhibitor contacts.^22,23^ With a mass of only 295 Da, para-CFT is one of the smallest dimerizers known so far.^1,23,24^

Due to their facile synthetic accessibility, thienopyrimidinone derivatives are well suited for rational drug design and are applied as anticancer, antiviral, antibacterial, and antiparasitic compounds.^25^ To adjust molecular interactions of a small molecule with a biological target, and to optimize metabolic stability and pharmacokinetics, fluorination is commonly used to improve the desired physicochemical properties of pharmaceuticals and agrochemicals.^26–34^ However, except for the serendipitous discovery that the potency of a plant stress receptor small molecule dimerizer was enhanced by adding fluorine substituents^31^, the full potential of fluorination for fine-tuning molecular glue efficiency has not yet been systematically explored.

Here, inspired by earlier studies that used fluorine walks to tune the binding properties of small molecule ligands^26–29,31^, we investigated how fluorination of a covalent thienopyrimidinone inhibitor influences the inactivation kinetics and the chemically induced dimerization of Tpx from *T. brucei*. We found that regardless of the fluorination pattern, all tested molecules had trypanocidal properties and acted as covalent Tpx inhibitors. The unmodified oxidoreductase is monomeric in solution, even at high protein concentrations, but binding of the inhibitors triggered dimerization. Using analytical size exclusion chromatography (SEC), multi angle light scattering (MALS), and isothermal titration calorimetry (ITC), we found that the inhibitor fluorination pattern strongly influenced the strength of dimerization, with induced dimer affinities spanning two orders of magnitude in the micromolar range.

To elucidate the underlying molecular mechanisms for dimerization, we combined functional in vitro assays with molecular dynamics (MD) simulations as well as ^1^H, ^15^N-and ^19^F-nuclear magnetic resonance (NMR) spectroscopy, which takes advantage of the high gyromagnetic ratio and sensitivity to the immediate chemical environment of the ^19^F-nucleus.^22,35–41^ In chemically induced Tpx homodimers, two enzyme-bound inhibitor molecules are sandwiched in the newly formed dimer interface, thereby constituting an essential component of the interaction surface. We identified a small set of intra-and intermolecular contacts that stabilize the interface. These interactions were preserved across all fluorinated inhibitors, regardless of the substitution pattern, highlighting a shared binding mode. However, fluorination altered the dynamics of the bound inhibitor and amino acids in the binding site. Notably, a surface-exposed tryptophan residue emerged as a key modulator of both inhibitor binding and dimer formation, acting as a gatekeeper to either support or hinder interface formation depending on the fluorination pattern.

This study demonstrates that site-specific fluorination can effectively modulate protein-inhibitor interactions, and that even subtle changes in small molecule structure can lead to pronounced effects in chemically induced dimer affinity. These insights offer a promising strategy to control protein-protein interactions with atomic precision using fluorination and establish a molecular framework for optimizing desirable properties of molecular glues in CID applications.

## Results

### Covalent inhibitor fluorination modulates enzyme inactivation kinetics of the essential oxidoreductase Tpx from *T. brucei*

The causative agent of African Sleeping Sickness, *Trypanosoma brucei*, critically relies on an enzymatic peroxide clearance cascade to maintain cellular redox homeostasis and to detoxify peroxides (**Fig. 1a**).^18^ As part of the trypanosomal redox metabolism, the essential oxidoreductase tryparedoxin (Tpx, for sequence and protein structure, see **Supplementary Fig. 1a-b**) transmits electrons to vital enzymes like peroxidases (Px) via a dithiol exchange mechanism (**Fig. 1a**).^18,42,43^ Earlier studies have shown that the Tpx active site residue C40 can be targeted by covalent inhibitors such as the trypanocidal thienopyrimidinone derivative para-CFT (**1, Fig. 1b).** To explore the role of inhibitor fluorination in (trypano)-toxicity, Tpx inactivation kinetics, and Tpx dimerization, we generated a small compound library of para-CFT analogs (**2-5, Fig. 1b**, for inhibitor synthesis and analyses, see **Supplementary Scheme 1** and Extended Material and Methods).

**Fig. 1:**
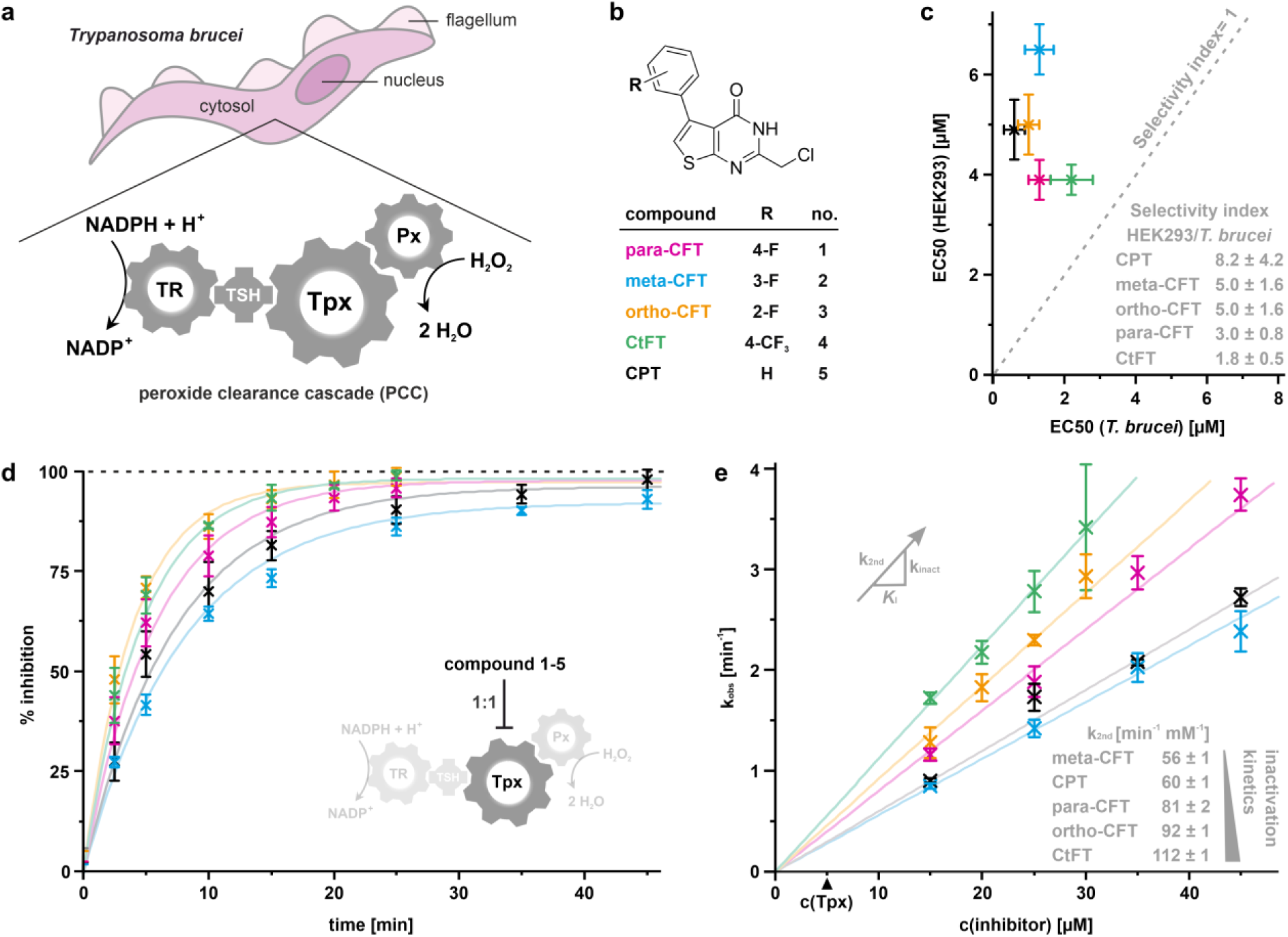
Inhibitor fluorination pattern modulates cytotoxicity and Tpx inactivation kinetics. **a**, Schematic depiction of a unicellular *T. brucei* parasite and its cytosolic peroxide clearance cascade consisting of NADPH, trypanothione reductase (TR), trypanothione (TSH), tryparedoxin (Tpx) and a peroxidase (Px). **b**, Thienopyrimidinone scaffold-based fluorine walk library inspired by para-CFT (**1**), a covalent inhibitor and dimerizer of tryparedoxin (Tpx) from *T. brucei*.^21–23^ para-CFT derivatives (**2**-**5**) are referred to as meta-CFT (2-(chloromethyl)-5-(3-fluorophenyl)thieno[2,3-*d*]pyrimidin-4(*3H*)-one), ortho-CFT (2-(chloro-methyl)-5-(2-fluorophenyl) thieno[2,3-*d*]-pyrimidin-4(*3H*)-one), CtFT (2-(chloromethyl)-5-(4-(trifluoromethyl)phenyl)thieno[2,3-*d*]pyrimidin-4(*3H*)-one), and CPT (2-(chloromethyl)-5-phenylthieno[2,3-*d*]pyrimidin-4(*3H*)-one). **c**, Half-maximal effective concentration (EC_50_) values of **1-5** in cell viability assays with *T. brucei brucei* 449 (strain: Lister 427, bloodstream form) and HEK293 cells (n= 8 and n=4, respectively, see also **Supplementary Table 1**). Data points represent the mean value ± standard deviation.. The inset table shows selectivity indices, calculated as the fraction of the corresponding EC_50_ values for *Trypanosoma* and HEK293 cells (**Supplementary Table 1**). **d**, Time-dependent inhibition curves of Tpx after the addition of Tpx-equimolar amounts of compounds **1-5**. Values were determined through reconstitution of the *T. brucei* peroxide clearance cascade in a cuvette-based assay and photometric measurement of NADPH consumption after H_2_O_2_ addition. Data points represent the mean value ± standard deviation of three technical replicates. **e**, Observed inhibition constants (k_obs_) after the addition of excess amounts of inhibitors **1-5** to 5 µM Tpx (for time-resolved inhibition curves, see **Supplementary Fig. 3a-e**). Data points represent the mean ± standard deviation of three technical replicates. k_obs_ correlated with inhibitor concentration, and linear fits yielded k_2nd_ values (shown on the bottom right, errors arise from the linear fit).

The new synthetic compounds (**2**-**5**) showed activity against bloodstream *T. brucei* similar to the parent compound para-CFT (**1**), with EC_50_ (24 h) values in the low micromolar range from 0.6 ± 0.3 µM for CPT (**5**) to 2.2 ± 0.6 µM for CtFT (**4**) (**Fig. 1c, Supplementary Table 1**). Slightly higher EC_50_ (24 h) values were obtained using human embryonic kidney (HEK293) cells, resulting in selectivity indices, defined as the fraction of EC_50_ values for human vs. parasitic cells, marginally greater than 1 (**Fig. 1c**, inset). This currently precludes a potential use as trypanocidal drugs for these molecules. However, we found them to be highly selective for Tpx in the in vitro reconstituted *T. brucei* redox cascade (**Fig. 1a, Supplementary Fig. 2**). This selectivity was observed despite the high abundance of nucleophilic cysteines from other cascade members, which include a reductase (TR), a peroxidase (Px) and the glutathione-like molecule trypanothione (TSH) (see **Fig. 1a**, cogwheel cartoon).

When inhibitors **1–5** were added in equimolar amounts to Tpx, the entire cascade was fully inactivated (**Fig. 1d**). As expected for selective Tpx inhibition with a 1:1 stoichiometry, addition of 0.5 molar equivalents of inhibitors relative to Tpx resulted in 50% inactivation of the cascade (**Supplementary Fig. 3a-e**). Intact-protein mass spectrometry (MS) on Tpx WT and the active site mutants C40S and C43S confirmed that all inhibitors covalently interact with nucleophilic residue C40, but not C43 (**Supplementary Table 2**). This matched our expectations since active site residue C43, whose primary function is to reset the oxidoreductase to its oxidized state after the reduction of a substrate, is a weaker nucleophile than C40^19^, and thus does not readily interact with covalent inhibitors. In sum, this confirms that the in vitro activity assay provides a direct, time-resolved readout of selective Tpx inactivation, reflecting the extent of enzyme inhibition, and enabling the determination of fluorination-pattern dependent inhibition kinetics of compounds **1-5**.

In the activity assay with Tpx equimolar inhibitor amounts, ortho-CFT and CtFT (**3, 4**) caused the fastest, and meta-CFT (**2**) the slowest Tpx inactivation (**Fig. 1d**). To quantify these differences, we determined the concentration-dependent inhibition constants (k_obs_) for the inhibitors, and derived the corresponding, concentration-independent second-order rate constants (k_2nd_).^44^ These ranged from 112 ± 1 min^-1^ mM^-1^ for CtFT (**4**) to 56 ± 1 min^-1^ mM^-1^ for meta-CFT (**2**) (**Fig. 1e, Supplementary Fig. 3a-e**). Of note, due to the relatively fast inactivation kinetics, our experimental setup precluded us from obtaining k_obs_ at inhibitor concentrations greater than 45 µM, which is required to observe saturation and precisely determine Tpx binding parameters k_inact_ and *K*_I_ for each compound.^44^

Nonetheless, our results show that all tested compounds (**1-5**) covalently inhibit Tpx by interacting with the active site residue C40, thereby enabling a systematic study to elucidate the consequences of molecular glue fluorination for protein-ligand interactions and chemically induced dimerization.

### Inhibitor fluorination pattern modulates induced dimer affinity

Unmodified *T. brucei* Tpx is a monomeric protein that dimerizes only upon binding of para-CFT (**1**) due to the formation of intermolecular protein-protein and inhibitor-inhibitor contacts, as we have previously shown using X-ray crystallography (**Fig. 2a**).^23^ To explore whether compounds **2-5** with alternative fluorination patterns at the inhibitor’s phenyl moiety also induce Tpx dimerization in solution, we initially conducted analytical size exclusion chromatography (SEC), using the elution volumes of unmodified and BM(PEG)_2_-crosslinked Tpx as references for monomeric and dimeric Tpx, respectively (**Fig. 2b, Supplementary Fig. 1c, 4a**).

**Fig. 2:**
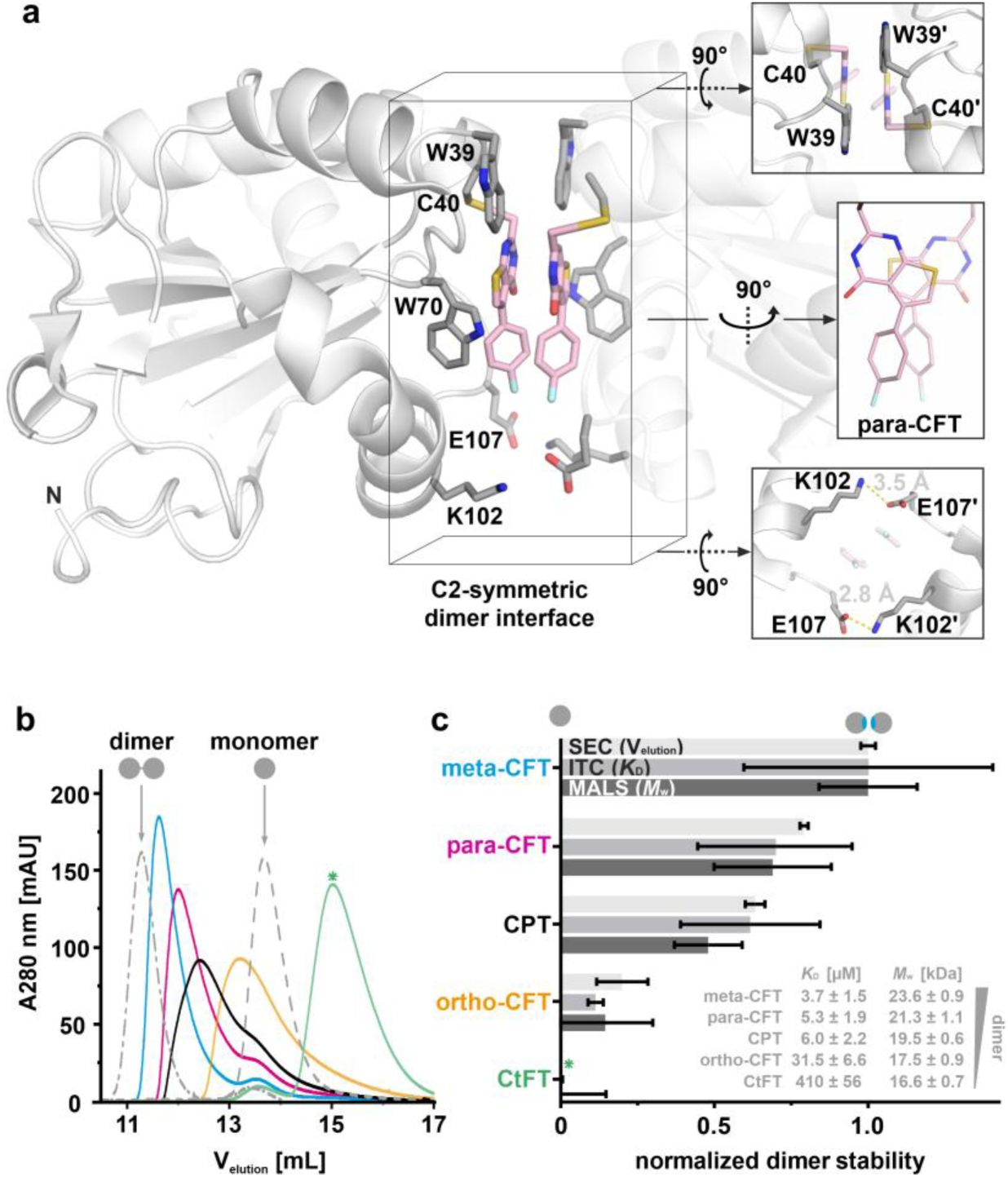
Chemically induced dimerization of Tpx is modulated by the fluorination pattern of the molecular glue. **a**, X-ray crystal structure of the C_2_-symmetric [Tpx/para-CFT]_2_ homodimer (PDB: 6GXG, chains A, B)^23^, highlighting residues W39, K102 and E107, as well as two stacked para-CFT (**1**) molecules in the dimer interface. **b**, Chemically induced dimerization of Tpx (100 µM) by our inhibitor library (colored lines) was probed with SEC, using unmodified (monomer) and BM(PEG)_2_-crosslinked Tpx (dimer) as references (grey dashed lines). An asterisk marks the delayed elution volume seen for Tpx bound to CtFT (**4**) (green trace), which results from increased column interactions (see **Supplementary Fig. 5** and main text for details). **c**, Apparent Tpx dimer stability based on analytical SEC (n =3), SEC-MALS (n = 1), and dilution ITC (n = 3) (**Supplementary Fig. 4, 6**). For comparison of each method, the dimer with the highest affinity, [Tpx/meta-CFT]_2_, was arbitrarily set to a normalized dimer stability of 1. The inset table shows dimer dissociation constants (*K*_D_) from ITC (mean value with combined measurement errors and standard deviation from three technical replicates), and average molecular weights *(M*_w_) obtained from SEC-MALS (value with measurement error). SEC data is presented as mean value ± standard deviation of three technical replicates. A green asterisk signifies the absence of the SEC data of the Tpx/CtFT complex for this plot due to the aberrant SEC running behavior.

Consistent with increased particle size, Tpx bound to inhibitors **1**-**3** and **5** eluted at earlier elution volumes than unmodified Tpx but later than the crosslinked dimer. As expected for a dynamic monomer/dimer equilibrium, elution volumes also varied with analyte concentration and the chromatograms displayed tailing (**Fig. 2b, Supplementary Fig. 4b**). In contrast, Tpx bound to CtFT (**4**) eluted even later than unmodified, monomeric Tpx (**Fig. 2b**, green trace) due to secondary interactions with the SEC column matrix as confirmed by the reversal of this behavior with a more hydrophobic mobile phase (**Supplementary Fig. 5**).

Since the unique SEC running behavior of covalent Tpx/CtFT complexes precluded a straight-forward conclusion about its dimerization tendency, we combined SEC with multi-angle light scattering (MALS). SEC-MALS at 100 µM protein concentration confirmed Tpx dimerization with inhibitors **1**-**3** and **5**, while the Tpx/CtFT (**4**) complex remained monomeric (**Fig. 2c, Supplementary Fig. 4c**). Dissociation constants (*K*_D_) of the chemically induced Tpx dimers derived from dilution ITC revealed a strong dependence on the fluorination pattern, and spanned two orders of magnitude, from meta-CFT (**2**, *K*_D_= 3.7 ± 1.5 µM) to CtFT (**4**, *K*_D_ = 410 ± 56 µM) (**Fig. 2c, Supplementary Fig. 6**).

Thus, combining analytical SEC, SEC-MALS, and ITC measurements, we found that the capability of inhibitors **1**-**5** to act as Tpx dimerizers increased as follows: CtFT (**4**) << ortho-CFT (**3**) < CPT (**5**) < para-CFT (**1**) < meta-CFT (**2**) (**Fig. 2c**). Notably, chemically induced dimerization and inhibition kinetics were inversely correlated (**Supplementary Fig. 3f**), suggesting that *S*-alkylation of the Tpx C40 side chain with strong dimerizers (**1**-**2, 5**) requires more structural changes in the binding interface than *S*-alkylation with weak dimerizers (**3, 4**), which would increase the activation energy of the reaction and thus slow down the binding process.

### The dimer interface is preserved across all Tpx/inhibitor complexes

To investigate the structural consequences of inhibitor fluorination for the induced Tpx dimer interface, we initially aimed to crystallize Tpx with our fluorine walk compound library (**1-5**). However, except for our previously determined Tpx/para-CFT dimer structure (PDB: 6GXG^23^, **Fig. 2a**), this was not successful. Instead, we used ^1^H, ^15^N-NMR spectroscopy to map intramolecular protein-inhibitor as well as intermolecular dimer contacts in solution.^45^ Taking advantage of our previously determined backbone amide resonance assignments for unmodified Tpx in the reduced and oxidized state^46^, the inhibitor-bound states for all five Tpx/inhibitor complexes could be successfully assigned (**Supplementary Fig. 7**). This allowed us to map the respective inhibitor-induced chemical shift perturbations (CSP) onto the Tpx/para-CFT dimer crystal structure (**Fig. 3a, Supplementary Fig. 8**). In all cases, the same residues were affected suggesting that compounds (**1-5**) all induce the formation of similar protein dimer interfaces. Most amide resonances of residues within the dimer interface displayed extensive line broadening, an effect less pronounced for meta-CFT (**2**) and CtFT (**4**) induced dimers with the lowest and highest dimer *K*_D_ values, respectively. This suggested that line broadening is a direct consequence of monomer/dimer exchange and modulated by the population of the respective state. For residues with linear chemical shift trajectories (exemplified for residues A11, L112, and R128 in **Fig. 3b**, left boxes), their extent largely mirrored the relative dimer stability determined by SEC, SEC-MALS and ITC, indicative of a gradual transition from monomer to dimer across the titration.

**Fig. 3:**
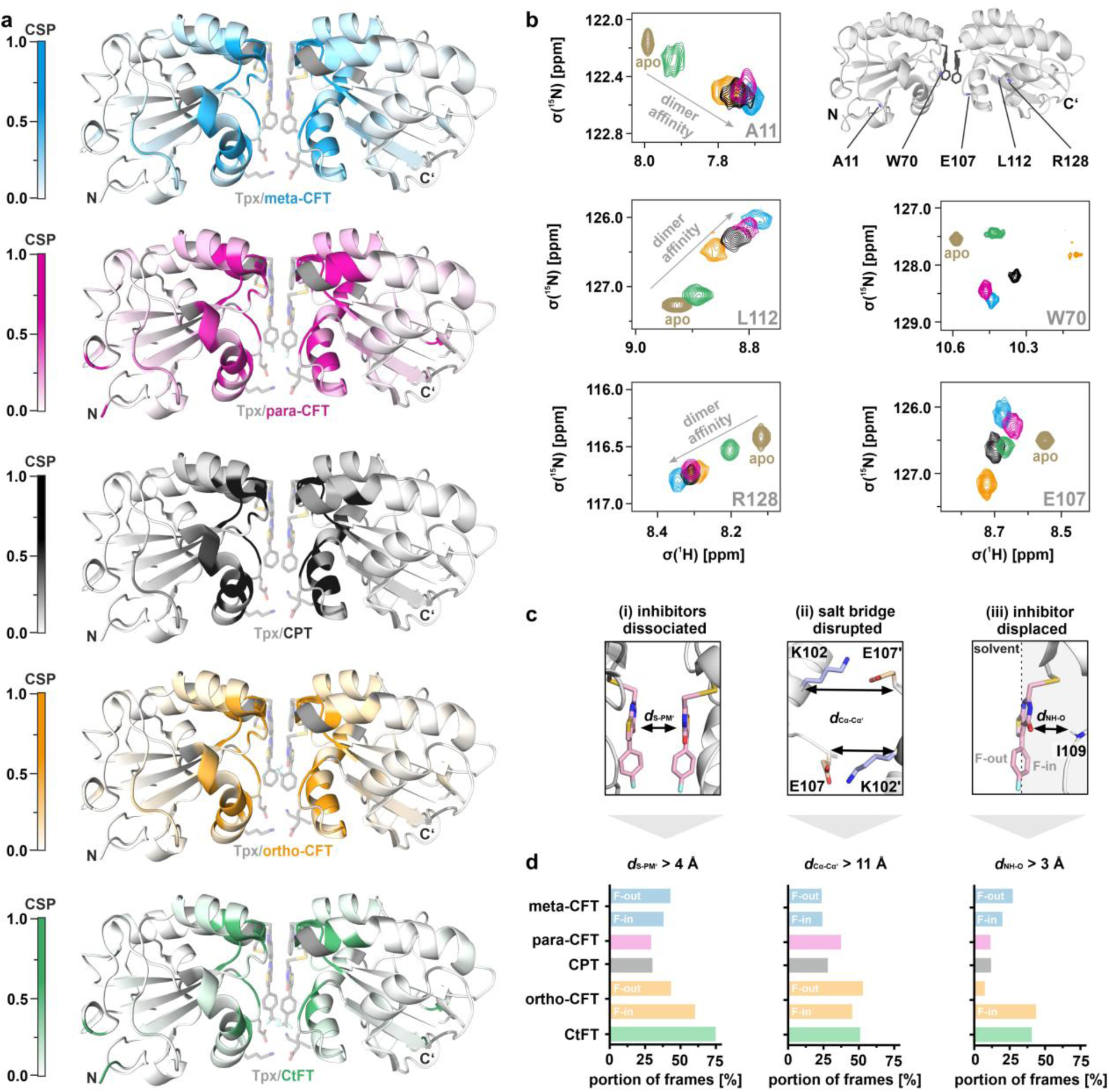
Chemically induced Tpx dimers share a common interface that is stabilized by a small set of inter-and intramolecular contacts. **a**, Chemical shift perturbation (CSP) for Tpx upon binding of inhibitors **1-5**, mapped onto the Tpx/para-CFT dimer X-ray structure (PDB: 6GXG^23^) (see **Supplementary Fig. 7, 8** for full ^1^H, ^15^N-HSQC spectra, per-residue CSP, and “open book” representation). Tpx dimers with bound meta-and ortho-CFT (blue and orange interface) are arbitrarily shown here in “F-in”-and “F-out”-conformation, respectively. **b**, Selected resonances from ^1^H, ^15^N-HSQC spectra of Tpx in the reduced apo (sand) and inhibitor-bound states (colored). Most amide resonances in the dimer interface display line broadening or a linear trajectory matching induced dimer affinities, exemplarily shown for A11, L112 and R128. NH resonances of residues in immediate contact with the inhibitors, e.g. from the W70 side chain and the E107 backbone, did not show this trend as they are affected both by inhibitor binding and dimerization (see main text for details). **c**, The dimer interface was probed by molecular dynamics (MD) simulations (five replicates of 100 ns simulations each, **movies 1-7**) using the Tpx crystal structure with para-CFT (**1**) (6GXG, chains A and B^23^) as a starting model. For meta-and ortho-fluorinated compounds (**2, 3**), two different phenyl ring conformations with fluorine facing either towards the protein (“F-in”) or the solvent (“F-out”) were considered (see main text and **Supplementary Fig. 9, 10** for details). Tracking three key intra-and intermolecular contact distances during the simulations was sufficient to correlate MD frame populations with the dimer affinities measured in vitro (Fig. 2c): (i) inhibitor stacking (monitored by *d*_S-PM_), (ii) interchain salt bridges between K102 and E107’ (monitored by *d*_Cα-Cα’_), and (iii) inhibitor desorption from the protein surface (monitored by *d*_NH-O_). For details, see Extended Material and Methods and main text. **d**, The number of frames where threshold distances *d*_S-PM_, *d*_Cα-Cα’_, and *d*_NH-O_ exceeded critical values, deemed incompatible with stable dimer formation, were quantified. The weak dimerizers ortho-CFT and CtFT (**3, 4**) exceeded these critical thresholds most often.

Interestingly, amide resonances of Tpx residues which, based on the crystallized Tpx/para-CFT complex^23^ (**Fig. 2a**), were presumed to participate in direct inhibitor interactions, did not show the linear chemical shift behavior (**Fig. 3b**, right boxes). Examples include the W70 side chain which interacts with the inhibitor’s thiophene moiety via T-shaped π-interactions, and the E107 backbone which contacts the inhibitor’s phenyl substituent (**Fig. 2a**).^23^ Although it is tempting to interpret the chemical shift differences across inhibitors as direct evidence for fluorination pattern induced differences in inhibitor binding modes, these signals may nonetheless also be affected by dimerization. For instance, a similar E107 chemical shift for CPT and CtFT (**4, 5**) does not necessarily imply the same binding mode, since **5** is a strong Tpx dimerizer, whereas **4** is not. Furthermore, both inhibitors (**4, 5**) induced markedly different chemical shift perturbations on the W70 side chain NH resonance, likely reflecting differences in dimer affinity, inhibitor interactions, or both (**Fig. 3b**). To further probe the interface of the chemically induced dimers, we thus complemented our NMR studies with molecular dynamics (MD) simulations.

### Dimer stability is governed by a small set of inter-and intramolecular contacts

To set up MD simulations with energetically favorable inhibitor conformations, we first used density-functional theory (DFT) calculations of the isolated inhibitors to assess the energy barriers associated with rotating the dihedral angle between the phenyl and thiophene rings (**Supplementary Fig. 9a**). All inhibitors (**1**-**5**) displayed two rotational barriers at torsion angles of 0° and ∼180° between the phenyl and thiophene rings. For ortho-CFT (**3**), a steric clash between the inhibitor’s fluorine and oxygen atoms resulted in a particularly high energy barrier (∼80 kJ/mol). For all molecules, we observed local energetic minima at dihedral angles of ∼48°. Since this matched with the 47° dihedral angle observed between the phenyl and thiophene rings in the crystal structure of Tpx-bound para-CFT (**1**) (**Fig. 2a**)^23^, and was in line with crystal structures of the isolated inhibitors that also showed non-planar conformations (**Supplementary Fig. 9b-d, Supplementary Table 3**), we considered a dihedral angle of 47° as a suitable starting point for our MD simulation of Tpx in complex with all inhibitor molecules.

For Tpx-bound meta-and ortho-CFT (**2, 3**), which have an asymmetric phenyl substitution pattern and become atropisomers as soon as their phenyl ring rotation is restrained, we defined two different inhibitor orientations, “F-in” and “F-out”, where the inhibitor fluorine atom either faces towards the binding pocket or towards the solvent (**Fig. 3c**, right panel). To consider this, we set up three different MD simulation series (n = 5, 100 ns each) for meta-and ortho-CFT induced Tpx dimers with both protomers in the “F-in”, both protomers in the “F-out”, or both with different orientations (**Supplementary Fig. 10, movies 1-7**). In line with the notion that the interaction with Tpx strongly increases the rotational barriers of the inhibitor molecules and locks the phenyl substituent’s conformation to create atropisomers, axial chirality of Tpx/inhibitor complexes was retained throughout all five simulations. A comparison between MD simulations of dimeric and monomeric Tpx WT/inhibitor complexes (for meta-and ortho-CFT simulated in “F-in”-and “F-out”-conformation, respectively) showed that the conformational “trapping” of the inhibitor molecule is a result of Tpx binding, not subsequent dimerization. Nonetheless, dimerization further restrains the accessible conformational space of the inhibitors’ phenyl substituents (**Supplementary Fig. 10, movies 8-14**).

Across all MD simulations, a small set of one intra-and two inter-molecular interactions was sufficient to correlate relative dimer stabilities obtained from SEC, SEC-MALS, ITC, and ^1^H, ^15^N-NMR spectroscopy with dimer contacts observed in silico: (i) intermolecular inhibitor stacking represented by the distance between the thiophene sulfur and pyrimidinone centroid of the two sandwiched molecules (*d*_S-PM’_), (ii) an intermolecular salt bridge between residues K102/E107’ (and K102’/E107) (*d*_Cα-Cα’_), and (iii) the desorption of the inhibitor from the binding site, represented by the increase in distance between the thienopyrimidinone core and the I109 backbone NH (*d*_NH-O_) (**Fig. 3c, d, movies 1-7**, see Extended Material and Methods for details). Exceeding critical thresholds for *d*_S-PM_, *d*_Cα-Cα’_ and *d*_NH-O_ (> 4 Å, > 11 Å, and > 3 Å, respectively) was deemed to be incompatible with a stable dimer arrangement and interpreted as partial dimer dissociation (see **Supplementary Fig. 11** for exemplary frames of intact and partially dissociated dimer interfaces).

In agreement with our in vitro data, the least potent dimerizers ortho-CFT and CtFT (**3, 4**), exhibited the largest number of frames with “partially dissociated” interfaces (**Fig. 3d**, orange and green bars, **movies 4-6**). Notably, desorption of the thienopyrimidinone moiety of ortho-CFT (**3**) occurred more often for the “F-in”-conformation than for the “F-out” conformation (**Fig. 3c-d, movies 4-5**), suggesting that ortho-fluorination directly interfered with inhibitor interactions in the binding pocket, a finding also supported by ^19^F-NMR (see below). Such behavior may also result in distinct dynamic responses in neighboring amino acids such as the W70 side chain, resulting in e.g. line broadening of the corresponding indole NH resonance (**Fig. 3b**, top right panel).

Overall, all new molecular glues (**2-5**) induced highly similar Tpx dimer interfaces as the parent compound para-CFT (**1**). However, depending on the respective inhibitor fluorination pattern, our NMR and MD analyses revealed local variations in intra-and intermolecular dimer interactions involving the molecular glue itself as well as protein residues surrounding the inhibitor binding site. These differences presumably hold the key to understanding the observed differences in dimer affinity.

### Inhibitor fluorination mediates specific interaction profiles in a common binding pocket

Our MD simulations indicated that intramolecular protein/inhibitor contacts (e.g. *d*_NH-O_ in **Fig. 3c-d**) significantly contribute to the stability of the chemically induced Tpx dimers. Because inhibitor-bound Tpx WT exists in a monomer-dimer equilibrium, it is difficult to study inhibitor binding to the monomer in the presence of concurrent dimerization. To nevertheless investigate how inhibitor fluorination affects interactions with the Tpx binding pocket, we used the W39A mutant that still covalently binds all inhibitors as seen by intact-protein MS (**Supplementary Table 2**), but was previously shown to greatly reduce dimerization upon para-CFT binding.^22,23^ SEC analysis confirmed that also in the presence of the new molecular glues (**2-5**), there is no appreciable dimerization of the Tpx W39A mutant (100 µM protein, **Supplementary Fig. 12**), thereby enabling the investigation of monomeric Tpx W39A/inhibitor complexes by ^1^H, ^15^N-NMR spectroscopy (**Fig. 4**).

**Fig. 4:**
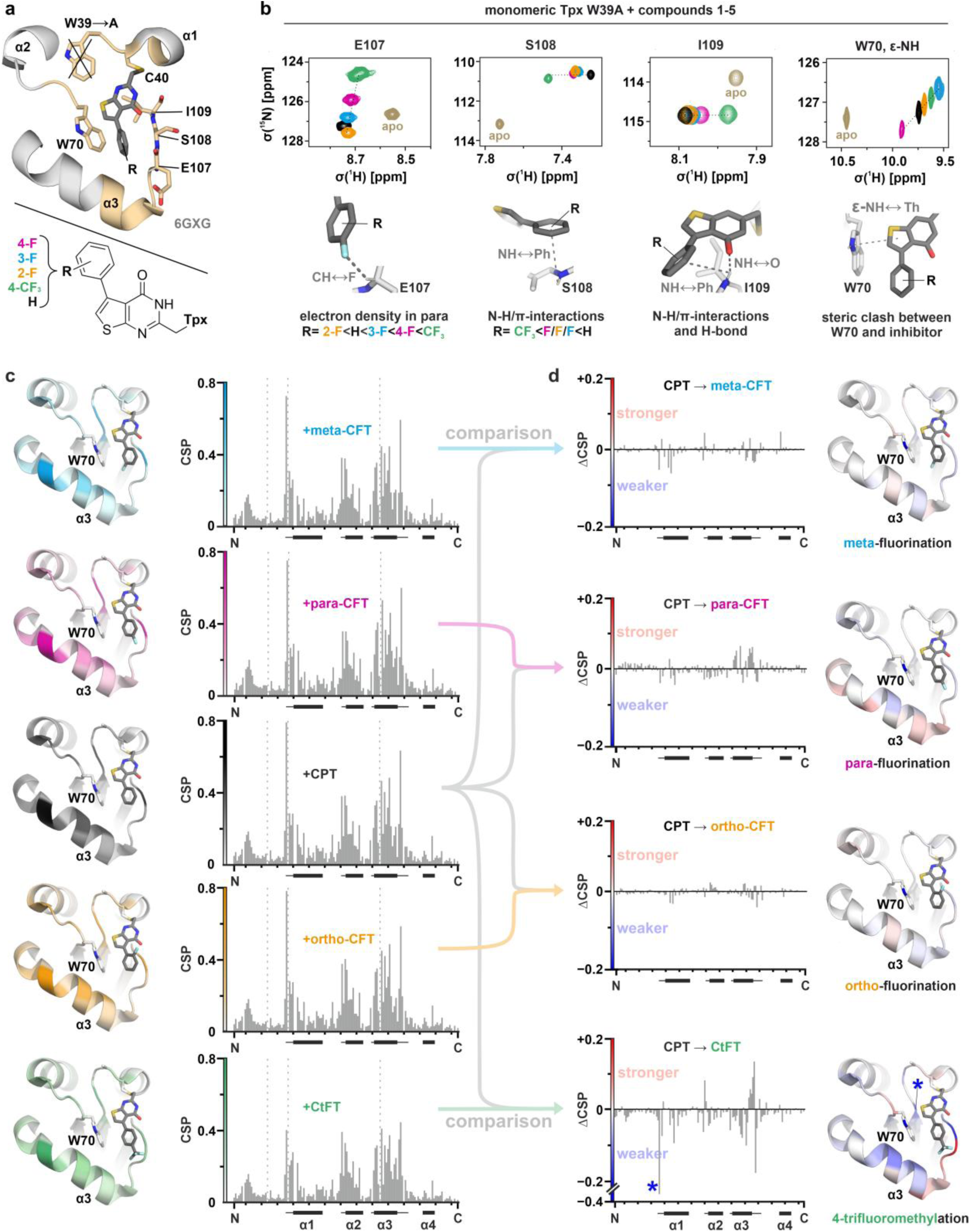
Inhibitors bound to monomeric Tpx W39A occupy a consensus binding pocket but display fluorination pattern-specific protein interactions. **a**, Structural model of the monomeric TpxW39A/inhibitor complex showing the consensus inhibitor binding pocket (sand) for covalently attached compounds **1-5**, derived from the chemical shift perturbation pattern in ^1^H,^15^N-NMR experiments and MD simulations (Fig. 2a**, Supplementary Fig. 14, 15a**). Heteroatom color scheme: N: blue; O: red; S: yellow; F: light blue. **b**, Amide resonances of selected amino acids reflect interactions with the inhibitors’ fluorine, phenyl, oxygen, and thiophene moieties (for details see main text). Hydrogen bonds and aromatic interactions are indicated with dashed lines. **c**, CSPs of monomeric Tpx W39A upon inhibitor binding mapped onto the structural model from (**a**) (left) and the protein sequence (right). CSP per residue is shown as bar graphs, line broadening is indicated by dashed bars. The plot starts with residue G1 and ends with N144 (indicated with “N” and “C”, respectively), ticks indicate increments of ten, starting with G10. The Tpx α-helices are indicated by black cylinders on the x-axis, and adjacent loops with high CSP are indicated by lines. **d**, Pairwise chemical shift perturbation difference (ΔCSP) between the non-fluorinated ligand CPT (**5**, grey arrows) and its respective fluorinated analogs (**1-4**, colored arrows) plotted on the sequence (left) and the structural model from (**a**) (right). Values of 0 mean no CSP difference between fluorinated and non-fluorinated inhibitor (shown in white on the structure). Values greater or lower than 0 indicate an enhanced or reduced effect on the respective residue and are a direct consequence of fluorination (shown in red and blue, respectively). Inhibitor interactions with residue S36, located in the back of the binding pocket and indicated by an asterisk (*), are greatly reduced for the trifluoromethylated inhibitor CtFT (**4**).

First, we assigned the backbone amide resonances of unmodified and inhibitor-bound Tpx W39A and determined the corresponding chemical shift perturbations induced by each inhibitor (**Supplementary Fig. 13, 14**). These data were used to define a consensus binding pocket for monomeric Tpx (**Fig. 4a**, sand color). This pocket comprises the N-terminal loops of α-helix1 (active site) and α-helix2 (containing W70), as well as the C-terminal end of α-helix3 and the adjacent loop containing residues E107, S108, and I109 (**Fig. 4a**). Notably, this binding pocket matches the intramolecular contacts observed in the inhibitor-bound dimer crystal structure (6GXG^23^), and was also reproduced in the MD simulations of monomeric Tpx W39A with inhibitors **1-5** (**Supplementary Fig. 15a**).

Despite their shared binding site, differences in inhibitor fluorination led to distinct effects on individual residues. In many cases, the chemical shift trajectories directly reflect the inhibitor fluorination pattern and the resulting electron density distribution of the fluorophenyl ring (**Fig. 4b, Supplementary Fig. 14**). For example, the E107 backbone amide chemical shift trajectory depended on the electron density at the inhibitor’s para position. This is highest for CtFT (**4**, green) due to its electron-dense CF_3_-group, and lowest for ortho-CFT (**3**, orange), where the Fluorine-substituent deprives the inhibitor phenyl ring of electron density in the para position, a loss that except for meta-CFT (**2**), cannot be compensated via the mesomeric (+M) effect. Similarly, S108 backbone amide shielding correlated with the electron density of the inhibitor’s phenyl ring, an effect most pronounced for the non-fluorinated, electron-rich phenyl moiety in CPT (**5**, black), and least pronounced for CtFT (**4**, green) with its electron-poor phenyl π-system. Not surprisingly, the S108 backbone NH chemical shift was nearly identical for all monofluorinated inhibitors **1-3**. The I109 backbone amide trajectory probably resulted from a combination of hydrogen bonding and NH/π-interactions with the inhibitor, and intriguingly showed the opposite trajectory compared to the S108 residue. We propose that selective shielding and deshielding of the two neighboring residues S108 and I109 by a magnetically induced inhibitor phenyl ring current could explain this phenomenon (**Supplementary Fig. 15b**). W70 side chain NH shifts did not correlate with the inhibitor’s phenyl ring electron density, likely due to side chain rotations induced by the inhibitor as seen in our crystal structure of the Tpx/para-CFT complex, that can strongly affect interactions with the inhibitor (**Supplementary Fig. 15c**). Overall, the fluorination pattern directly affects how an inhibitor interacts with the binding site residues. Importantly, these effects can be graded as seen here for selected examples based on the electron density of the phenyl ring or the position of the fluorine group.

### Inhibitor fluorination controls depth of binding pocket engagement

To assess fluorination-dependent protein surface engagement in the absence of dimerization, we compared the chemical shifts of monomeric Tpx W39A bound to the nonfluorinated inhibitor CPT (**5**) and to fluorinated inhibitors (**1–4**). Since CPT (**5**) shares the core scaffold with inhibitors (**1–4**) but lacks fluorine, any changes in chemical shift perturbations (ΔCSP) should therefore reflect fluorination-dependent interactions with the protein surface.

To this end, we first mapped the backbone chemical shift perturbations (CSP) of each inhibitor onto the protein structure and sequence individually (**Fig. 4c**). As observed previously with Tpx WT (**Fig. 3a**), CtFT (**4**) caused the smallest overall chemical shift changes in monomeric Tpx W39A (**Fig. 4c, bottom**), suggesting that the contacts between the inhibitor and the protein are more limited in this case compared to the other molecules due to the bulky trifluoromethyl group.

Next, we determined the CSP differences (ΔCSP) between the non-fluorinated inhibitor CPT (**5**) and each of the fluorinated analogs **1-4** (**Fig. 4d**). Regions with stronger or weaker CSPs for the fluorinated compounds relative to CPT (**5**) were mapped onto the protein structure in red and blue, respectively. The largest differences between fluorinated analogs and CPT were found for α-helix3 and the adjacent loop, suggesting that this is the main contact site for the inhibitor’s phenyl ring with the protein surface. Ortho-and meta-fluorination had only minor effects, while para-substitutions were strongly “sensed” by the α-helix3 and the succeeding loop. The trifluoromethyl group in CtFT (**4**) weakened the interactions (negative ΔCSP, blue) with the middle of α-helix3 and at the back of the binding pocket, notable through loss of contact with residue S36 (marked with an asterisk in **Fig. 4d**, bottom), suggesting “looser” adsorption of the inhibitor’s thienopyrimidinone moiety to the protein surface. At the same time, stronger interactions with residues in the C-terminal loop of α-helix3 on the protein surface (positive ΔCSP, red) were observed for this inhibitor (**4**) consistent with it occupying a less deeply buried binding position.

Together, our combined NMR and MD analyses with the monomeric Tpx W39A mutant showed that all inhibitors adopt a similar orientation in the Tpx binding site, but with distinct protein surface adsorption behaviors, in particular for the trifluoromethylated compound CtFT (**4**).

### The W39 side chain engages in a “steric quarrel” with covalently bound inhibitors

Our data suggested that the “loose” binding pose of CtFT (**4**) may be the reason for its weak dimerizing capabilities. However, structural or dynamic differences between ortho-CFT (**3**) with an intermediate *K*_D_ for dimerization and the strong dimerizers (**1, 2, 5**) were not as readily apparent from the protein NMR spectra of the Tpx W39A mutant. We thus wondered whether the side chain of W39 may play a non-negligible role in the positioning of molecular glues in the Tpx dimer interface.

Advantageously, the low dimer affinity for CtFT (**4**) allowed us to directly compare the inhibitor-dependent CSPs for monomeric Tpx WT and W39A and revealed stronger interactions of CtFT (**4**) with the mutant (**Fig. 5a-b**, highlighted red). Tpx WT and the W39A mutant CSP cannot be directly compared in the same way of the other molecular glues, as they primarily reflect the respective differences in dimer affinity (**Supplementary Fig. 16**). We thus ran MD simulations of monomeric Tpx WT and W39A inhibitor complexes (**Fig. 5c, Supplementary Fig. 17, movies 8–14**). Consistent with the NMR data for CtFT (**4**), the covalently attached ligands adopted more stable binding poses in the absence of W39, whose indole side chain may otherwise restrict access to the binding pocket (**Fig. 5c**).

**Fig. 5:**
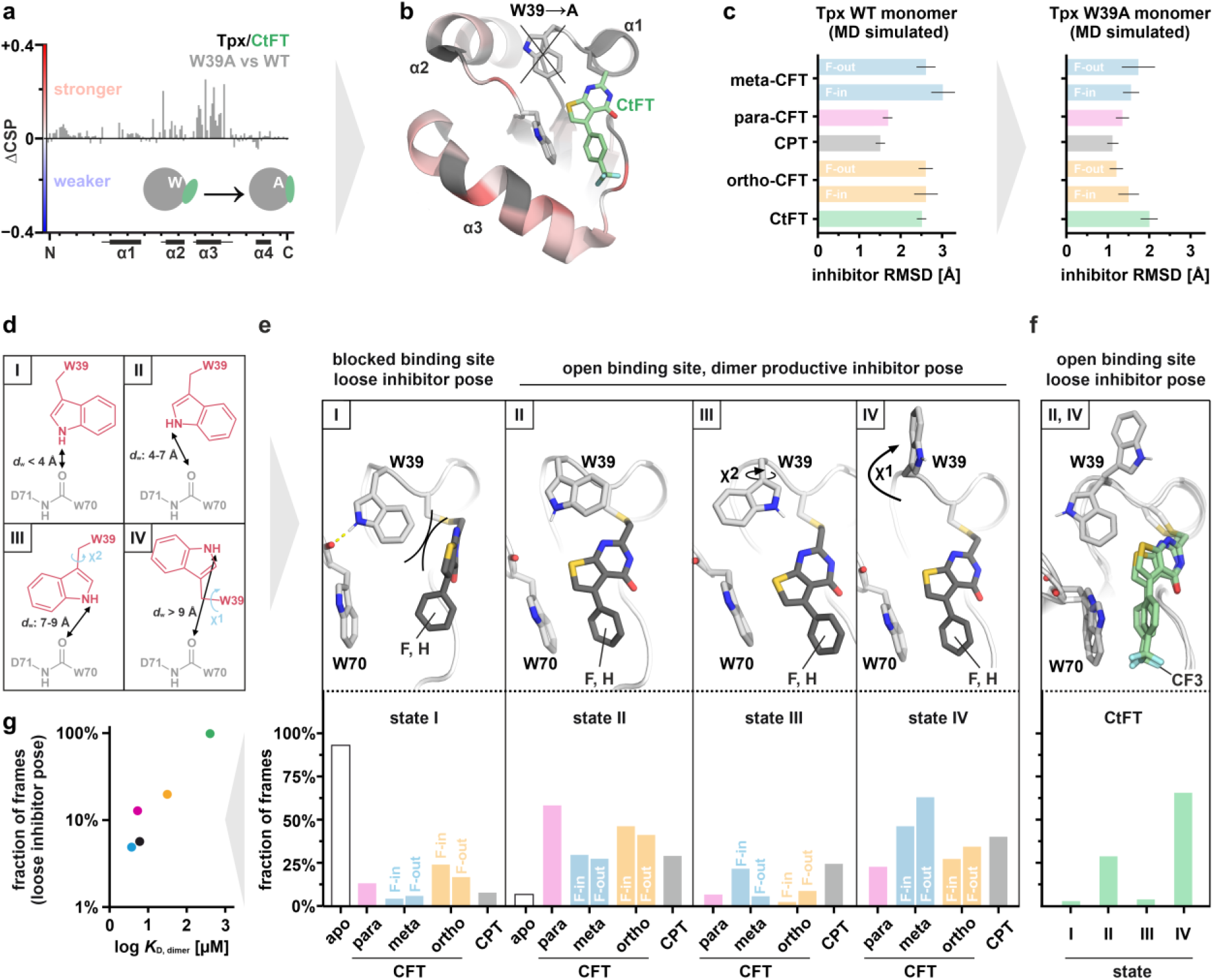
The Tpx W39 side chain modulates inhibitor surface attachment. **a, b**, Differences in chemical shift perturbation (ΔCSP) between Tpx WT and W39A bound to CtFT (**4**) mapped on the sequence (**a**) and structure of Tpx (**b**) show that loss of the W39 side chain in the mutant resulted in stronger inhibitor adsorption of (**4**) to the Tpx surface compared to the WT (red regions). **c**, Inhibitors are more dynamic bound to Tpx WT than to the W39A mutant, i.e. show a higher RMSD compared to the starting structure (PDB: 6GXG^23^, chain A) in MD simulations of monomeric Tpx/inhibitor complexes (100 ns, n = 5). **d**, Across available Tpx structures, W39 displays diverse conformations, here clustered into states **I-IV** based on the distance between the W39 indole NH proton and the W70 carbonyl oxygen (*d*_w_, indicated by a double arrow), and the W39 dihedral angles χ1 and χ2. **e**, Representative conformations of inhibitors and W39, W70 side chains in states I-IV from MD simulations of monomeric Tpx WT/inhibitor complexes (100 ns, n = 5, **Supplementary Fig. 17, movies 8-14**). At the bottom, the fraction of frames seen across all MD simulations is shown. **f**, For CtFT (**4**), only states II and IV were found to be significantly occupied and in both resulting in a “loose” binding stance of the inhibitor. Heteroatom color scheme: N: blue; O: red; S: yellow; F: light blue. **g**, Inhibitor-induced Tpx dimer affinity from in vitro experiments correlates with the fraction of MD simulation frames where the respective molecular glue adopts a loose, dimer disrupting stance.

Comparison of available crystal structures for oxidized^47^ and inhibitor-bound^23^ *T. brucei* Tpx WT, the AlphaFold2 model of the oxidized enzyme^48,49^ and our MD simulations revealed substantial structural plasticity for the W39 side chain. The W39 sidechain orientations could be clustered into four main states (I-IV) (**Fig. 5d**): In state I, the W39 side chain forms a hydrogen bond with the W70 backbone carbonyl, thereby adopting a conformation that effectively occludes the binding pocket. In states II-IV, the hydrogen bond is missing, resulting in a reorientation of the W39 side chain (state II), a flip “sideways” around the χ_2_ angle (state III) or a flip “upward” around the χ_1_ angle (state IV), thereby completely detaching W39 from the protein surface. Notably, once state III was adopted, it persisted throughout the remainder of the MD simulations, suggesting an energetic minimum (**movies 10, 13**).

To quantify the occupancy of states I-IV across inhibitor-bound complexes, we tracked the distance between the W39 indole NH and the W70 backbone oxygen (*d*_w_) in our MD simulations. Distances of *d*_w_ < 4 Å were defined as state I, between 4-7 Å as state II, between 7-9 Å as state III, and > 9 Å as state IV, respectively (**Fig. 5d, Supplementary Fig. 17**, see Extended Material and Methods for details).

In the absence of inhibitors, 92% of frames were in state I with a “closed” binding pocket (**Fig. 5e, white bars**). This “closed” conformation is also observed in the AlphaFold2 model^48,49^ of apo state *T. brucei* Tpx and crystal structures of orthologous Tpx from other trypanosomatid species^50,51^ (shown in **Supplementary Fig. 13d-e**). Spontaneous “opening” of the binding pocket (state II) occurred in only 8% of MD frames, while states III and IV were never seen without an inhibitor.

In contrast, inhibitor-bound proteins showed diverse W39 conformations (**Fig. 5e-f, colored bars**, see **Supplementary Fig. 17** for snapshots of explicit W39 side chain and inhibitor conformations): Across all inhibitors, Tpx/ortho-CFT complexes occupied the “closed” conformation of the binding pocket (state I) most frequently (**Fig. 5e, orange bars, movies 11, 12**). Incidentally, the W39 side chain orientation prevented thienopyrimidinone adsorption, resulting in an inhibitor pose similar the “loose” conformation we previously saw for CtFT (**Fig. 4d, bottom, 5f, Supplementary Fig. 17**). Nonetheless, ortho-CFT (**3**) also populated the other states II-IV, which allowed for inhibitor adsorption to the binding pocket. The combination of both “loose” and tightly bound inhibitor stances may thus be consistent with the intermediate dimer affinity observed for ortho-CFT (**3**), indicating that inhibitor access to the binding pocket, rather than W39 orientation per se, defines dimer affinity (**Fig. 5g**). Consistent with this notion, our MD simulations showed that stable dimer interfaces can form regardless of the W39 χ-angles (**Supplementary Fig. 11, movies 1-7**) and that both the strongest and weakest dimerizers meta-CFT (**2**) and CtFT (**4**) favor state IV (**Fig. 5e-f, blue and green bars**). However, while meta-CFT (**2**) efficiently inserts into the binding pocket when the W39 side chain has cleared the way, the bulky CF_3_-group in CtFT (**4**) still prevented adsorption and thus a dimerization-promoting inhibitor stance (**Supplementary Fig. 17)**.

Further supporting the idea that tight attachment of an inhibitor to the protein surface is the main driver for efficient dimerization, Tpx-bound para-CFT (**1**) and CPT (**5**) with near identical dimer affinities preferred different W39 orientations that all nonetheless allowed for efficient inhibitor adsorption. Para-CFT (**1**) strongly favored state II, possibly stabilized through hydrogen bond formation between the fluorine in para-position and the Cα proton of E107 (**Fig. 2a**).^23^ In contrast, Tpx/CPT complexes showed no clear preference between state II-IV (**Fig. 5e, pink, grey bars**), suggesting multiple W39 conformations can support effective inhibitor adsorption.

Overall, the MD simulations showed that the weak dimerizers (**3, 4**) more often adopted “loose” binding poses, i.e. with inhibitors not adsorbed to the protein surface, compared to the strong dimerizers (**1, 2, 5**). Importantly, the fraction of such poses directly correlated with dimer affinity (**Fig. 5g**). Together with the ^1^H, ^15^N-NMR data, the MD simulations suggest that W39 and the inhibitor compete for the protein binding pocket, resulting in a “molecular quarrel” where W39 modulates inhibitor adsorption to the binding pocket and thereby dimerization efficiency.

### Fluorination modulates bound inhibitor dynamics and protein dimerization

Our MD simulations suggested that the gatekeeping residue W39 engages bound inhibitors in a “molecular quarrel” because both compete for the same binding pocket. This should result in differences in inhibitor dynamics for different fluorination patterns depending on the presence or absence of W39 (**Fig. 5e**). With its high gyromagnetic ratio and exceptional chemical shift sensitivity, ^19^F is a powerful NMR probe.^35,41^ Here, we took advantage of the fluorine groups in inhibitors **1-4** to investigate the Tpx/inhibitor complexes from the compounds’ perspective (**Fig. 6a, Supplementary Table 4**).

**Fig. 6:**
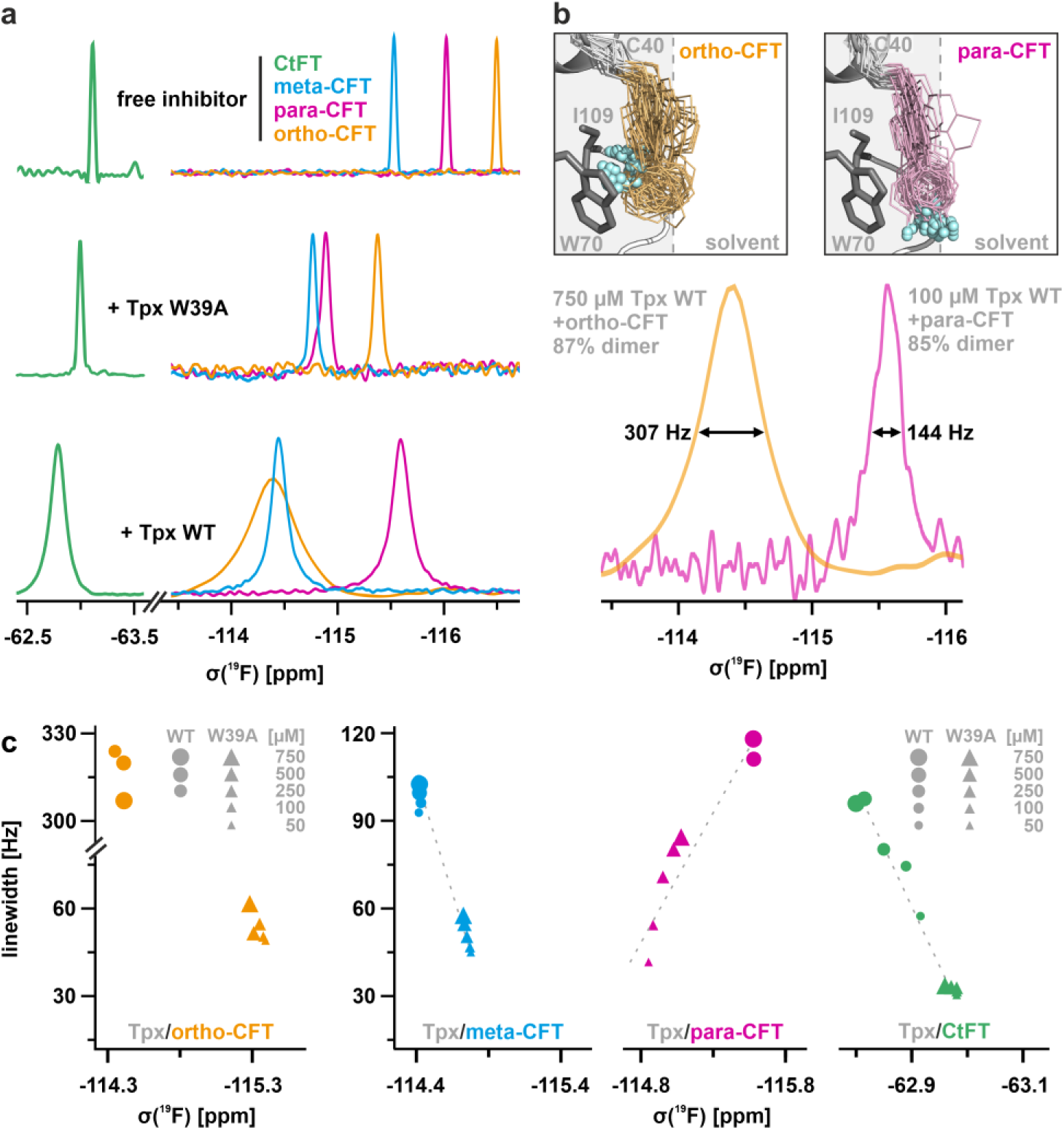
Protein-bound inhibitor dynamics probed by ^19^F-NMR. **a**, 1D ^19^F-NMR spectra of fluorinated Tpx inhibitors **1-4** in the free (100 µM), Tpx W39A-bound (100 µM) and Tpx WT-bound states (at 750 µM to strongly favor the dimeric state). **b**, Tpx WT-bound para-and ortho-CFT (**1, 3**) in MD simulations (top) and ^19^F-NMR spectra (bottom) at similar monomer:dimer ratios. The depicted conformational inhibitor ensembles stem from inhibitor orientations in 33 MD frames, evenly sampled over one 100ns simulation. Ortho-CFT (**3**) is shown here in the “F-in”-conformation. The W70 and I109 side chains are shown in grey. **c**, Linewidths of ^19^F-NMR signals of Tpx-bound compounds **1-4** at different protein concentrations (**Supplementary Fig. 18, Supplementary Table 4**). Due to poor signal to noise, data points of Tpx WT with bound para-and ortho-CFT (**1, 3**) at low concentrations (50 and 100 µM, respectively) are not shown.

Overall, covalent modification of either Tpx WT or W39A with molecular glues **1-4** resulted in ^19^F-chemical shift changes compared to free inhibitors, and increased linewidths in agreement with the increased molecular weight upon covalent complex formation (**Fig. 6a, Supplementary Table 4**). As expected, the monomeric Tpx W39A/inhibitor complexes had narrower linewidths than the corresponding Tpx WT dimers. In line with our complementary ^1^H, ^15^N-NMR and MD data, the ^19^F-NMR signals of free and protein-bound CtFT (**4**) differed the least, likely due to the solvent exposed CF_3_-group, the low dimerization capability, and/or the lower sensitivity of CF_3_ groups to changes in chemical environment compared to single fluorine substituents (**Fig. 6, green lines and symbols**).^39^

Signals from ortho-CFT (**3**) were exceptionally broad when bound to Tpx WT but not the W39A mutant, suggesting an outsized role of this residue for inhibitor dynamics (**Fig. 6, orange lines and symbols**). The observed linewidth in the WT could be due to the exchange dynamics between monomer and dimer, or due to the intrinsic dynamics of the protein-bound inhibitor (**3**). Under our experimental conditions we expected ∼87% dimer species for the Tpx WT/ortho-CFT complex based protein concentration and dimer *K*_D_ value. As a comparison, we also recorded the ^19^F-NMR spectrum of the Tpx WT/para-CFT complex at a similar monomer/dimer ratio (∼85% dimer species) by adjusting the protein concentration to the dimer affinity (**Fig. 6b, bottom**). Here, the signal remained significantly narrower, suggesting that the difference in linewidth for the two molecules reflects differences in protein-bound inhibitor dynamics. Our MD simulations suggest that the fluorine substituent in ortho-CFT (**3**), particularly in the “F-in”-conformation, is in a heterogeneous chemical environment due to inhibitor dynamics, while the fluorine in para-CFT (**1**) remains in a more stable orientation, “sensing” a homogenous chemical environment (**Fig. 6b, movies 8, 11, 12**). Since the W70 side chain is in direct contact with ortho-CFT (**3**), the distinct dynamic behavior of this inhibitor may also explain the broad ^1^H, ^15^N-NMR signal of the W70 side chain observed in [Tpx WT/ortho-CFT]_2_ dimers (**Fig. 3b**).

To elucidate the contribution of dimer formation on the ^19^F-spectra, we diluted the Tpx WT and W39A protein inhibitor complexes from 750 µM to 50 µM (**Fig. 6c, Supplementary Fig. 18**). As expected, the mutant consistently showed narrower linewidths than the Tpx WT complexes throughout the entire dilution series. For all inhibitors bound to WT protein, the extent of changes in ^19^F-linewidths and chemical shifts directly reflected the respective dimer affinities, i.e. remaining nearly constant for the stronger dimerizers (**1-3, 5**), and approximating the values of the monomeric Tpx W39A complexes, indicative of dissociation, for the weakest dimerizer CtFT (**4**) (**Fig. 6c, green dots**). Of note, the ^19^F-NMR spectra also showed that at very high protein concentrations (∼750 µM), the W39A mutation showed residual dimer formation as apparent from linewidths and chemical shifts, an effect most pronounced for para-CFT (**1**) (**Fig. 6c, purple triangles, Supplementary Fig 18**). However, importantly, all our previous ^1^H, ^15^N-HSQC spectra of the Tpx W39A mutant were recorded at 100 µM, well below the dimerization threshold of this mutant (e.g. **Fig. 4**).

Our ¹⁹F-NMR data reflect the previously determined dimer affinities and show that small changes in inhibitor fluorination influence the conformational dynamics of the bound molecules. For instance, CtFT (**4**) adopts a stable, but dimer-incompetent pose with its CF_3_ group in a less deeply buried, chemically more homogenous binding position than the other molecules. Consistent with the intermediate dimer affinity seen by SEC, SEC-MALS and ITC, as well as ^1^H, ^15^N-NMR measurements and MD simulations, ortho-CFT (**3**) showed W39-dependent conformational dynamics that disrupted stable binding, thereby resulting in a heterogenous chemical environment and pronounced line broadening.

## Discussion

Chemically induced dimerization is a powerful tool in chemical biology and cell biology, that has paved the way for numerous innovative therapeutic applications.^1–9^ So far, a few isolated observations suggest that fluorination can affect protein heterodimerization, for instance in the case of quinabactin, a molecular glue that increases drought resistance in plants.^31^ However, although fluorine walks are widely used to modify the physicochemical properties of small molecules and have yielded numerous potent pharmaceuticals and agrochemicals from non-fluorinated precursor compounds in the past,^26–31^ the role of molecular glue fluorination for chemically induced dimerization has not yet been systematically explored.

Here, we built on our prior, serendipitous finding that the covalent modification of the essential parasitic oxidoreductase Tryparedoxin (Tpx) from *T. brucei* with a fluorinated inhibitor identified in a high-throughput screening induces enzyme homodimerization.^21,23^ Protein/protein contacts in the form of two salt bridges, as well as hydrophobic interactions between the aromatic side chains of W39 residues from both protomers stabilize the dimer as seen in the crystal structure of the Tpx/para-CFT complex.^23^

Using a fluorine walk library of covalent thienopyrimidinone inhibitors based on the originally identified Tpx dimerizer para-CFT (**1**), we demonstrate that fluorination is a straightforward method for tuning chemically induced protein dimerization (CID). The significant differences in dimerization capability we observed prompted a comprehensive investigation using a combination of biophysical methods. This approach was initially motivated by our inability to crystallize most protein/inhibitor complexes, but this limitation proved fortuitous, as the ability of a compound to act as a molecular glue for Tpx critically depends on its fluorination pattern and the resulting dynamic interplay between the bound inhibitor and a surface-exposed aromatic residue in solution. In our CID system, two dimerizer molecules form an integral part of the dimer interface, explaining why inhibitor orientation and dynamics are crucial parameters for chemically induced Tpx dimer stability.

In many enzymes, gatekeeping aromatic residues are crucial regulators of the catalytic process.^52^ In the case of Tpx, mutation of W39 to alanine results in a catalytically dead enzyme^23^, underscoring this residue’s pivotal function for the enzymatic process. However, to the best of our knowledge, this is the first time that the role of a gatekeeping aromatic residue has been discussed in the context of chemically induced dimerization. The W39 side chain can adopt many different orientations that are compatible with inhibitor-induced protein dimerization. Its dynamics are affected by inhibitor binding and at the same time, it engages in a “molecular quarrel” with inhibitors competing for the same binding site, thereby increasing their dynamics. Inhibitor fluorination critically affects whether and how efficiently an inhibitor can adhere to the binding pocket (**Fig. 7**). In contrast, W70 presents a “wing gate”^21^, i.e. a residue that is important for restricting ligand access to the binding pocket. In Tpx, its orientation and dynamics are affected by inhibitor interactions (**Fig. 3b, Supplementary Fig. S15c**), but it is not the primary modulator of inhibitor/surface interactions.

**Fig. 7:**
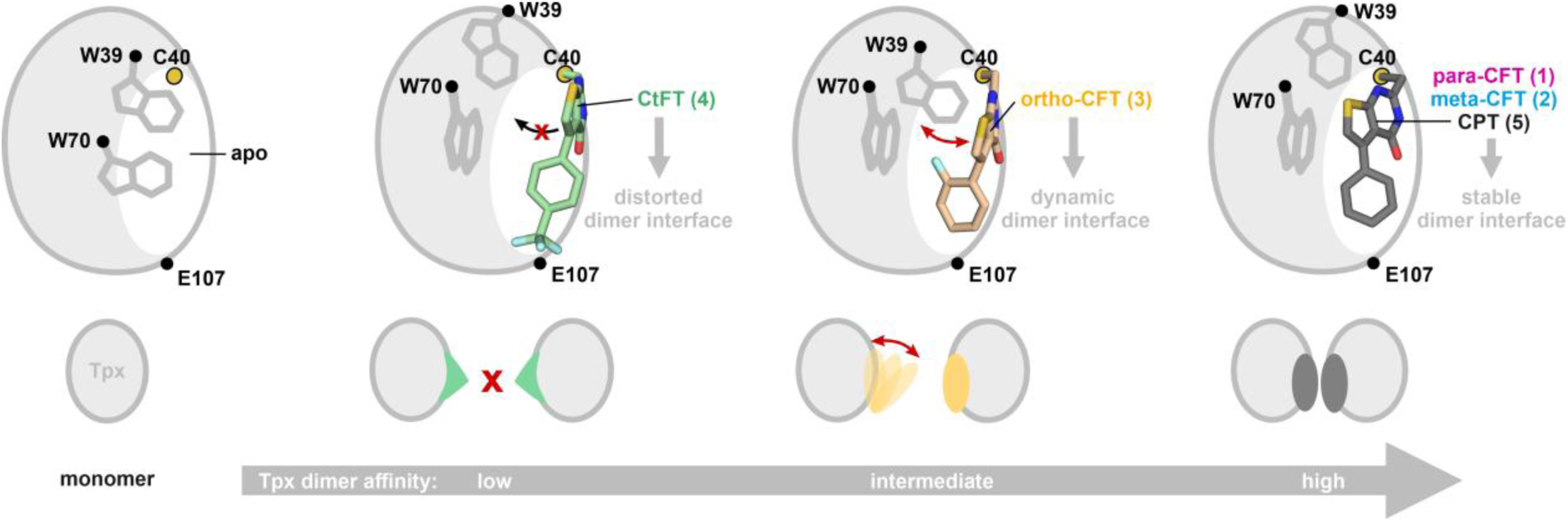
Consequences of the inhibitor fluorination pattern for Tpx adsorption and dimerization. Tryptophan side chain and inhibitor orientations were taken from representative MD-simulation frames (**Supplementary Fig. 17**). Both the dynamics of surface aromatic residues (W39, W70) and the molecular glue modulate protein dimer affinity. All inhibitors occupy a consensus binding pocket (Fig. 4a) and compete with the W39 sidechain for this site. Trifluoromethylation (**4**) results in a dimer-unproductive inhibitor stance because this molecule cannot adsorb tightly to the protein surface regardless of the orientation of W39. Bound ortho-CFT (**3**) can engage with the canonical binding pocket, but displays distinct dynamics due to a structural clash with W39 that results in a reduced dimer affinity compared to strong dimerizers (**1, 2** and **5**) which tightly adsorb to the induced Tpx binding pocket.

The very good agreement between dimer affinities observed in vitro and the occurrence of dimer disruptive inhibitor stances in MD simulations (**Fig. 3c-d, 5f**) underscores the value of an integrated approach including computational methods in understanding the underlying structure activity relationships (SARs) of molecular glues for potential future designs. MD simulations also provided crucial insights into the possible consequences of fluorophenyl conformations (“F-in/F-out”) for meta-and ortho-fluorinated dimerizers (**2, 3**), that were otherwise inaccessible through wet lab experimental techniques. While quantification of possible atropisomers and interconversion rates remain elusive with the current set of methods, it is tempting to speculate that even better dimerizers could be designed by e.g. restraining the fluorophenyl ring rotation prior to protein binding or “symmetrize” meta-and ortho-CFT (**2, 3**) by introducing a second fluorine substituent.

Although compounds **1**-**5** show limited selectivity for trypanosomes over HEK293 cells and are less potent than conventional non-covalent trypanocides such as suramin^53^, they still exhibit antiparasitic activity and act as covalent Tpx inhibitors that are highly selective for residue C40 in our in vitro activity assay (**Fig. 1, Supplementary Fig. 2**). Since Tpx is an essential enzyme that is highly conserved across human-and animal-pathogenic trypanosomatids^18,19^, and C40 is its active site nucleophile that cannot be mutated without loss of enzyme activity^23^, a potential application of the present compound series for drug development should not be dismissed. Their size and physicochemical properties position inhibitors **1-5** as promising fragments for drug design that fulfil the “Rule of 3”, which requires having desirable initial chemical properties, i.e. molecular weight, lipophilicity (logP) and number of hydrogen bond donors and acceptors.^54–56^

Nonetheless, and regardless of a future application in drug development, the molecular glues presented here constitute the first set of monovalent protein homodimerizers and therefore open new avenues as chemical biology tools. Current CID systems such as rapamycin-inspired molecules mediate ultra-high affinity, quasi irreversible protein/protein interactions.^6,57^ Here, a reversibly self-assembling CID system, covering high, intermediate and low micromolar protein affinities, may present a versatile alternative. Furthermore, covalent attachment of the ligand obviates concerns regarding dimerizer off-rates.

Our study demonstrates that shallow binding pockets on protein surfaces can present effective dimerization sites. Local side chain dynamics and fluorination can be exploited to modulate chemically induced dimerization efficiency. The fluorine walk is a powerful, yet hitherto underexplored approach for tuning the properties of molecular glues. In summary, we provide a molecular framework for understanding how inhibitor dynamics influence dimerization, offering a blueprint to guide future optimization of CID systems for therapeutic and research applications.

## Materials and Methods

### Materials

*T. brucei brucei* 449 Lister 427 parasites and HEK293 cells were generous gifts from Prof. Luise Krauth-Siegel, Heidelberg, and Prof. Thorsten Heinzel, Jena. Ni-NTA beads were purchased from Qiagen. Deuterium oxide (D_2_O) and ^15^N-ammonium chloride (^15^NH_4_Cl) were purchased from Eurisotop. U-^13^C-D-glucose was purchased from Cambridge Isotope Laboratories. NADPH was purchased from Roche and the crosslinker 1,8-bismaleimidodiethyleneglycol (BM(PEG)_2_), L-glutamine, trypsin, and penicillin-streptomycin were purchased from Thermo Fisher Scientific. DMEM and FBS for HEK293 cell culture were purchased from Sigma Aldrich. All other materials were obtained from Carl Roth GmbH unless specified otherwise and were of analytical grade or better.

### Molecular cloning of the parasitic oxidoreductase Tryparedoxin (Tpx)

The plasmid encoding Tpx wildtype (WT) is based on a pETtrx_1b vector and was a generous gift from Prof. Luise Krauth-Siegel, Heidelberg. Expression vectors for the point mutants Tpx W39A, Tpx C40S and Tpx C43S were obtained via Quick change PCR with the KAPA HiFi PCR Kit (Kapa Biosystems). The following primer pairs were used to obtain Tpx W39A (^5’^CTGCCTCCGCGTGCCCCCCATGC^3’^ and ^5’^GGGGGCACGCGGAGGCAGAAAAG-TAAAG^3’^), Tpx C40S (^5’^CTTTTCTGCCTCCTGGTCCCCCCCATGCC^3’^ and ^5’^GTAAAACCC-CGGCATGGGGGGGACCAGGAGGC^3’^); and Tpx C43S (^5’^CTGGTGCCCCCCATCCCGGGG-TTTTACACC^3’^ and ^5’^GGTGTAAAACCCCGGGATGGGGGGCACCAG^3’^).

### Expression and purification of *T. brucei* redox cascade components

Purified *T. brucei* trypanothione reductase (TR), trypanothione disulfide (TS_2_), and the plasmid encoding *T. brucei* glutathione peroxidase-type tryparedoxin peroxidase 3 (Px, pETtrx_1b vector) were generous gifts from Prof. Luise Krauth-Siegel, Heidelberg. TR was stored at a concentration of 1.3 mM at 4°C in 50 mM potassium phosphate buffer pH 7.0 containing 1 mM EDTA, and 1.8 M ammonium sulfate. TS_2_ was stored at a concentration of 22 mM in water at-20°C. *T. brucei* tryparedoxin 1 (Tpx) and glutathione peroxidase-type tryparedoxin peroxidase 3 (Px) were heterologously overexpressed as His_6_-tagged thioredoxin (Trx) fusion proteins in *E. coli* BL21-Gold(DE3) cells (Agilent) using either 2xYT medium or M9 minimal medium supplemented with ^15^N-NH_4_Cl (1 g/L) and ^13^C-D-glucose (2 g/L) as the sole nitrogen or carbon sources.^58^ Cells were lysed and proteins purified as described before:^46^ In brief, the Trx-His_6_-Tpx (or Trx-His_6_-Px) fusion protein was separated from the lysate via Ni-NTA affinity chromatography and cleaved with His-tagged *tobacco edge virus* (TEV) protease. After dialysis against Tpx buffer (25 mM sodium phosphate buffer pH 7.5, 150 mM NaCl) at 4 °C for 15 h, Tpx (or Px) was separated from the His-tagged Trx cleavage product and the TEV protease via Ni-NTA affinity chromatography, and purified by size exclusion chromatography using a HiLoad 16/600 Superdex 75 pg column (Cytiva). Proteins were eluted in Tpx buffer at 4 °C and purity of selected fractions were checked via SDS-PAGE (15% gel) before pooling. Purified proteins were stored at-80 °C.

### Synthesis and analysis of Tpx inhibitors

For details on synthesis and analysis, please see Supporting Information. All chemicals were purchased from commercial sources and used without prior purification. Air and humidity sensitive reactions were carried out under argon using the Schlenk technique. Deuterated chloroform was stored over basic aluminum oxide. Solvents for HPLC-MS were purchased from Fisher Scientific in Optima-HPLC-MS^®^ grade. Flash column chromatography was performed using silica gel with an average grain size of 35–70 µm and purchased from Acros Organics. Automated flash column chromatography was carried out using an Isolera Flash Purification System from Biotage with integrated UV-detector. Thin layer chromatography was performed using silica coated aluminum plates from Merck (TLC Silica 60 F_254_).

NMR measurements of small molecules in deuterated solvents purchased from Cambridge Isotope Laboratories were carried out using Avance III HD 300 and Avance III HD 400 spectrometers. Chemical shifts (δ) are reported in parts-per-million (ppm) relative to the residual solvent protons (^1^H-NMR) or the deuterium coupled ^13^C solvent signal (^13^C-NMR). Coupling constants of protons (J) are reported in Hertz (Hz). Spin multiplicities are reported using the following abbreviations or appropriate combinations of such: s (singlet), d (doublet), t (triplet), m (multiplet). NMR spectra were processed using MestReNova x64 14.01. (Mestrelab Research S.L.). Electron spray ionization high resolution mass spectrometry (ESI-HRMS) analysis was performed using a 6545 QTof Ultima 3 mass spectrometer with LockSpray-interface at capillary voltages of 2900 V. Electron spray ionization mass spectrometry (ESI) were recorded using a 1200-Series or a 1260-Series Infinity II HPLC-system from Agilent, equipped with a binary pump and integrated diode array detector. The latter was coupled to a LC/MSD-Trap-XTC-mass spectrometer or a LC/MSD Infinitylab LC/MSD (G6125B LC/MSD) from Agilent. An *Ascentis Express* C_18_-column (30 mm x 2.1 mm) with an average particle size of 2.7 μm was used as stationary phase. Fourier-transform infrared (FT-IR) spectra were recorded on a Bruker FT-IR Tensor 27 apparatus and reported in wavenumbers (υ). Melting temperatures were measured using a KSP1N from A. Krüss Optronic.

### Preparation of Tpx/Inhibitor complexes and crosslinked Tpx dimers

100-400 µM purified Tpx in 25 mM sodium phosphate buffer pH 7.5, 150 mM NaCl (Tpx buffer) were incubated with 4 mM tris(2-carboxyethyl)phosphine (TCEP) and a 3-fold excess of inhibitor (4 mM in DMSO) or a 4-fold excess of the crosslinker 1,8-bismaleimidodiethyleneglycol (BM(PEG)_2_, 10 mM in DMSO) for 30 min at 25 °C. Covalently modified Tpx was purified via size exclusion chromatography with Tpx buffer using a Superdex 75 Increase 10/300 GL column (Cytiva) at 4 °C.

### Intact-protein Mass Spectrometry (MS)

20 µM of Tpx WT, C40S and C43S in Tpx buffer were incubated with 400 µM TCEP and 2% DMSO containing no or 80 µM of inhibitors **1**-**5** for 30 min at 25°C. Crosslinked Tpx dimers were generated and purified as described above.

High-resolution intact-protein mass spectrometry was performed on a Dionex Ultimate 3000 HPLC system (Thermo Fisher) coupled to a Bruker maxis HD (after collision cell upgrade) with an electrospray ionization source (end plate offset: 500 V, capillary: 4500 V, nebulizer: 3.0 bar, dry gas: 12.0 L/min, dry temp: 250 °C). Samples were desalted on-line using a Massprep desalting cartridge (Waters) prior to MS-measurement. Internal calibration using sodium formate was performed at the end of each run. Deconvolution of protein spectra of individual peaks was performed using UniDec GUI 6.0.2.^59^

### Trypanosome and HEK293 cell survival assays

Half maximal effective concentration (EC_50_) values of compounds **1**-**5** against *T. brucei brucei* 449 Lister 427 parasites were determined with an ATPlite Luminescence assay as described previously.^21,23^ The EC_50_ was defined as the compound concentration at which a fifty percent decrease in relative luminescence occurred, which indicated that half of the trypanosomes had died. Toxicity of compounds **1-5** in human embryonic kidney (HEK293) was determined with a resazurin assay, which is based on the reduction of resazurin to fluorescent resorufin by living cells.^60^ HEK293 cells were maintained in DMEM high glucose supplemented with 10% FBS, 1% penicillin/streptomycin and 1% L-glutamine at 37 °C and 5% CO_2_. For the toxicity screen, 40000 cells per well were seeded in a 48 well plate (400 µL medium/well) and incubated at 37 °C for 24 hours before treatment with inhibitors **1-5**. Cells were treated with 0.1 µM, 0.3 µM, 0.7 µM, 1 µM, 3 µM, 7 µM, 10 µM, 30 µM, 70 µM and 100 µM of the respective inhibitor, DMSO treatment served as negative control. The overall DMSO concentration was kept at 1 vol.% DMSO per well. After 24 hours of treatment, the medium was exchanged with a medium/resazurin mixture (0.015 mg/mL resazurin per well) and the cells were incubated for 3 h at 37 °C before transferring 100 µl medium from each well into a black opaque-walled 96-well plate. Fluorescence was directly measured using a CLARIOstar Plus reader (BMG Labtech GmbH, Ortenberg, Germany) with a 545-20 nm excitation / 600-40 nm emission filter. Each experiment was done in biological quadruplets and cell viability was analyzed in comparison to the DMSO control by plotting relative fluorescence units (RFU) against inhibitor concentration (Prism, Graphpad). A fifty percent decrease in relative fluorescence indicated the compound concentration where half of the HEK293 cells had died, which was defined as the EC_50_.

### In vitro Tpx activity assay and inactivation kinetics

To reconstitute the peroxide clearance cascade (PCC), its components (at final concentrations of 360 µM NADPH, 0.1 U/mL TR, 110 µM TS_2_, 5 µM Tpx, 180 nM Px, 100 µM H_2_O_2_) were mixed in a total volume of 200 µL in a 10 mm quartz cuvette. First, 15 µL NADPH (4.8 mM in TR buffer (40 mM HEPES pH 7.5, 1 mM Na-EDTA), 10 µL TR (2 U/mL in TR buffer), 10 µL TS_2_ (2.2 mM in sterile filtered, deionized water), 20 µL Tpx (50 µM in Tpx buffer), 10 µL Px (3.6 µM in Tpx buffer), and 2 µL DMSO were added to 128 µL of Px buffer (100 mM Tris pH 7.5, 150 mM NaCl). The reaction mixture was incubated at 25 °C and absorption at 340 nm was observed with a UVmc2 double-beam UV-Visible spectrophotometer (SAFAS) until the baseline stabilized before starting the reaction with 5 µL H_2_O_2_ (from a 4 mM stock in sterile filtered, deionized water). The absorption at 340 nm was measured for 30 s and fitted linearly (SAFAS SP2000). Of note, this observed PCC activity includes both the enzymatic and non-enzymatic oxidation of NADPH by atmospheric oxygen.

To determine the contribution of the non-enzymatic NADPH oxidation, the reaction mixture was also prepared without Px and measured as described above. Subtracting this value from the observed PCC activity yielded the actual PCC activity, which was set to 100% activity (0% inhibition) for further analyses.

Measuring PCC inhibition (I) at defined time points (t) after the addition of DMSO-dissolved inhibitors **1**-**5** yielded time-resolved inhibition curves. Substoichiometric (2.5 µM, stoichiometry relative to Tpx), equimolar (5 µM) and excess amounts of inhibitor (15 µM, 20 µM, 25 µM, 30 µM for ortho-CFT and CtFT; 15 µM, 25 µM, 35 µM, 45 µM for para-CFT, meta-CFT and CPT) were added to the PCC mixture prior to starting the reaction with H_2_O_2_. For equimolar inhibitor amounts, measurements were carried out as technical triplicates of biological triplicates, in all other cases as technical triplicates. Fitting inhibition (I) at defined time points (t) with **Eq. 1**:

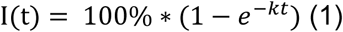

for a given inhibitor concentration yielded k_obs_, which was plotted linearly against inhibitor concentration (shown as X-fold excess relative to Tpx on plots) to calculate second-order rate constants k_2nd_ for compounds **1-5** (OriginPro 2022).^44^

### In vitro inhibitor selectivity assays

To determine the selectivity of the inhibitors in the in vitro PCC reaction mix for each enzyme, we modified the PCC assay to leave out individual components. To determine TR inhibition by compounds **1**-**5**, the PCC assay was carried out using only 150 µM NADPH, 10 mU/mL TR, and 100 µM TS_2_. For the basal TR activity, we added 6.3 µL NADPH (4.8 mM in TR buffer), 10 µL TR (0.2 U/mL in TR buffer), and 2 µL DMSO (or 12 µL to emulate the CtFT conditions, see below) to 172.6 µL TR buffer (or 162.6 µL for the CtFT condition) before starting the reaction with 9.1 µL TS_2_ (2.2 mM in sterile filtered, deionized water). The absorption at 340 nm was measured for 30 s at 25 °C and fitted with the Safas SP2000 software.

To determine TR inhibition, a final concentration of 4.5 µM inhibitor (3 µM for ortho-CFT and CtFT) in 2 µL DMSO (12 µL for CtFT) was added to the reaction mixture and incubated for 5 min at 25 °C before starting the reaction with TS_2_. TR inhibition was determined in a technical triplicate as described above.

To check Tpx activity, we incubated the PCC in the absence of Px (360 µM NADPH, 0.1 U/mL TR, 110 µM TS_2_, 5 µM Tpx) with 5 µM of each inhibitor **1**-**5** or DMSO for 25 min (45 min for meta-CFT and CPT) at 25 °C before adding 190 nM Px and immediately starting the reaction with 100 µM H_2_O_2_.^21^ PCC activity was determined from the absorption decrease at 340 nm as described above in a technical triplicate. The ratio of PCC activity in the presence of inhibitor and the corresponding DMSO control yielded the PCC inhibition due to Tpx binding of inhibitors **1-5**.

To check Px activity, we reduced 25 µM Px with 4 mM TCEP in 100 µL Tpx buffer for 30 min at 25 °C before adding 35 µM inhibitor (500 µM in 7 µL DMSO) or pure DMSO and incubating for another 25 min (45 min for meta-CFT and CPT) at 25 °C. The reaction mixture was washed three times with 4 mL Tpx buffer using centrifugal filters with a cut-off of 10 kDa (Merck) to remove TCEP, DMSO, and unbound inhibitor. Px was diluted to 3.6 µM and used in the PCC assay as described above in a technical triplicate.

### Analytical size exclusion chromatography (SEC)

100 µM (or 400 µM) purified Tpx with 4 mM TCEP in 200 µL Tpx buffer were incubated with 2.5 µL, 5 µL, and 15 µL (or 60 µL when 400 µM Tpx were used) inhibitor from a 4 mM stock in DMSO or 15 µL (60 µL when 400 µM Tpx were used) pure DMSO for 30 min at 25 °C. The reaction mixtures were applied to a Superdex 75 Increase 10/300 GL column (Cytiva) and eluted with Tpx buffer at 4 °C, while recording the absorption at 280 nm. Elution volumes, defined as the peak maxima, were determined with the Chromlab Software (version 6.1, Biorad) in a biological triplicate. 50 µM crosslinked Tpx dimer were eluted as a control.

SEC analyses of 100 µM Tpx WT and Tpx/inhibitor complexes with para-CFT (**1**) and CtFT (**4**), as well as 50 µM crosslinked Tpx_2_, were repeated in the presence of Tpx-buffer with 10% and 30% isopropanol to investigate secondary interactions of Tpx constructs with the SEC column.^61^

### Circular dichroism (CD) spectroscopy

To assess the structural integrity of Tpx with isopropanol, CD spectroscopy was used. Purified Tpx (10 µM) in 2.5 mM NaP_i_ pH 7.5, 15 mM NaCl was incubated for 30 min at RT in the presence or absence of 30% isopropanol before recording CD spectra (195-260 nm) in a 1 mm quartz cuvette (Hellma) with a J-1500 Circular Dichroism Spectrophotometer (Jasco). As blanks, buffers with no or 30% isopropanol were used, respectively. The ellipticity *θ* (in degrees) was measured in a technical triplicate by subtracting the buffer spectra (mean of three measurements) from the protein spectra. The mean residue ellipticity ([*θ*]) was calculated with **Eq. 2**:

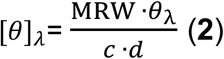

where *θ*_λ_ is the measured ellipticity (in degrees) at a given wavelength λ (in nm), *c* the protein concentration (in g/mL), and *d* the pathlength (in cm). The MRW is the mean residue weight and was calculated with **Eq. 3**:

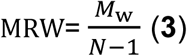

where *M*_w_ is the protein molecular weight (in g/mol) and *N* the total number of protein residues.

### Size exclusion chromatography coupled to multi-angle light scattering (SEC-MALS)

To determine the oligomeric state of Tpx/inhibitor complexes by SEC-MALS, 100 µM of purified Tpx/inhibitor complexes were applied to a SEC column (Superdex75 10/300, GE Healthcare) in Tpx buffer (25 mM NaP_i_ pH 7.5, 150 mM NaCl) at a flow rate of 0.5 ml/min at room temperature. The molecular weights of the eluting proteins were measured using a Dawn Heleos 8+ light scattering detector and an Optilab T-rEX refractive index detector (Wyatt Technologies). Data were analyzed with the ASTRA software (version 6.1.5.22, Wyatt Technologies), assuming a Zimm model.^62^

### Isothermal titration calorimetry (ITC)

Dissociation coefficients (*K*_D_) of dimeric Tpx/inhibitor complexes were determined via dilution ITC following our previously established protocol.^23^ Purified complexes in Tpx buffer were titrated to Tpx buffer, and the heat signals of each injection was recorded. To observe dimer dissociation, measurements were conducted with 78 µM (Tpx/para-CFT), 122 µM (Tpx/meta-CFT), 277 µM (Tpx/ortho-CFT), 821 µM (Tpx/CtFT), and 176 µM (Tpx/CPT). A MicroCal PEAQ-ITC Automated (Malvern Panalytical) was used to determine the *K*_D_ of chemically induced Tpx dimers with compounds **1, 3** and **5**. After the initial wait time of 2 min, 19 serial injections of 2 μL with a spacing of 3 min where performed. The reference power was set to 11 μcal sec^-1^ and stirring speed to 750 rpm. A MicroCal PEAQ-ITC (Malvern Panalytical) with the same experimental setup was used to determine the *K*_D_ value of covalent Tpx WT complexes with meta-CFT (**2**, 10 injections), and CtFT (**4**, 20 injections). Each titration was performed in a technical triplicate, and controls included water to water and unmodified Tpx to buffer titrations. Transferability between instruments was checked by repeating the Tpx/para-CFT titration. All thermograms were analyzed with the MicroCal PEAQ-ITC analysis software (Malvern Panalytical).

### ^19^F-nuclear magnetic resonance (^19^F-NMR) spectroscopy

Tpx/inhibitor complexes with fluorinated compounds **1-4** were prepared and purified as described above. Samples were supplemented with 10% D_2_O for field-frequency locking. Spectra of 750 µM, 500 µM, 250 µM, 100 µM, and 50 µM inhibitor-bound Tpx WT or Tpx W39A complexes were recorded on a Bruker AVANCE III HD 600 MHz spectrometer equipped with a QCI cryoprobe (Bruker, Rheinstetten) with 512 scans (4096 scans for Tpx WT/ortho-CFT complexes due to line broadening) at 298 K. Spectra were processed using TopSpin 4.1 and referenced to trifluoroacetic acid (TFA, signal set to-76.55 ppm).

For free inhibitors, 100 µM of compounds **1-4** were dissolved in Tpx buffer with 10% D_2_O and 8% DMSO (13% for CtFT). ^19^F-NMR spectra were measured and processed as described above. The effect of DMSO on the fluorine signal was determined via TFA referencing and corrected (-0.16 ppm for 8% DMSO,-0.27 ppm for 13% DMSO).

### ^1^H, ^15^N-NMR chemical shift perturbation measurements

^15^N-labelled Tpx WT and Tpx W39A, as well as Tpx/inhibitor complexes with compounds **1**-**5** were prepared as described above and supplemented with 10% D_2_O for field-frequency locking, and 0.1 mM 2,2-dimethyl-2-silapentane-5-sulfonic acid (DSS) for hydrogen referencing (DSS signal set to 0 ppm). HSQC spectra were recorded for Tpx WT (150 µM + 4 mM TCEP), Tpx WT/inhibitor complexes (120-180 µM), Tpx W39A (100 µM + 4 mM TCEP) and Tpx W39A/inhibitor complexes (100 µM) at 298 K with a Bruker Avance Neo 800 MHz spectrometer equipped with a TCI cryoprobe (Bruker, Rheinstetten), and processed using TopSpin 4.1. Data analysis was carried out in CcpNMR analysis 3.1.^63^ and the inhibitor induced ^1^H, ^15^N-chemical shift differences **Δ**δ were calculated using **Eq. 4**^64^:

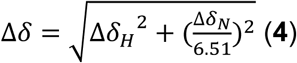

where Δδ_H_ and Δδ_N_ are the ^1^H-and ^15^N-chemical shift difference, respectively.

A set of triple resonance assignment experiments (HNCA, HN(CO)CA) implemented in the standard Bruker TopSpin pulse sequences library were recorded for ^15^N,^13^C-labelled Tpx WT (300 µM + 4 mM TCEP), Tpx WT/para-CFT (180 µM), Tpx WT/CtFT (180 µM), Tpx WT/ortho-CFT (350 µM), Tpx W39A (600 µM + 4 mM TCEP) and Tpx W39A/para-CFT (200 µM) at 298 K, processed and analyzed as described above to obtain the backbone NMR assignments of the complexes.

### Density functional theory (DFT) calculations

The molecular structures of compounds **1-5** were constructed using ChemBioDraw 22 and initially optimized with the MM2 (Merck Molecular Force Field) method in ChemBioDraw 3D. Further optimization was performed using density functional theory (DFT) at the B3LYP/6-31G(d) level^65^, incorporating Grimme’s DFT-D3 dispersion correction.^66^ To explore conformational flexibility, potential energy surface (PES) scans were conducted by systematically varying the dihedral angle between the thienopyrimidinone core and the phenyl ring of the Tpx inhibitor in 10-degree increments. These calculations were carried out using the M06-2X/def2-TZVP level of theory within the SMD solvation model for water^67–69^, also accounting for Grimme’s DFT-D3 dispersion correction. All calculations were performed using the Gaussian 16 program package.^70^

### Molecular dynamics (MD) simulations

MD simulations of Tpx/inhibitor complexes (Tpx WT dimer, Tpx WT monomer, and Tpx W39A monomer) were set up using our Tpx/para-CFT dimer crystal structure (PDB: 6GXG^23^, chain A). For meta-and ortho-CFT (**2**-**3**), two fluorine orientations, either pointing towards the protein binding pocket or the solvent (“F-in” and “F-out”, respectively) were considered. The PyMOL Molecular Graphics System (Version 3.0, Schrödinger, Inc.) was used to prepare and edit structures, and acpype^71^ was used to parameterize the cysteine bound inhibitors as dipeptides, adding the parametrization for the respective inhibitor to the amberff14SB^72^ gromacs port as residue “CYP”. The systems were prepared for MD-simulations by placing the Tpx/inhibitor complexes in a box with 1 nm padding, solvated with TIP3P water^73^ and neutralized with sodium ions.

MD simulations were carried out using GROMACS 2021.2.^74–78^ Energy minimization was performed for a maximum of 1000 steps until the maximum force was below 1000 kJ/mol/nm with positional restraints on the protein, then for a maximum of 2500 steps without restraints. Subsequently, *nVT-* and *npT-*equilibrations for 100 ps followed, setting a temperature of 300 K with the *V-rescale* modified Berendsen thermostat^79^ using a time constant of 0.1 ps and a pressure of 1 bar with the Parrinello-Rahman barostat^80^ using a time constant of 2 ps. Hydrogen bonds were constrained using the LINCS algorithm.^81^ Simulations ran for 100 ns (5x 10^7^ steps of 2 fs). All MD simulations were evaluated using GROMACS (modules *gmx angle*, *gmx pairdist*, and *gmx rms*), Python 3^82^ scripts, and *jupyter* notebooks of the Anaconda distribution^83^ (modules *matplotlib 3.8.0*^84^ and *numpy 1.20.3*^85^). In our analysis of MD simulations, we included all replicates (n = 5 for each set of conditions) and, in the case of dimer simulations, both protein chains. For Tpx complexes with asymmetrically substituted compounds **2** and **3**, we distinguished between “F-in” and “F-out” conformations. Hence, for the analysis of corresponding Tpx dimers with compounds **2** and **3**, we evaluated three different MD simulations with either both inhibitors in the same orientation (both chains “F-in” or both “F-out”), or with mixed conformations (one chain in “F-in”, and one chain in “F-out” conformation), and assumed statistical distribution (25% “F-in/F-in”, 25% “F-out/F-out”, and 50% “F-in/F-out”) (see Extended Material and Methods for details).

## Data availability statement

The crystal structures of isolated compounds **1-5** can be found under the CCDC deposition numbers 1862408, 2421180, 2421181, 2421182, and 2423222. The crystal structure of the Tpx/para-CFT dimer can be found under the PDB accession number 6GXG.^23^ The backbone NMR assignments of Tpx WT in the reduced and oxidized state can be obtained from BMRB accession numbers 27049 and 27050^46^, and the assignment of Tpx W39A in the reduced state under the BMRB accession number 53259.

## Acknowledgements

We gratefully acknowledge Prof. Luise Krauth Siegel for continuous support, Dr. Philipp Klein, Mainz, for the synthesis of para-CFT, Prof. Jens Wöhnert, Frankfurt, and Prof. Tanja Schirmeister, Mainz, for access to ITC instruments, Dr. Dieter Schollmeyer, Mainz, for small molecule crystal structure analysis, and Dr. Christoph Wiedemann, Jena, for technical support and fruitful discussions. This study made use of NMRbox: National Center for Biomolecular NMR Data Processing and Analysis, a Biomedical Technology Research Resource (BTRR), which is supported by NIH grant P41GM111135 (NIGMS). The authors acknowledge the computational resources (HPC-cluster “Draco”) provided by the Computing Center of the Friedrich-Schiller-University Jena. Financial support by the Deutsche Forschungsgemeinschaft (DFG, German Research Foundation) within the framework of the graduate research center “Life Sciences – Life Writing” (GRK2015, project number 244248598) through a PhD fellowship awarded to E.S., an individual research grant (Project ID 438511573 to UAH) and the Cluster of Excellence “Balance of the Microverse” EXC2051— Project-ID 390713860 (to P.S., M.L. and U.A.H.). M.L. is grateful for support by the research profile line LIFE of FSU Jena and the Deutsche Forschungsgemeinschaft (DFG, German Research Foundation) via the Emmy-Noether-Program (project ID 528114058). U.A.H. acknowledges an instrumentation grant for a high-field NMR spectrometer by the REACT-EU EFRE Thuringia (Recovery assistance for cohesion and the territories of Europe, European Fonds for Regional Development, Thuringia) initiative of the European Union.

## References

1. Stanton, B. Z., Chory, E. J. & Crabtree, G. R. Chemically induced proximity in biology and medicine. *Science (New York*, N.Y.) 359; 10.1126/science.aao5902 (2018).

2. Voß, S., Klewer, L. & Wu, Y.-W. Chemically induced dimerization: reversible and spatiotemporal control of protein function in cells. Current Opinion in Chemical Biology 28, 194–201; 10.1016/j.cbpa.2015.09.003 (2015).

3. Fegan, A., White, B., Carlson, J. C. T. & Wagner, C. R. Chemically controlled protein assembly: techniques and applications. Chemical Reviews 110, 3315–3336; 10.1021/cr8002888 (2010).

4. Ross, B., Mehta, S. & Zhang, J. Molecular tools for acute spatiotemporal manipulation of signal transduction. Current Opinion in Chemical Biology 34, 135–142; 10.1016/j.cbpa.2016.08.012 (2016).

5. Schreiber, S. L. The Rise of Molecular Glues. Cell 184, 3–9; 10.1016/j.cell.2020.12.020 (2021).

6. Camacho-Soto, K., Castillo-Montoya, J., Tye, B., Ogunleye, L. O. & Ghosh, I. Small molecule gated split-tyrosine phosphatases and orthogonal split-tyrosine kinases. Journal of the American Chemical Society 136, 17078–17086; 10.1021/ja5080745 (2014).

7. Maniaci, C. & Ciulli, A. Bifunctional chemical probes inducing protein-protein interactions. Current Opinion in Chemical Biology 52, 145–156; 10.1016/j.cbpa.2019.07.003 (2019).

8. Békés, M., Langley, D. R. & Crews, C. M. PROTAC targeted protein degraders: the past is prologue. Nat Rev Drug Discov 21, 181–200; 10.1038/s41573-021-00371-6 (2022).

9. Zhao, L., Zhao, J., Zhong, K., Tong, A. & Da Jia. Targeted protein degradation: mechanisms, strategies and application. Sig Transduct Target Ther 7, 113; 10.1038/s41392-022-00966-4 (2022).

10. Dong, G., Ding, Y., He, S. & Sheng, C. Molecular Glues for Targeted Protein Degradation: From Serendipity to Rational Discovery. Journal of Medicinal Chemistry 64, 10606–10620; 10.1021/acs.jmedchem.1c00895 (2021).

11. Grohmann, C., Marapana, D. S. & Ebert, G. Targeted protein degradation at the host-pathogen interface. Molecular Microbiology 117, 670–681; 10.1111/mmi.14849 (2022).

12. Espinoza-Chávez, R. M. et al. Targeted Protein Degradation for Infectious Diseases: from Basic Biology to Drug Discovery. ACS bio & med chem Au 3, 32–45; 10.1021/acsbiomedchemau.2c00063 (2023).

13. Control of Neglected Tropical Diseases. Ending the neglect to attain the sustainable development goals: a framework for monitoring and evaluating progress of the road map for neglected tropical diseases 2021−2030. World Health Organization (2021).

14. Büscher, P., Cecchi, G., Jamonneau, V. & Priotto, G. Human African trypanosomiasis. The Lancet 390, 2397–2409; 10.1016/S0140-6736(17)31510-6 (2017).

15. Shaw, A. P. M., Cecchi, G., Wint, G. R. W., Mattioli, R. C. & Robinson, T. P. Mapping the economic benefits to livestock keepers from intervening against bovine trypanosomosis in Eastern Africa. Preventive Veterinary Medicine 113, 197–210; 10.1016/j.prevetmed.2013.10.024 (2014).

16. Giordani, F., Morrison, L. J., Rowan, T. G., Koning, H. P. de & Barrett, M. P. The animal trypanosomiases and their chemotherapy: a review. Parasitology 143, 1862–1889; 10.1017/S0031182016001268 (2016).

17. Swallow, B. M. Impacts of trypanosomiasis on african agriculture (Food and Agriculture Organization of the United Nations, Rome, 2000).

18. Krauth-Siegel, R. L. & Comini, M. A. Redox control in trypanosomatids, parasitic protozoa with trypanothione-based thiol metabolism. Biochimica et biophysica acta 1780, 1236–1248; 10.1016/j.bbagen.2008.03.006 (2008).

19. Gommel, D. U. et al. Catalytic characteristics of tryparedoxin. European Journal of Biochemistry 248, 913–918; 10.1111/j.1432-1033.1997.t01-1-00913.x (1997).

20. Comini, M. A., Krauth-Siegel, R. L. & Flohé, L. Depletion of the thioredoxin homologue tryparedoxin impairs antioxidative defence in African trypanosomes. Biochem J 402, 43– 49; 10.1042/BJ20061341 (2007).

21. Fueller, F., Jehle, B., Putzker, K., Lewis, J. D. & Krauth-Siegel, R. L. High throughput screening against the peroxidase cascade of African trypanosomes identifies antiparasitic compounds that inactivate tryparedoxin. The Journal of biological chemistry 287, 8792–8802; 10.1074/jbc.M111.338285 (2012).

22. Dietschreit, J. C. B. et al. Predicting 19 F NMR Chemical Shifts: A Combined Computational and Experimental Study of a Trypanosomal Oxidoreductase-Inhibitor Complex. Angewandte Chemie International Edition 59, 12669–12673; 10.1002/anie.202000539 (2020).

23. Wagner, A. et al. Inhibitor-Induced Dimerization of an Essential Oxidoreductase from African Trypanosomes. Angewandte Chemie International Edition 58, 3640–3644; 10.1002/anie.201810470 (2019).

24. Dewey, J. A. et al. Molecular Glue Discovery: Current and Future Approaches. Journal of Medicinal Chemistry 66, 9278–9296; 10.1021/acs.jmedchem.3c00449 (2023).

25. Lagardère, P., Fersing, C., Masurier, N. & Lisowski, V. Thienopyrimidine: A Promising Scaffold to Access Anti-Infective Agents. *Pharmaceuticals (Basel*, Switzerland) 15; 10.3390/ph15010035 (2021).

26. Berninger, M. et al. Fluorine walk: The impact of fluorine in quinolone amides on their activity against African sleeping sickness. European Journal of Medicinal Chemistry 152, 377–391; 10.1016/j.ejmech.2018.04.055 (2018).

27. Lasing, T. et al. Synthesis and antileishmanial activity of fluorinated rhodacyanine analogues: The’fluorine-walk’ analysis. Bioorganic & Medicinal Chemistry 28, 115187; 10.1016/j.bmc.2019.115187 (2020).

28. Lindsley, C. W. et al. Practical Strategies and Concepts in GPCR Allosteric Modulator Discovery: Recent Advances with Metabotropic Glutamate Receptors. Chemical Reviews 116, 6707–6741; 10.1021/acs.chemrev.5b00656 (2016).

29. Garai, S. et al. Application of Fluorine-and Nitrogen-Walk Approaches: Defining the Structural and Functional Diversity of 2-Phenylindole Class of Cannabinoid 1 Receptor Positive Allosteric Modulators. Journal of Medicinal Chemistry 63, 542–568; 10.1021/acs.jmedchem.9b01142 (2020).

30. Zhou, Y. et al. Next Generation of Fluorine-Containing Pharmaceuticals, Compounds Currently in Phase II-III Clinical Trials of Major Pharmaceutical Companies: New Structural Trends and Therapeutic Areas. Chemical Reviews 116, 422–518; 10.1021/acs.chemrev.5b00392 (2016).

31. Cao, M.-J. et al. Combining chemical and genetic approaches to increase drought resistance in plants. Nat Commun 8, 1183; 10.1038/s41467-017-01239-3 (2017).

32. Purser, S., Moore, P. R., Swallow, S. & Gouverneur, V. Fluorine in medicinal chemistry. Chem. Soc. Rev. 37, 320–330; 10.1039/B610213C (2008).

33. Müller, K., Faeh, C. & Diederich, F. Fluorine in pharmaceuticals: looking beyond intuition. *Science (New York*, N.Y.) 317, 1881–1886; 10.1126/science.1131943 (2007).

34. Smart, B. E. Fluorine substituent effects (on bioactivity). Journal of Fluorine Chemistry 109, 3–11; 10.1016/S0022-1139(01)00375-X (2001).

35. Gronenborn, A. M. Small, but powerful and attractive: 19F in biomolecular NMR. Structure 30, 6–14; 10.1016/j.str.2021.09.009 (2022).

36. Dolbier, W. R. (ed.). Guide to fluorine nuclear magnetic resonance for organic chemists. 2nd ed. (John Wiley & Sons, Hoboken, New Jersey, 2016).

37. Hellmich, U. A., Pfleger, N. & Glaubitz, C. F-MAS NMR on proteorhodopsin: enhanced protocol for site-specific labeling for general application to membrane proteins. Photochemistry and photobiology 85, 535–539; 10.1111/j.1751-1097.2008.00498.x (2009).

38. Maus, H. et al. The effects of allosteric and competitive inhibitors on ZIKV protease conformational dynamics explored through smFRET, nanoDSF, DSF, and 19F NMR. European Journal of Medicinal Chemistry 258, 115573; 10.1016/j.ejmech.2023.115573 (2023).

39. An Overview of Fluorine NMR. In Guide to fluorine nuclear magnetic resonance for organic chemists, edited by W. R. Dolbier. 2nd ed. (John Wiley & Sons, Hoboken, New Jersey, 2016), pp. 9–53.

40. Arntson, K. E. & Pomerantz, W. C. K. Protein-Observed Fluorine NMR: A Bioorthogonal Approach for Small Molecule Discovery. Journal of Medicinal Chemistry 59, 5158–5171; 10.1021/acs.jmedchem.5b01447 (2016).

41. Rose-Sperling, D., Tran, M. A., Lauth, L. M., Goretzki, B. & Hellmich, U. A. 19F NMR as a versatile tool to study membrane protein structure and dynamics. Biological Chemistry 400, 1277–1288; 10.1515/hsz-2018-0473 (2019).

42. Nogoceke, E., Gommel, D. U., Kiess, M., Kalisz, H. M. & Flohé, L. A unique cascade of oxidoreductases catalyses trypanothione-mediated peroxide metabolism in Crithidia fasciculata. Biological Chemistry 378, 827–836; 10.1515/bchm.1997.378.8.827 (1997).

43. Schlecker, T. et al. Substrate specificity, localization, and essential role of the glutathione peroxidase-type tryparedoxin peroxidases in Trypanosoma brucei. Journal of Biological Chemistry 280, 14385–14394; 10.1074/jbc.M413338200 (2005).

44. Strelow, J. M. A Perspective on the Kinetics of Covalent and Irreversible Inhibition. SLAS Discovery 22, 3–20; 10.1177/1087057116671509 (2017).

45. Williamson, M. P. Using chemical shift perturbation to characterise ligand binding. Progress in Nuclear Magnetic Resonance Spectroscopy 73, 1–16; 10.1016/j.pnmrs.2013.02.001 (2013).

46. Wagner, A., Diehl, E., Krauth-Siegel, R. L. & Hellmich, U. A. Backbone NMR assignments of tryparedoxin, the central protein in the hydroperoxide detoxification cascade of African trypanosomes, in the oxidized and reduced form. Biomol NMR Assign 11, 193–196; 10.1007/s12104-017-9746-7 (2017).

47. Alphey, M. S. et al. Tryparedoxins from Crithidia fasciculata and Trypanosoma brucei: photoreduction of the redox disulfide using synchrotron radiation and evidence for a conformational switch implicated in function. Journal of Biological Chemistry 278, 25919–25925; 10.1074/jbc.M301526200 (2003).

48. Jumper, J. et al. Highly accurate protein structure prediction with AlphaFold. Nature 596, 583–589; 10.1038/s41586-021-03819-2 (2021).

49. Varadi, M. et al. AlphaFold Protein Structure Database in 2024: providing structure coverage for over 214 million protein sequences. Nucleic Acids Res 52, D368–D375; 10.1093/nar/gkad1011 (2024).

50. Fiorillo, A., Colotti, G., Boffi, A., Baiocco, P. & Ilari, A. The crystal structures of the tryparedoxin-tryparedoxin peroxidase couple unveil the structural determinants of Leishmania detoxification pathway. PLoS neglected tropical diseases 6, e1781; 10.1371/journal.pntd.0001781 (2012).

51. Hofmann, B. et al. Structures of tryparedoxins revealing interaction with trypanothione. Biological Chemistry 382, 459–471; 10.1515/BC.2001.056 (2001).

52. Gora, A., Brezovsky, J. & Damborsky, J. Gates of enzymes. Chemical Reviews 113, 5871–5923; 10.1021/cr300384w (2013).

53. Kaminsky, R., Schmid, C. & Brun, R. An in vitro selectivity index for evaluation of cytotoxicity of antitrypanosomal compounds. In Vitro Toxicol. 9, 315–324 (1996).

54. Kirsch, P., Hartman, A. M., Hirsch, A. K. H. & Empting, M. Concepts and Core Principles of Fragment-Based Drug Design. *Molecules (Basel*, Switzerland) 24; 10.3390/molecules24234309 (2019).

55. Congreve, M., Carr, R., Murray, C. & Jhoti, H. A’rule of three’ for fragment-based lead discovery? Drug Discovery Today 8, 876–877; 10.1016/S1359-6446(03)02831-9 (2003).

56. Jhoti, H., Williams, G., Rees, D. C. & Murray, C. W. The’rule of three’ for fragment-based drug discovery: where are we now? Nat Rev Drug Discov 12, 644–645; 10.1038/nrd3926-c1 (2013).

57. Banaszynski, L. A., Liu, C. W. & Wandless, T. J. Characterization of the FKBP.rapamycin.FRB ternary complex. Journal of the American Chemical Society 127, 4715–4721; 10.1021/ja043277y (2005).

58. Marley, J., Lu, M. & Bracken, C. A method for efficient isotopic labeling of recombinant proteins. J Biomol NMR 20, 71–75; 10.1023/A:1011254402785 (2001).

59. Marty, M. T. et al. Bayesian deconvolution of mass and ion mobility spectra: from binary interactions to polydisperse ensembles. Analytical chemistry 87, 4370–4376; 10.1021/acs.analchem.5b00140 (2015).

60. O’Brien, J., Wilson, I., Orton, T. & Pognan, F. Investigation of the Alamar Blue (resazurin) fluorescent dye for the assessment of mammalian cell cytotoxicity. European Journal of Biochemistry 267, 5421–5426; 10.1046/j.1432-1327.2000.01606.x (2000).

61. Hellberg, U., Ivarsson, J.-P. & Johansson, B.-L. Characteristics of Superdex® prep grade media for gel filtration chromatography of proteins and peptides. Process Biochemistry 31, 163–172; 10.1016/0032-9592(95)00044-5 (1996).

62. Zimm, B. H. The Scattering of Light and the Radial Distribution Function of High Polymer Solutions. J. Chem. Phys. 16, 1093–1099; 10.1063/1.1746738 (1948).

63. Skinner, S. P. et al. CcpNmr AnalysisAssign: a flexible platform for integrated NMR analysis. J Biomol NMR 66, 111–124; 10.1007/s10858-016-0060-y (2016).

64. Mulder, F. A., Schipper, D., Bott, R. & Boelens, R. Altered flexibility in the substrate-binding site of related native and engineered high-alkaline Bacillus subtilisins. Journal of Molecular Biology 292, 111–123; 10.1006/jmbi.1999.3034 (1999).

65. Becke, A. D. Density-functional thermochemistry. III. The role of exact exchange. J. Chem. Phys. 98, 5648–5652; 10.1063/1.464913 (1993).

66. Grimme, S., Antony, J., Ehrlich, S. & Krieg, H. A consistent and accurate ab initio parametrization of density functional dispersion correction (DFT-D) for the 94 elements H-Pu. J. Chem. Phys. 132, 154104; 10.1063/1.3382344 (2010).

67. Zhao, Y. & Truhlar, D. G. The M06 suite of density functionals for main group thermochemistry, thermochemical kinetics, noncovalent interactions, excited states, and transition elements: two new functionals and systematic testing of four M06-class functionals and 12 other functionals. Theor Chem Account 120, 215–241; 10.1007/s00214-007-0310-x (2008).

68. Weigend, F. & Ahlrichs, R. Balanced basis sets of split valence, triple zeta valence and quadruple zeta valence quality for H to Rn: Design and assessment of accuracy. Physical chemistry chemical physics: PCCP 7, 3297–3305; 10.1039/B508541A (2005).

69. Marenich, A. V., Cramer, C. J. & Truhlar, D. G. Universal solvation model based on solute electron density and on a continuum model of the solvent defined by the bulk dielectric constant and atomic surface tensions. The journal of physical chemistry. B 113, 6378–6396; 10.1021/jp810292n (2009).

70. Frisch, M. J., et al. Gaussiañ16 Revision C.01 (2016).

71. Da Sousa Silva, A. W. & Vranken, W. F. ACPYPE - AnteChamber PYthon Parser interfacE. BMC Res Notes 5, 367; 10.1186/1756-0500-5-367 (2012).

72. Maier, J. A. et al. ff14SB: Improving the Accuracy of Protein Side Chain and Backbone Parameters from ff99SB. Journal of chemical theory and computation 11, 3696–3713; 10.1021/acs.jctc.5b00255 (2015).

73. Jorgensen, W. L., Chandrasekhar, J., Madura, J. D., Impey, R. W. & Klein, M. L. Comparison of simple potential functions for simulating liquid water. J. Chem. Phys. 79, 926–935; 10.1063/1.445869 (1983).

74. Bekker, H., Berendsen, H., Dijkstra, E., Achterop, S. & Renardus, M. Gromacs: A parallel computer for molecular dynamics simulations (1993).

75. Berendsen, H., van der Spoel, D. & van Drunen, R. GROMACS: A message-passing parallel molecular dynamics implementation. Computer Physics Communications 91, 43–56; 10.1016/0010-4655(95)00042-E (1995).

76. van der Spoel, D. et al. GROMACS: fast, flexible, and free. Journal of Computational Chemistry 26, 1701–1718; 10.1002/jcc.20291 (2005).

77. Páll, S., Abraham, M. J., Kutzner, C., Hess, B. & Lindahl, E. Tackling Exascale Software Challenges in Molecular Dynamics Simulations with GROMACS. Proc. EASC 8759, 3– 27; 10.1007/978-3-319-15976-8_1.

78. Abraham, M. J. et al. GROMACS: High performance molecular simulations through multi-level parallelism from laptops to supercomputers. SoftwareX 1–2, 19–25; 10.1016/j.softx.2015.06.001 (2015).

79. Bussi, G., Donadio, D. & Parrinello, M. Canonical sampling through velocity rescaling. J. Chem. Phys. 126, 14101; 10.1063/1.2408420 (2007).

80. Parrinello, M. & Rahman, A. Polymorphic transitions in single crystals: A new molecular dynamics method. J. Appl. Phys. 52, 7182–7190; 10.1063/1.328693 (1981).

81. Hess, B., Bekker, H., Berendsen, H. J. C. & Fraaije, J. G. E. M. LINCS: A linear constraint solver for molecular simulations. J. Comput. Chem. 18, 1463–1472; 10.1002/(SICI)1096-987X(199709)18:12<1463::AID-JCC4>3.0.CO;2-H (1997).

82. van Rossum, G. The Python language reference. 3rd ed. (Python Software Foundation; SoHo Books, Hampton, NH, Redwood City, Calif., 2010).

83. Anaconda Software Distribution. Anaconda Documentation. Available at https://docs.anaconda.com/ (Anaconda Inc, 2020).

84. The Matplotlib Development Team. Matplotlib: Visualization with Python (Zenodo, 2023).

85. Harris, C. R. et al. Array programming with NumPy. Nature 585, 357–362; 10.1038/s41586-020-2649-2 (2020).

