## Supporting Information for "Inhibitor fluorination pattern tunes chemically induced protein dimerization"

**Extended Material and Methods – Synthesis of Tpx inhibitors**

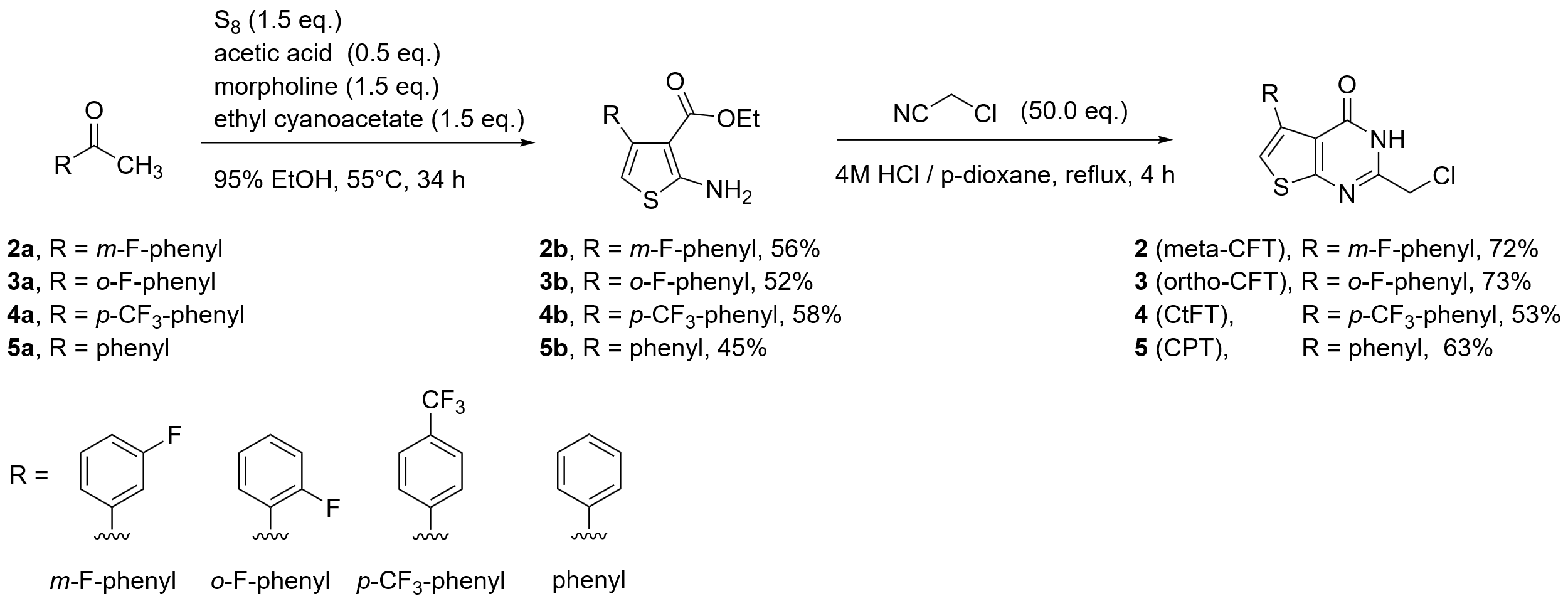

**Supplementary Scheme 1: Synthesis of Tpx inhibitors (2-5).** Following our previously established protocol for para-CFT (**1**)^1^, acetylated aromatic compounds (**2a-5a**) were converted into thiophens (**2b-5b**) using a Gewald-reaction and turned into the final products (**2-5**) via a condensation with chloroacetonitrile.

**General procedure for the Gewald-reaction**

The reaction procedure was adapted with slight modification from Tormyshev et al.^2^ In a screw-cap reaction tube equipped with stirring bar, the respective ketone (3.63 mmol, 1.0 eq.) and ethyl cyanoacetate (5.43 mmol, 1.5 eq.) were dissolved in 0.5 mL 95% ethanol. To the resulting mixture, morpholine (5.43 mmol, 1.5 eq.) and glacial acetic acid (1.81 mmol, 0.5 eq.) were successively added. The reaction mixture was stirred for 4 h at 55 °C before adding three times sulfur (each addition 1.81 mmol, 0.5 eq) over an interval of 10 h. After cooling to room temperature, the reaction mixture was diluted with 50 mL ethyl acetate and extracted three times with 50 mL brine. The organic phase was dried over sodium sulfate, filtered, and concentrated under reduced pressure.

**Ethyl 2-amino-4-(3-fluorophenyl)thiophene-3-carboxylate (2b)**

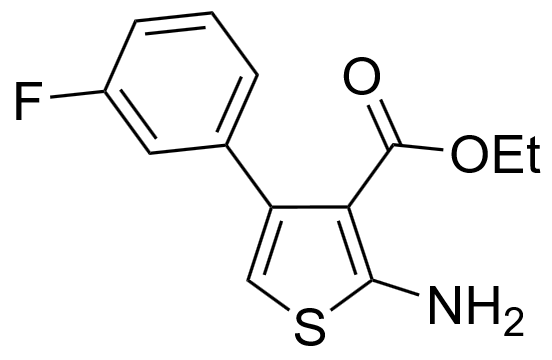
The reaction was performed using 500 mg (3.62 mmol) of 1-(3-fluorophenyl)ethan-1-one **2a**. The crude product was purified using flash column chromatography using 5:1 cyclohexane / ethyl acetate. The product was isolated after recrystallization from 1 mL cyclohexane as yellow crystals (540 mg, 2.04 mmol, 56% Lit.^3^: 53%).

**Melting temperature:** 63.5–67.6 °C (cyclohexane).

**R*_f_* :** 0.3 (5:1 cyclohexane / ethyl acetate).

**IR** (ATR): υ (cm^-1^) = 3424, 3311, 1611, 1485, 1318, 1277, 1235, 1128, 1023, 781, 691.

**^1^H-NMR, COSY (300 MHz, DMSO-*d*_6_)**: δ/ppm = 7.48 – 7.23 (br, m, 3H, -N*H*_2_, *H*-5’), 7.18 – 7.01 (m, 3H, *H*-2’, *H*-4’, *H*-6’), 6.25 (s, 1H, *H*-5), 3.96 (q, *J* = 7.1 Hz, 2H, C*H*_2_), 0.92 (t, *J* = 7.1 Hz, 3H, C*H*_3_).

**^13^C-NMR, HSQC, HMBC (75 MHz, DMSO-*d*_6_)**: δ/ppm = 165.3 (*C_q_*-2), 164.5
(-*C_q_*OO-), 161.4 (d, ^1^*J* (C-F) = 242.1 Hz, *C_q_*-3’), 140.6 (d, ^3^*J* (C-F) = 8.5 Hz, *C_q_*-1’), 139.0 (d, ^4^*J* (C-F) = 2.2 Hz, *C_q_*-4), 129.0 (d, ^3^*J* (C-F) = 8.6 Hz, *C*-5’), 124.8 (d,
^4^*J* (C-F) = 2.6 Hz, *C*-6’), 115.5 (d, ^2^*J* (C-F) = 21.6 Hz, *C*-4’), 113.2 (d,
^2^*J* (C-F) = 20.8 Hz, *C*-2’), 105.9 (*C*-5), 102.6 (*C_q_*-3), 58.7 (*C*H_2_), 13.7 (*C*H_3_).

**^19^F-NMR (282 MHz, DMSO-*d*_6_)**: δ/ppm = -115.7 (ddd, *J* = 10.8, 9.0, 6.0 Hz).

**ESI-HRMS:** calculated for [C_13_H_12_FNO_2_S+H]^+^ m/z = 266.0646, found: m/z = 266.0653.

The analytical data is in accordance with literature.^3^

**Ethyl 2-amino-4-(2-fluorophenyl)thiophene-3-carboxylate (3b)**

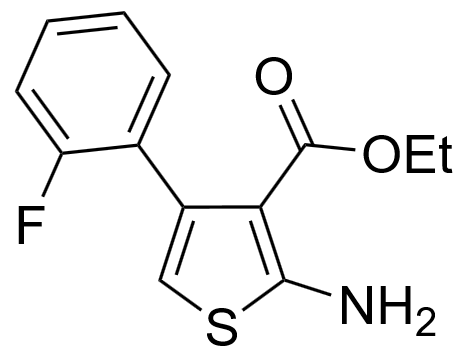
The reaction was performed using 500 mg (3.62 mmol) of 1-(2-fluorophenyl)ethan-1-one **3a**. The crude product was purified using flash column chromatography using 5:1 cyclohexane / ethyl acetate and the pure product was isolated as yellow crystals (500 mg 1.88 mmol, 52%, Lit.^3^: 35%).

**Melting temperature:** 99.7–100.2 °C.

**R*_f_* :** 0.3 (5:1 cyclohexane / ethyl acetate).

**IR** (ATR): υ (cm^-1^): ­3426, 3314, 1580, 1486, 1470, 1321, 1280, 1225, 1136, 1097, 783, 760. **^1^H-NMR, COSY (300 MHz, DMSO-*d*_6_)**: δ/ppm = 7.43 – 7.22 (m, 3H, *H*-Phenyl, -N*H*_2_), 7.20 – 7.06 (m, 2H, *H*-Phenyl, *H*-3’), 6.27 (s, 1H, *H*-5), 3.91 (q, *J* = 7.1 Hz, 2H, C*H*_2_), 0.85 (t, *J* = 7.1 Hz, 3H, C*H*_3_).*

**^13^C-NMR, HSQC, HMBC (75 MHz, DMSO-*d*_6_)**: δ/ppm = 164.6 (-*C_q_*OO-), 164.5 (*C_q_*-2), 159.6 (d, ^1^*J* (C-F) = 244.2 Hz, *C_q_*-2’), 133.6 (*C_q_*-4), 130.6 (d, *J* (C-F) = 3.8 Hz, *C*-Phenyl), 128.9 (d, *J* = 8.2 Hz, *C*-Phenyl), 126.5 (d, ^2^*J* (C-F)= 16.0 Hz, *C_q_*-1’), 123.7 (d, *J* (C-F) = 3.3 Hz), 114.6 (d, ^2^*J* (C-F) = 22.2 Hz, *C*-3’), 106.3 (*C*-5), 103.5 (*C_q_*-3), 58.6 (*C*H_2_), 13.6 (*C*H_3_).*

**^19^F-NMR (282 MHz, DMSO-*d*_6_)**: δ/ppm = -114.8 (ddd, *J* = 10.6, 7.5, 5.1 Hz).

**ESI-HRMS:** calculated for [C_13_H_12_FNO_2_S+H]^+^ m/z = 266.0646, found: m/z = 266.0649.

*A full assignment of all ^1^H-NMR and ^13^C-NMR signals was not possible. The respective signals were indicated as *H*-Phenyl and *C*-Phenyl.

The analytical data is in accordance with literature.^3^

**Ethyl 2-amino-4-(4-(trifluoromethyl)phenyl)thiophene-3-carboxylate (4b)**

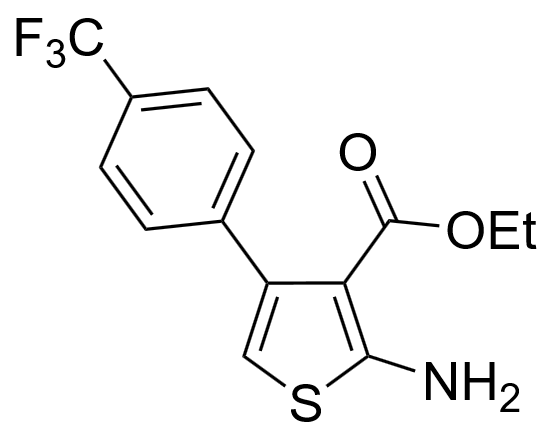
The reaction was performed using 2000 mg (10.63 mmol) of 1-(4-(trifluoromethyl)phenyl)ethan-1-one **4a**. The crude product was purified using flash column chromatography using 5:1 cyclohexane / ethyl acetate. The product was isolated after recrystallization from 4 mL cyclohexane as yellow crystals (1942 mg, 6.16 mmol, 58%, Lit.^3^: 60%).

**Melting temperature:** 102.6–102.9 °C.

**R*_f_* :** 0.3 (5:1 cyclohexane / ethyl acetate).

**IR** (ATR): υ (cm^-1^) = 1639, 1594, 1321, 1271, 1104, 1066, 1019, 784.

**^1^H-NMR, COSY (300 MHz, Chloroform-*d*)**: δ/ppm = 7.63 – 7.53 (m, 2H, *H*-3‘, *H*-5‘), 7.47 – 7.34 (m, 2H, *H*-2‘, *H*-6‘), 6.20 – 6.04 (m, 3H, -N*H*_2_, *H*-5), 4.04 (q, *J* = 7.1 Hz, 2H, C*H*_2_), 0.92 (t, *J* = 7.2 Hz, 3H, C*H*_3_).

**^13^C-NMR, HSQC, HMBC (75 MHz, Chloroform-*d*)**: δ/ppm = 165.5 (-*C_q_*OO-), 164.2 (*C_q_*-2), 142.3 (*C_q_*-1‘), 140.3 (*C_q_*-3), 129.4 (*C*-2‘, *C*-6‘), 129.1 (q, ^2^*J*(C-F) = 32.3 Hz, *C_q_*-4‘), 124.5 ppm (q, ^1^*J*(C-F)=271.9 Hz, *C_q_*F_3_), 124.3 (q, ^3^*J*(C-F) = 3.8 Hz, *C*-3‘, *C*-5‘), 106.4 (*C*-5), 105.9 (*C_q_*-3), 59.8 (*C*H_2_), 13.8 (*C*H_3_).

**^19^F-NMR (282 MHz, Chloroform-*d*)**: δ/ppm = -62.4 (s).

**ESI-HRMS:** calculated for [C_14_H_12_F_3_NO_2_S+H]^+^ m/z = 316.0614, found: m/z = 316.0616.

The analytical data is in accordance with literature.^3,4^

**Ethyl 2-amino-4-phenylthiophene-3-carboxylate (5b)**

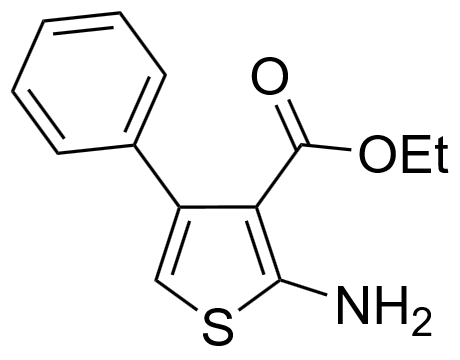
The reaction was performed using 400 mg (3.33 mmol) of acetophenone **5a**. The crude product was purified using flash column chromatography using 5:1 cyclohexane / ethyl acetate. The product was isolated as orange crystals (373 mg, 1.15 mmol, 45%, Lit.^2^: 64%).

**Melting temperature:** 94.0–95.1 °C, (Lit.)^5^: 97–99 °C.

**R*_f_* :** 0.3 (5:1 cyclohexane / ethyl acetate).

**FT-IR** (ATR): υ (cm^-1^) = ­3424, 3311, 1650, 1636, 1580, 1488, 1318, 1279, 1134, 1024, 731, 696.

**^1^H-NMR, COSY (300 MHz, DMSO-*d*_6_)**: δ/ppm = 7.47 – 7.11 (m, 7H, -N*H*_2_, *H*-2’, *H*-3’, *H*-4’, *H*‑5’, *H*-6’), 6.16 (s, 1H, *H*-5), 3.94 (q, *J* = 7.1 Hz, 2H, C*H*_2_), 0.88 (t, *J* = 7.1 Hz, 3H, C*H*_3_).

**^13^C-NMR, HSQC, HMBC (75 MHz, DMSO-*d*_6_)**: δ/ppm = 165.1 (*C_q_*-2),164.7 (-*C_q_*OO-), 140.5 (*C_q_*-4), 138.3 (*C_q_*-1’), 128.6 (*C*-3’, *C*-5’), 127.2 (*C*-2’, *C*-6’), 126.5 (*C*-4’), 105.1 (*C*-5), 102.9 (*C_q_*‑3), 58.6 (*C*H_2_), 13.7 (*C*H_3_).

**ESI-MS:** m/z (%) = 248.1 (100) [M+H]^+^.

The analytical data is in accordance with literature.^2,5^

**General procedure for the pyrimidinone condensation starting from 2-amino-thiophene-3-carboxylates**

The reaction procedure was adapted with slight modification from Klein et al.^1^

In a 50 mL round bottom flask equipped with stirring bar and reflux condenser, a 4 M HCl / p‑dioxane mixture was prepared by slow addition of 4.3 mL of methanol (0.11 mol) to an ice-cooled mixture of acetyl chloride (0.11 mol) and 25 mL p-dioxane. The respective 2-amino-thiophene-3-carboxylate (1.00 mmol, 1 eq.) and chloroacetonitrile (25 mmol, 25 eq.) were added to the acidic solution and heated to reflux for 4 h. After cooling to room temperature, the mixture was diluted with 50 mL of water and extracted three times with 50 mL ethyl acetate. The organic phase was washed once with 50 mL saturated sodium hydrogen carbonate (aq.) solution and 50 mL brine, dried over sodium sulfate and concentrated under reduced pressure.

**2-(Chloromethyl)-5-(3-fluorophenyl)thieno[2,3-*d*]pyrimidin-4(3*H*)-one (meta-CFT, 2)**

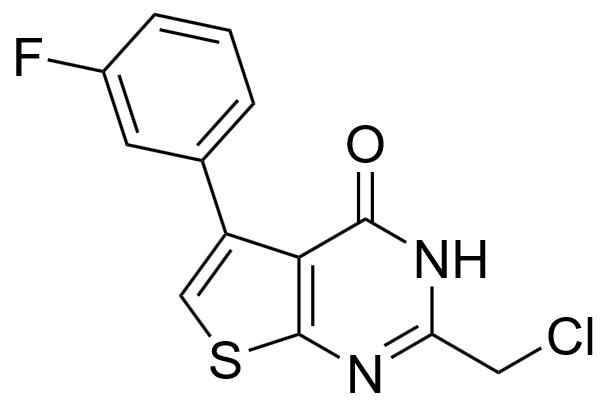
The reaction was performed using 400 mg (1.51 mmol) of compound **2b**. The crude product was purified using flash column chromatography using a 9:1 to 4:1 gradient of cyclohexane / ethyl acetate. The product was isolated as colorless crystals (320 mg, 1.09 mmol, 72%).

**Melting temperature:** Decomposition > 237.3 °C.

**R*_f_* :** 0.3 (2:1 cyclohexane / ethyl acetate).

**IR** (ATR): υ (cm^-1^) = 1651, 1589, 1220, 1171, 1050, 753, 719, 691, 666, 644, 610.

**^1^H-NMR, COSY (300 MHz, DMSO-*d*_6_)**: δ/ppm = 12.82 (s, 1H, N*H*), 7.66 (s, 1H,
*H*-6), 7.51 – 7.32 (m, 3H, *H-*4’, *H-*5’, *H-*6’), 7.29 – 7.09 (m, 1H, *H-*2’), 4.60 (s, 2H, C*H*_2_).

**^13^C-NMR, HSQC, HMBC (75 MHz, DMSO-*d*_6_)**: δ/ppm = 165.3 (*C_q_*-7a), 161.5 (d,
^1^*J* (C-F) = 242.2 Hz, *C_q_*-3’), 157.8 (*C_q_*-4), 153.2 (*C_q_*-2), 137.2 (d, ^3^*J* (C-F) = 8.6 Hz,

*C_q_*-1’), 137.0 (d, ^4^*J* (C-F) = 2.3 Hz, *C_q_*-5), 129.5 (d, ^3^*J* (C-F) = 8.5 Hz, *C*-5’), 125.4 (d, ^4^*J* (C-F) = 2.7 Hz, *C*-6’), 122.6 (*C*-6), 119.8 (*C_q_*-4a), 116.4 (d, ^2^*J* (C-F) = 22.4 Hz,
*C*-4’), 114.2 (d, ^2^*J* (C-F) = 20.9 Hz, *C*-2’), 42.4 (*C*H_2_).

**^19^F-NMR (282 MHz, DMSO-*d*_6_)**: δ/ppm = -114.9 (ddd, *J* = 11.6, 9.2, 5.7 Hz).

**ESI-HRMS:** calculated for [C_13_H_8_ClFN_2_OS+H]^+^ m/z = 295.0103, found: m/z = 295.0114.

**2-(Chloromethyl)-5-(2-fluorophenyl)thieno[2,3-*d*]pyrimidin-4(3*H*)-one (ortho-CFT, 3)**

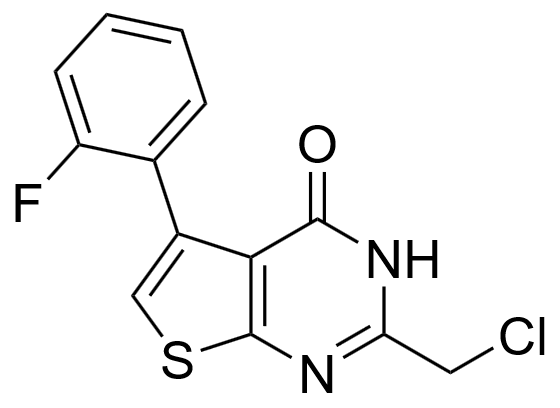
The reaction was performed using 350 mg (1.32 mmol) of compound **3b**. The crude product was purified using flash column chromatography using a 9:1 to 4:1 gradient of cyclohexane / ethyl acetate. The product was isolated as colorless crystals (285 mg, 0.97 mmol, 73%).

**Melting temperature:** Decomposition > 238.5 °C.

**R*_f_* :** 0.3 (2:1 cyclohexane / ethyl acetate).

**IR** (ATR): υ (cm^-1^) = 1651, 1589, 1219, 1171, 1050, 753, 719, 691, 666, 644, 610.

**^1^H-NMR, COSY (300 MHz, DMSO-*d*_6_)**: δ/ppm = 12.77 (s, 1H, N*H*), 7.60 (s, 1H,
*H*-6), 7.53 – 7.37 (m, 2H, *H*-Phenyl), 7.36 – 7.16 (m, 2H, *H*-Phenyl), 4.59 (s, 2H, C*H*_2_).*

(*A full assignment of all ^1^H-NMR and ^13^C-NMR signals was not possible. The respective signals were indicated as *H*-Phenyl and *C*-Phenyl.)

**^13^C-NMR, HSQC, HMBC (75 MHz, DMSO-*d*_6_)**: δ/ppm = 164.3 (*C_q_*-7a), 159.7 (d,
^1^*J* (C-F) = 245.8 Hz, *C_q_*-2’), 157.5 (*C_q_*-4), 153.3 (*C_q_*-2), 131.6 (d, *J* (C-F) = 3.0 Hz,
*C*-Phenyl), 131.2 (*C_q_*-5), 129.9 (d, *J* = 8.3 Hz, *C*-Phenyl), 123.9 (d, *J* = 3.6 Hz,
*C*-Phenyl), 123.4 (d, *J* = 15.5 Hz, *C_q_*-1’), 123.1 (*C*-6), 121.1 (*C_q_*-4a), 115.1 (d,
*J* = 21.9 Hz, *C*-3’), 42.5 (*C*H_2_).

**^19^F-NMR (282 MHz, DMSO-*d*_6_)**: δ/ppm = -114.8 – -114.9 (m).

**ESI-HRMS:** calculated for [C_13_H_8_ClFN_2_OS+H]^+^ m/z = 295.0103, found: m/z = 295.0106.

**2-(Chloromethyl)-5-(4-(trifluoromethyl)phenyl)thieno[2,3-*d*]pyrimidin-4(3*H*)-one (CtFT, 4)**

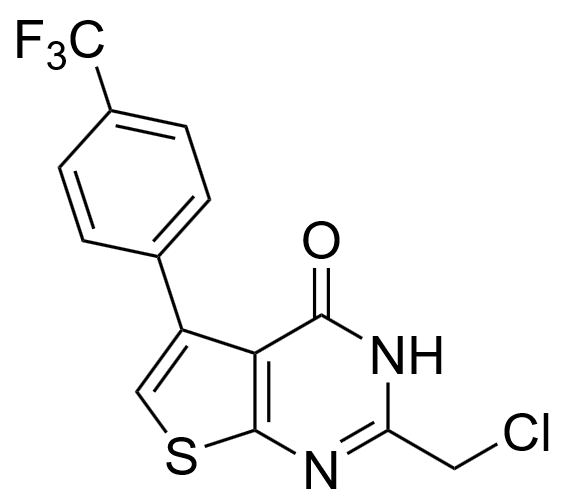
The reaction was performed with 220 mg (0.63 mmol) of compound **6**. The crude product was purified using flash column chromatography using 2:1 cyclohexane / ethyl acetate. The pure product was isolated as brown crystals after recrystallization from 5 mL ethanol (114 mg, 0.33 mmol, 53%).

**Melting temperature:** Decomposition > 244 °C.

**R*_f_* :** 0.5 (2:1 cyclohexane / ethyl acetate).

**IR** (ATR): υ (cm^-1^) = 1653, 1597, 1318, 1176, 1128, 1064, 853, 772, 597, 431.

**^1^H-NMR, COSY (400 MHz, DMSO-*d*_6_)**: δ/ppm = 12.88 (s, 1H, N*H*), 8.07 – 7.41 (m, 5H, *H-*2’, *H-*3’, *H-*5’, *H-*6’, *H-*6), 4.61 (s, 2H, C*H*_2_).

**^13^C-NMR, HSQC, HMBC (101 MHz, DMSO-*d*_6_)**: δ/ppm = 165.4 (*C_q_*-7a), 157.9
(*C_q_*-4), 153.4 (*C_q_*-2), 139.1 (*C_q_*-1’), 136.8 (*C_q_*-5), 130.1 (*C*-2’, *C*-6’), 127.8 (q,
^2^*J* (C-F) = 31.8 Hz, *C_q_*-4’), 124.5 (q, ^3^*J* (C-F) = 3.7 Hz, *C*-3’, *C*-5’), 124.4 (q,
^1^*J* (C-F) = 272.1 Hz, *C_q_*F_3_), 123.3 (*C*-6), 119.8 (*C_q_*-4a), 42.4 ( *C*H_2_).

**^19^F-NMR (282 MHz, DMSO-*d*_6_)**: δ/ppm = -61.4 (s).

**ESI-HRMS:** calculated for [C_14_H_8_ClF_3_N_2_OS+H]^+^ m/z = 345.0071, found: m/z = 345.0076.

**2-(Chloromethyl)-5-phenylthieno[2,3-*d*]pyrimidin-4(3*H*)-one (CPT, 5)**

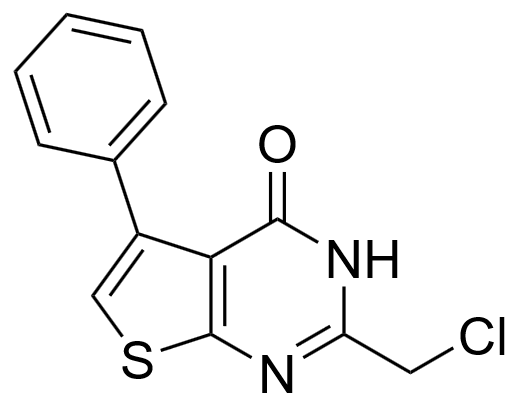
The reaction was performed using 280 mg (1.13 mmol) of compound **5**. The crude product was purified using flash column chromatography using a 6:1 to 2:1 gradient of cyclohexane / ethyl acetate. The pure product was isolated as colorless crystals after recrystallization from 21 mL ethanol (190 mg, 0.71 mmol, 63%).

**Melting temperature:** Decomposition > 227 °C.

**R*_f_* :** 0.3 (2:1 cyclohexane / ethyl acetate).

**IR** (ATR): υ (cm^-1^) = 2839, 1644, 1592, 752, 722, 693, 665, 643, 607.

**^1^H-NMR, COSY (300 MHz, DMSO-*d*_6_)**: δ/ppm = 12.78 (s, 1H, N*H*), 7.63 – 7.50 (m, 3H, *H*-3’, *H*-5’, *H*-6), 7.46 – 7.33 (m, 3H, *H*-2’, *H*-4’, *H*-6’), 4.61 (s, 2H, C*H*_2_).

**^13^C-NMR, HSQC, HMBC (75 MHz, DMSO-*d*_6_)**: δ/ppm = 165.3 (*C_q_*-7a), 157.8
(*C_q_*-4), 153.1 (*C_q_*-2), 138.5 (*C_q_*-1’), 135.1 (*C_q_*-5), 129.4 (*C*-3’, *C*-5’), 127.6 (*C*-2’, *C*-6’), 127.4 (*C*-4’), 121.7 (*C*-6), 119.9 (*C_q_*-4a), 42.5 (*C*H_2_).

**ESI-HRMS:** calculated for [C_13_H_9_ClN_2_OS+H]^+^ m/z = 277.1097, found: m/z = 277.0198.

**Extended Material and Methods – Analysis of MD simulations**

**Dimer interface contacts (i-iv, for Fig. 3 and Supplementary Fig. 11)**

To analyze Tpx WT/inhibitor dimer dissociation and intramolecular protein/inhibitor contacts, we quantified MD simulation frames where critical distances (i-iv) between inter- and intramolecular interaction sites exceeded defined thresholds: (i) inhibitor stacking was monitored via the distance between one inhibitor’s thiophene sulfur to the other inhibitor’s pyrimidinone centroid (*d*_S-PM’_, threshold: 4 Å based on crystal structures of the Tpx/para-CFT dimer and isolated compounds **1-5**). The distance was evaluated using the GROMACS module *gmx pairdist* with the sulfur atom as *reference* and the pyrimidone ring atoms as *selection* using the *-seltype* keyword *part_res_cog*; (ii) the inter-chain salt bridge between K102 and E107’ was monitored via the distance between the C_α_-atoms of both residues to prevent contributions from side chain flexibility (*d*_Cα‑Cα’,_ threshold: 11 Å based on the Tpx/para-CFT dimer structure) and evaluated with *gmx pairdist* using the C_α_-atoms as *reference* and *selection,* respectively; (iii) inhibitor displacement was monitored via the distance between the NH-proton of I109 and the inhibitor’s oxygen (*d*_NH-O_, threshold: 3 Å, based on the typical length of a strong hydrogen bond) and evaluated with *gmx pairdist* with the atoms as *selection* and *reference*; and (iv) W39 displacement was monitored via the distance between the W39 indole NH-proton and the carbonyl oxygen of W70 (*d*_W_, threshold 9 Å) and evaluated with *gmx pairdist*, accordingly (equal to tryptophan state IV in monomeric Tpx/inhibitor complexes, see below for details).

**Inhibitor interactions with Tpx W39A (for Supplementary Fig. 15)**

To analyze the inhibitor orientation in Tpx W39A/inhibitor complexes, the mean distances between protein and small molecule at four interaction sites (backbones of residues E107, S108, I109, as well as the W70 side chain) were determined for each MD simulation and averaged (n= 5). For the interaction with E107, the distance between the residue’s C_α_ proton and the inhibitor’s fluorine atom(s) was determined (*d*_CH-F_) using *gmx pairdist* with the proton as *reference* and the fluorine(s) as selection. For CtFT, the distance to the closest fluorine was evaluated. To check the positioning of the inhibitor’s phenyl moiety, the distances between the phenyl ring centroid (obtained from the command *-seltype part_res_cog* on the ring carbon atoms) and the respective backbone NH proton of Tpx residues S108 and I109 (*d*_NH-Ph_) were determined with *gmx pairdist*. An additional hydrogen bond between the I109 backbone NH-proton and the inhibitor’s oxygen (*d*_NH-O_) was monitored via the distance between these atoms and evaluated with *gmx pairdist*. To investigate inhibitor interactions with the W70 side chain, we also evaluated the distance between the inhibitor’s thiophene ring centroid (obtained from the command -*seltype part_res_cog* on the ring carbon atoms) and the W70 side chain NH-proton (*d*_NH-Th_) with *gmx pairdist*.

**Inhibitor RMSDs of monomeric Tpx/inhibitor complexes (for Fig. 5)**

To analyze the inhibitor’s conformational ensemble in monomeric complexes with Tpx WT and Tpx W39A, we aligned the MD frames of each simulation to the heavy atoms of residue C40 and the W70 backbone, and calculated the RMSD (all heavy atoms of the inhibitor) to the same inhibitor in the pose obtained from our Tpx/para-CFT dimer structure (6GXG, chain A) using *gmx rms* with the respective index groups. RMSDs from five independent MD simulations were averaged for each inhibitor (and, if need be, in “F-in” and “F-out” conformation, respectively).

**Definition and evaluation of tryptophan states (I-IV, for Fig. 5 and Supplementary Fig. 17)**

To analyze the interplay between Tpx-bound inhibitors **1-5** and residue W39, we defined four W-states (I-IV) based on observations from our MD-simulations and structural data from the literature.^1,6–10^ As a proxy for the W39 side chain orientation, we chose to monitor the distance between the W39 indole NH-proton and the backbone oxygen of Tpx residue W70 (*d*_w_) with *gmx pairdist*. We defined W-state I by an intact hydrogen bond between the W39 side chain and the W70 backbone which was assumed for a *d*_w_ < 4 Å. This state is present in the Tpx/para-CFT crystal structure (6GXG, chain C^1^), the Alpha Fold2 prediction for *T. brucei* Tpx WT in the apo state (AF‑O77404‑F1-v4^6,7^), and crystal structures of orthologous proteins (*L. major* Tpx^8^, *C. fasciculata* Tpx^9^, see **Supplementary Fig. 13**). A disruption of this hydrogen bond was termed state II and assumed for a *d*_w_ of 4-7 Å, which for instance can be seen in the crystal structure of the Tpx/para-CFT dimer interface (6GXG, chains A, B^1^). In our MD simulations, we also observed the rare event of a W39 side chain flip (χ2-axis), which led to the formation of a hydrogen bond with the pyrimidinone moiety of inhibitors **1-5**, and was observed for a *d*_w_ of 7-9 Å. A larger distance *d*_w_ (< 9 Å) indicated full solvent exposure of the W39 side chain through rotation (χ1-axis), and was termed state IV, which can be seen in the *T. brucei* Tpx WT apo crystal structure (1O73^10^).

**Supplementary Tables**

**Supplementary Table 1: Cytotoxicity of compounds 1-5 against parasitic and human cells.** EC_50_ values of compounds **1**-**5** for parasitic *T. brucei* *brucei* 449 cells (Lister 427 strain, bloodstream form) were determined with the ATP-lite assay^11^ after 24 and 48 h incubation (two biological replicates of four technical replicates, errors arise from standard deviation). Toxicity against human embryonic kidney (HEK293) cells was determined via resazurin assay^12^ after 24 h incubation time (four technical replicates, errors arise from standard deviation and fitting). The selectivity index (SI) was calculated based on the values obtained from in vivo cell-based toxicity assays (24 h) by dividing the value obtained for human cells by that for *T. brucei* (error calculated following gaussian law of error propagation).

| **Compound** | **Cells** | | **EC_50_ (24 h) / µM** | **EC_50_ (48 h) / µM** | | **SI*_in vivo_* (24 h)** |
| --- | --- | --- | --- | --- | --- | --- |
| para-CFT (**1**) | *T. brucei brucei* 449 | | 1.3 ± 0.3 | 1.5 ± 0.3 | 3.0 ± 0.8 | |
|  | HEK293 | | 3.9 ± 0.4 | / |  | |
| meta-CFT (**2**) | *T. brucei brucei* 449 | | 1.3 ± 0.4 | 1.1 ± 0.5 | 5.0 ± 1.6 | |
|  | HEK293 | | 6.5 ± 0.5 | / |  | |
| ortho-CFT (**3**) | *T. brucei brucei* 449 | | 1.0 ± 0.3 | 0.8 ± 0.3 | 5.0 ± 1.6 | |
|  | HEK293 | | 5.0 ± 0.6 | / |  | |
| CtFT (**4**) | *T. brucei brucei* 449 | | 2.2 ± 0.6 | 2.5 ± 0.6 | 1.8 ± 0.5 | |
|  | HEK293 | | 3.9 ± 0.3 | / |  | |
| CPT (**5**) | *T. brucei brucei* 449 | | 0.6 ± 0.3 | 1.4 ± 0.4 | 8.2 ± 4.2 | |
|  | | HEK293 | 4.9 ± 0.6 | / |  | |

**Supplementary Table 2: Covalent modification of Tpx variants by inhibitors 1-5 and crosslinking agent BM(PEG)_2_.** Purified Tpx WT, Tpx C43S, Tpx C40S, Tpx W39A were incubated with a 3-fold excess of compounds **1**-**5** under reducing conditions and evaluated via intact protein-MS. Tpx WT was additionally crosslinked with BM(PEG)_2_ and analyzed in the same way. The masses of the components, together with the calculated masses for the covalently modified Tpx constructs, and the detected masses via ESI-MS are shown in Dalton (Da). Values for unmodified proteins are shown in italics.

| **Tpx (+ Compound)** | **M_components_ / Da** | **M_calculated_ / Da** | **M_detected_ / Da** |
| --- | --- | --- | --- |
| **Tpx WT_red_** | **16076.2** | 16076.2 | *16076.0* |
| Tpx WT_red_ **+ 1** | 16076.2 **+ 294.7** | 16334.4*^a^* | 16334.0 |
| Tpx WT_red_ **+ 2** | 16076.2 **+ 294.7** | 16334.4*^a^* | 16333.0 |
| Tpx WT_red_ **+ 3** | 16076.2 **+ 294.7** | 16334.4*^a^* | 16334.0 |
| Tpx WT_red_ **+ 4** | 16076.2 **+ 344.7** | 16384.4*^a^* | 16384.0 |
| Tpx WT_red_ **+ 5** | 16076.2 **+ 276.7** | 16316.4*^a^* | 16316.2 |
| Tpx WT_red_ **+ BM(PEG)_2_** | **2x** 16076.2 **+ 308.3** | 32460.7*^b^* | 32460.0 |
| **Tpx C43S** | **16060.1** | 16060.1 | *16059.7* |
| Tpx C43S **+ 1** | 16060.1 **+ 294.7** | 16318.3*^a^* | 16318.0 |
| Tpx C43S **+ 2** | 16060.1 **+ 294.7** | 16318.3*^a^* | 16318.0 |
| Tpx C43S **+ 3** | 16060.1 **+ 294.7** | 16318.3*^a^* | 16318.0 |
| Tpx C43S **+ 4** | 16060.1 **+ 344.7** | 16368.3*^a^* | 16368.0 |
| Tpx C43S **+ 5** | 16060.1 **+ 276.7** | 16300.3*^a^* | 16299.9 |
| **Tpx C40S** | **16060.1** | 16060.1 | *16059.7* |
| Tpx C40S **+ 1** | 16060.1 **+ 294.7** | 16318.3*^a^* | *16059.7* |
| Tpx C40S **+ 2** | 16060.1 **+ 294.7** | 16318.3*^a^* | *16059.8* |
| Tpx C40S **+ 3** | 16060.1 **+ 294.7** | 16318.3*^a^* | *16059.7* |
| Tpx C40S **+ 4** | 16060.1 **+ 344.7** | 16368.3*^a^* | *16059.7* |
| Tpx C40S **+ 5** | 16060.1 **+ 276.7** | 16300.3*^a^* | *16059.7* |
| **Tpx W39A_red_** | **15961.1** | 15961.1 | *15961.*0 |
| Tpx W39A_red_ **+ 1** | 15961.1 **+ 294.7** | 16219.3*^a^* | 16219.4 |
| Tpx W39A_red_ **+ 2** | 15961.1 **+ 294.7** | 16219.3*^a^* | 16219.2 |
| Tpx W39A_red_ **+ 3** | 15961.1 **+ 294.7** | 16219.3*^a^* | 16219.3 |
| Tpx W39A_red_ **+ 4** | 15961.1 **+ 344.7** | 16269.3*^a^* | 16269.3 |
| Tpx W39A_red_ **+ 5** | 15961.1 **+ 276.0** | 16200.6*^a^* | 16201.1 |

*^a^* Covalent **Tpx** complex with compounds **1-5** after HCl elimination (-36.5 Da)

*^b^* Covalent adduct of two **Tpx** molecules, crosslinked via BMPEG_2_

**Supplementary Table 3: Crystal data and structure refinement for compounds 2-5. The crystal structure of isolated para-CFT (1) has been published in Wagner et al**.^1^

| **Crystal data and structure refinement for meta-CFT (2)** | |
| --- | --- |
| Empirical formula | C_13_H_8_ClFN_2_OS |
| Formula weight | 294.7 |
| Temperature/ K | 120 |
| Crystal system | monoclinic |
| Space group | P 2_1_/n |
| a/ Å | 9.7655(7) |
| b/ Å | 9.6269(4) |
| c/ Å | 13.3940(9) |
| α/ ° | 90.0 |
| ß/ ° | 105.817(5) |
| γ/ ° | 90.0 |
| Volume/ Å^3^ | 1211.51(13) |
| Z | 4 |
| ρ_calc_/ g cm^-3^ | 1.616 |
| μ/ mm^-1^ | 0.491 |
| F(000) | 600 |
| Crystal size/ mm³ | 0.11 x 0.22 x 0.30 |
| Radiation | Mo-Kα |
| 2θ range for data collection/ ° | 4 to 56 |
| Index ranges | -12 ≤ h ≤ 12 -12 ≤ k ≤ 12 -16 ≤ l ≤ 17 |
| Reflections collected | 8043 |
| Independent reflections | 2904 (R_int_ = 0.0129) |
| Data/restraints/parameters | 2904/0/208 |
| Goodness-of-fit on F² | 1.068 |
| Final R indexes [I>2σ(I)] | R1= 0.0292, wR2= 0.0753 |
| Final R indexes [all data] | R1= 0.0329, wR2= 0.0782 |
| Largest diff. peak/hole / eÅ^-3^ | 0.44, -0.15 |
| **Crystal data and structure refinement for ortho-CFT (3)** | |
| Empirical formula | C_13_H_8_ClFN_2_OS |
| Formula weight | 294.7 |
| Temperature/ K | 120 |
| Crystal system | monoclinic |
| Space group | P 2_1_/n |
| a/ Å | 9.6949(5) |
| b/ Å | 9.5481(5) |
| c/ Å | 13.7042(7) |
| α/ ° | 90.0 |
| ß/ ° | 108.088(4) |
| γ/ ° | 90.0 |
| Volume/ Å^3^ | 1205.87(11) |
| Z | 4 |
| ρ_calc_/ g cm^-3^ | 1.623 |
| μ/ mm^-1^ | 0.493 |
| F(000) | 600 |
| Crystal size /mm³ | 0.04 x 0.18 x 0.41 |
| Radiation Mo-Kα | Mo-Kα |
| 2θ range for data collection/ ° | 4 to 56 |
| Index ranges | -12 ≤ h ≤ 11 -12 ≤ k ≤ 12 -17 ≤ l ≤ 18 |
| Reflections collected | 6502 |
| Independent reflections | 2874 (R_int_ = 0.0248) |
| Data/restraints/parameters | 2874/1/203 |
| Goodness-of-fit on F² | 1.023 |
| Final R indexes [I>2σ(I)] | R1= 0.0371, wR2= 0.0844 |
| Final R indexes [all data] | R1= 0.0559, wR2= 0.0923 |
| Largest diff. peak/hole / eÅ^-3^ | 0.37, -0.31 |

**Supplementary Table 3, continued:**

| **Crystal data and structure refinement for CtFT (4)** | |
| --- | --- |
| Empirical formula | C_14_H_8_ClF_3_N_2_OS |
| Formula weight | 344.7 |
| Temperature/ K | 120 |
| Crystal system | monoclinic |
| Space group | P 2_1_ |
| a/ Å | 10.7196(7) |
| b/ Å | 7.9153(3) |
| c/ Å | 16.2165(11) |
| α/ ° | 90.0 |
| ß/ ° | 92.519(5) |
| γ/ ° | 90.0 |
| Volume/ Å^3^ | 1374.63(14) |
| Z | 4 |
| ρ_calc_/ g cm^-3^ | 1.666 |
| μ/ mm^-1^ | 0.466 |
| F(000) | 696 |
| Crystal size/ mm³ | 0.1 x 0.14 x 1.3 |
| Radiation | Mo-Kα |
| 2θ range for data collection/ ° | 4 to 56 |
| Index ranges | -14 ≤ h ≤ 14 -10 ≤ k ≤ 9 -21 ≤ l ≤ 21 |
| Reflections collected | 15442 |
| Independent reflections | 6426 (R_int_ = 0.0343) |
| Data/restraints/parameters | 6426/1/459 |
| Goodness-of-fit on F² | 1.086 |
| Final R indexes [I>2σ(I)] | R1= 0.0465, wR2= 0.1137 |
| Final R indexes [all data] | R1= 0.0612, wR2= 0.1201 |
| Largest diff. peak/hole/ eÅ^-3^ | 0.42, -0.38 |
| **Crystal data and structure refinement for CPT (5)** | |
| Empirical formula | C_13_H_9_ClN_2_OS |
| Formula weight | 276.7 |
| Temperature/ K | 120 |
| Crystal system | monoclinic |
| Space group | C 2/c |
| a/ Å | 21.2954(15) |
| b/ Å | 13.2035(13) |
| c/ Å | 8.6076(6) |
| α/ ° | 90.0 |
| ß/ ° | 97.275(6) |
| γ/ ° | 90.0 |
| Volume/ Å^3^ | 2400.7(3) |
| Z | 8 |
| ρcalc/gcm^-3^ | 1.531 |
| μ/ mm^-1^ | 0.479 |
| F(000) | 1136 |
| Crystal size /mm³ | 0.11 x 0.48 x 0.70 |
| Radiation | Mo-Kα |
| 2θ range for data collection /° | 4 to 56 |
| Index ranges | -27 ≤ h ≤ 27 -16 ≤ k ≤ 17 -8 ≤ l ≤ 11 |
| Reflections collected | 5925 |
| Independent reflections | 2856 (R_int_= 0.0166) |
| Data/restraints/parameters | 2856/0/198 |
| Goodness-of-fit on F² | 1.093 |
| Final R indexes [I>2σ(I)] | R1= 0.0329, wR2= 0.0787 |
| Final R indexes [all data] | R1= 0.040, wR2= 0.0841 |
| Largest diff. peak/hole / eÅ^-3^ | 0.43, -0.18 |

**Supplementary Table 4: ^19^F-NMR signal chemical shifts and linewidths of free and Tpx-bound inhibitors 1‑4.** Chemical shifts were referenced to TFA. For Tpx-bound molecules, samples were pre-incubated and purified via SEC as described in the material and methods section. To obtain spectra for free compounds **1-4** in Tpx buffer, 8 or 13 vol% DMSO were added to dissolve the molecules, requiring the introduction of a correction summand for the resulting ^19^F-chemical shifts (corrections are shown in the bottom right and are already included in the chemical shifts shown in this table). 1D ^19^F NMR spectra are shown in **Supplementary Fig. 18**.

| **Tpx/inhibitor**  **complexes** |  | **Tpx WT + inhibitor** | |  | **Tpx W39A + inhibitor** | |
| --- | --- | --- | --- | --- | --- | --- |
|  | **conc.**  **/ µM** | **chemical shift**  **/ ppm** | **linewidth / Hz** |  | **chemical shift**  **/ ppm** | **linewidth**  **/ Hz** |
| para-CFT (**1**) | 750 | -115.58 | 118 |  | -115.07 | 84 |
|  | 500 | -115.58 | 111 |  | -115.02 | 80 |
|  | 250 | -115.58 | 118 |  | -114.95 | 70 |
|  | 100 | -115.52 | 144***** |  | -114.88 | 54 |
|  | 50 | -115.54 | **/*** |  | -114.85 | 41 |
| meta-CFT (**2**) | 750 | -114.43 | 102 |  | -114.73 | 56 |
|  | 500 | -114.43 | 100 |  | -114.74 | 54 |
|  | 250 | -114.43 | 103 |  | -114.75 | 50 |
|  | 100 | -114.44 | 96 |  | -114.77 | 46 |
|  | 50 | -114.42 | 93 |  | -114.78 | 45 |
| ortho-CFT (**3**) | 750 | -114.41 | 307 |  | -115.28 | 63 |
|  | 500 | -114.41 | 320 |  | -115.31 | 53 |
|  | 250 | -114.32 | 324 |  | -115.35 | 56 |
|  | 100 | -114.41 | **/*** |  | -115.38 | 52 |
|  | 50 | **/*** | **/*** |  | -115.39 | 51 |
| CtFT (**4**) | 750 | -62.80 | 96 |  | -62.96 | 33 |
|  | 500 | -62.82 | 98 |  | -62.97 | 32 |
|  | 250 | -62.85 | 80 |  | -62.98 | 32 |
|  | 100 | -62.89 | 74 |  | -62.98 | 31 |
|  | 50 | -62.92 | 58 |  | -62.98 | 30 |
| **Free**  **inhibitors** | **conc.**  **/ µM** | **chemical shift****  **/ ppm** | **linewidth**  **/ Hz** |  | **DMSO**  **/ vol.%** | **correction**  **/ ppm** |
| para-CFT (**1**) | 100 | -115.98 | 31 |  | 8 | -0.15 |
| meta-CFT (**2**) | 100 | -115.50 | 31 |  | 8 | -0.15 |
| ortho-CFT (**3**) | 100 | -116.47 | 31 |  | 8 | -0.15 |
| CtFT (**4**) | 100 | -63.11 | 26 |  | 13 | -0.26 |

*poor signal to noise ratio

**chemical shifts for free inhibitors already include DMSO correction term

**Supplementary Figures**

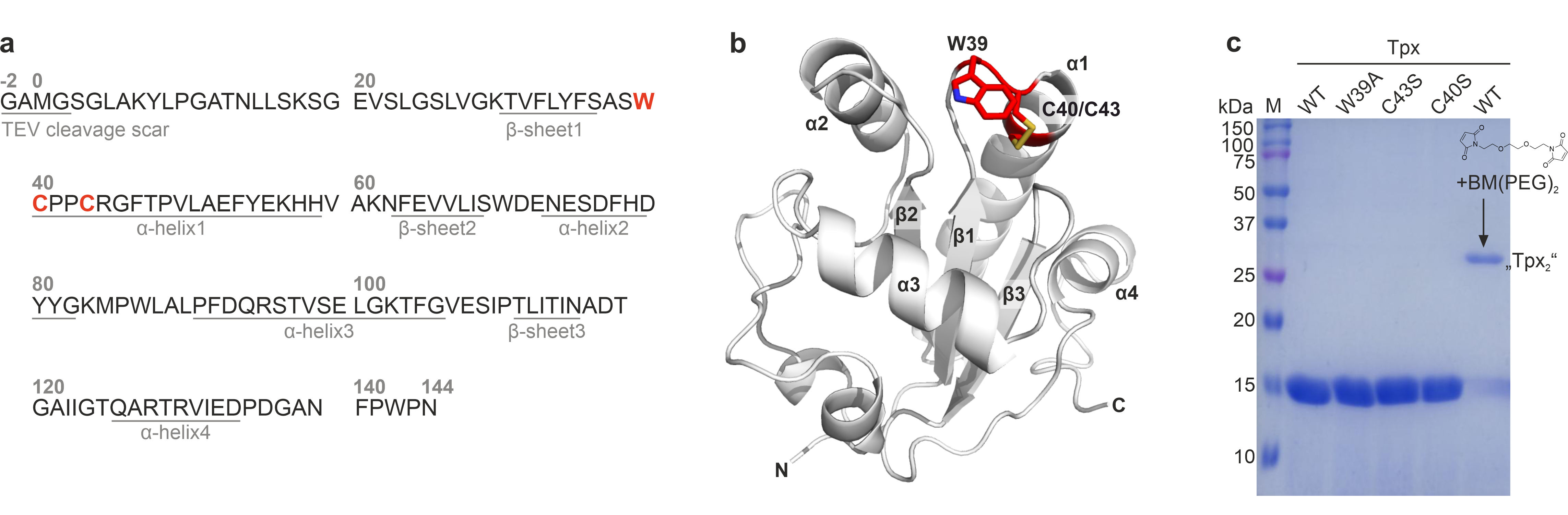

**Supplementary Fig. 1:** **Tpx constructs used in this study.** **a**, Sequence of the *T. brucei* Tpx wildtype (WT) protein (Uniprot accession: O77404). Tpx was purified with an N-terminal TEV protease recognition site, resulting in replacement of methionine residue 1 with a “GAMG” motif as a scar from TEV-cleavage in the final construct. Major secondary structure motifs (consisting of at least five amino acids) are underlined and numbered. Residues mutated in this study (W39, C40, C43) are highlighted in red**. b,** Tertiary structure of *T. brucei* Tpx WT. For clarity, we show the AlphaFold prediction (AF-O77404-F1-model_v4^6,7^) where the W39 indole conformation agrees with other published Tpx structures (e.g. *Crithidia fasciculata* Tpx1, PDB: 1EWX^9^, and *Leishmania major* Tpx1, PDB: 3S9F^8^) instead of the crystal structure of *T. brucei* Tpx (PDB:1O73)^10^, where this residue adopts a different conformation (see also **Supplementary Fig. 13**). The RMSD between the *T. brucei* Tpx WT crystal structure (PDB: 1O73^10^) and the AlphaFold prediction (AF-O77404-F1-model_v4^6,7^) is 0.55 Å. **c**, Purified Tpx WT, Tpx W39A, Tpx C43S, Tpx C40S (all ~16 kDa), and BM(PEG)_2_-crosslinked Tpx WT (~32 kDa, “Tpx_2_”) on a 15% SDS-PAGE gel (Coomassie-stained, marker: Biorad Precision Plus Protein Dual Color Standard). The structural formula of the Tpx crosslinker 1,8-bismaleimido-diethyleneglycol (BM(PEG)_2_) is shown in the last lane, above the protein band of crosslinked Tpx (“Tpx_2_”).

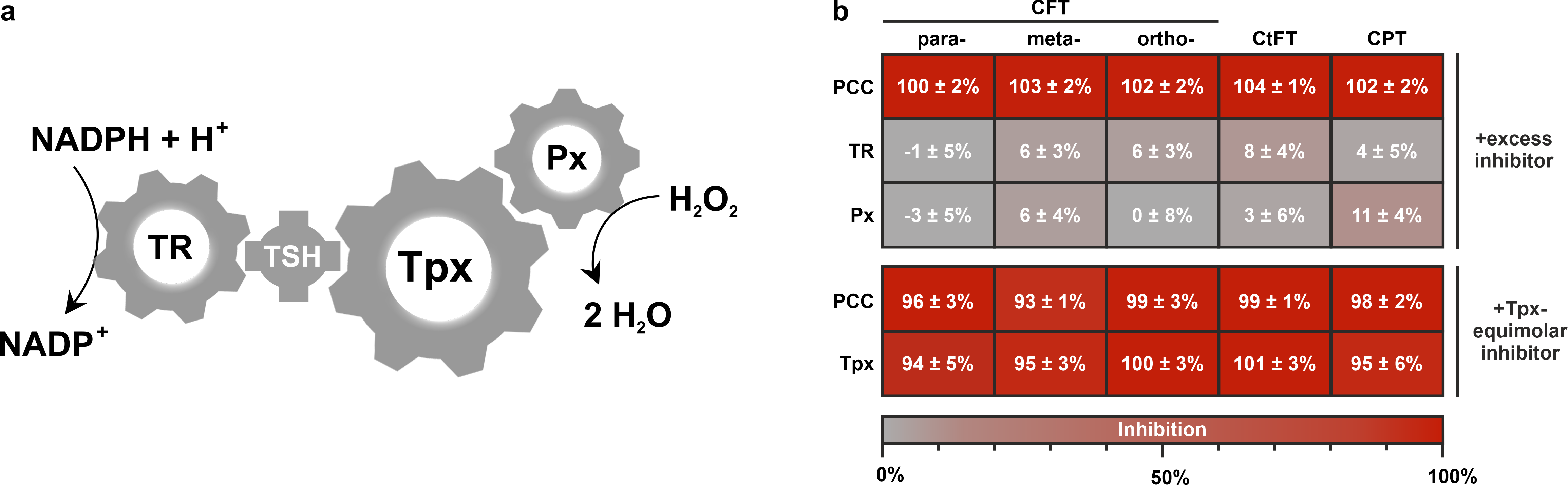

**Supplementary Fig. 2**: **Compounds 1-5 are selective Tpx inhibitors in vitro.** **a**, Schematic representation of the trypanosomal peroxide clearance cascade (PCC) consisting of trypanothione reductase (TR), trypanothione (TSH), tryparedoxin (Tpx), and a peroxidase (Px). In each enzymatic step, the transfer of reducing equivalents is accomplished through a thiol exchange reaction, which highlights the large number of reactive cysteine species as potential inhibitor off-targets in the parasitic redox cascade. **b,** Top: Relative enzyme inhibition after treatment of the entire PCC, or TR and Px separately, with excess amounts of inhibitors **1-5** (for details, see material and methods section). Bottom: Treatment of the entire PCC or Tpx alone with Tpx-equimolar amounts of compounds **1-5**.

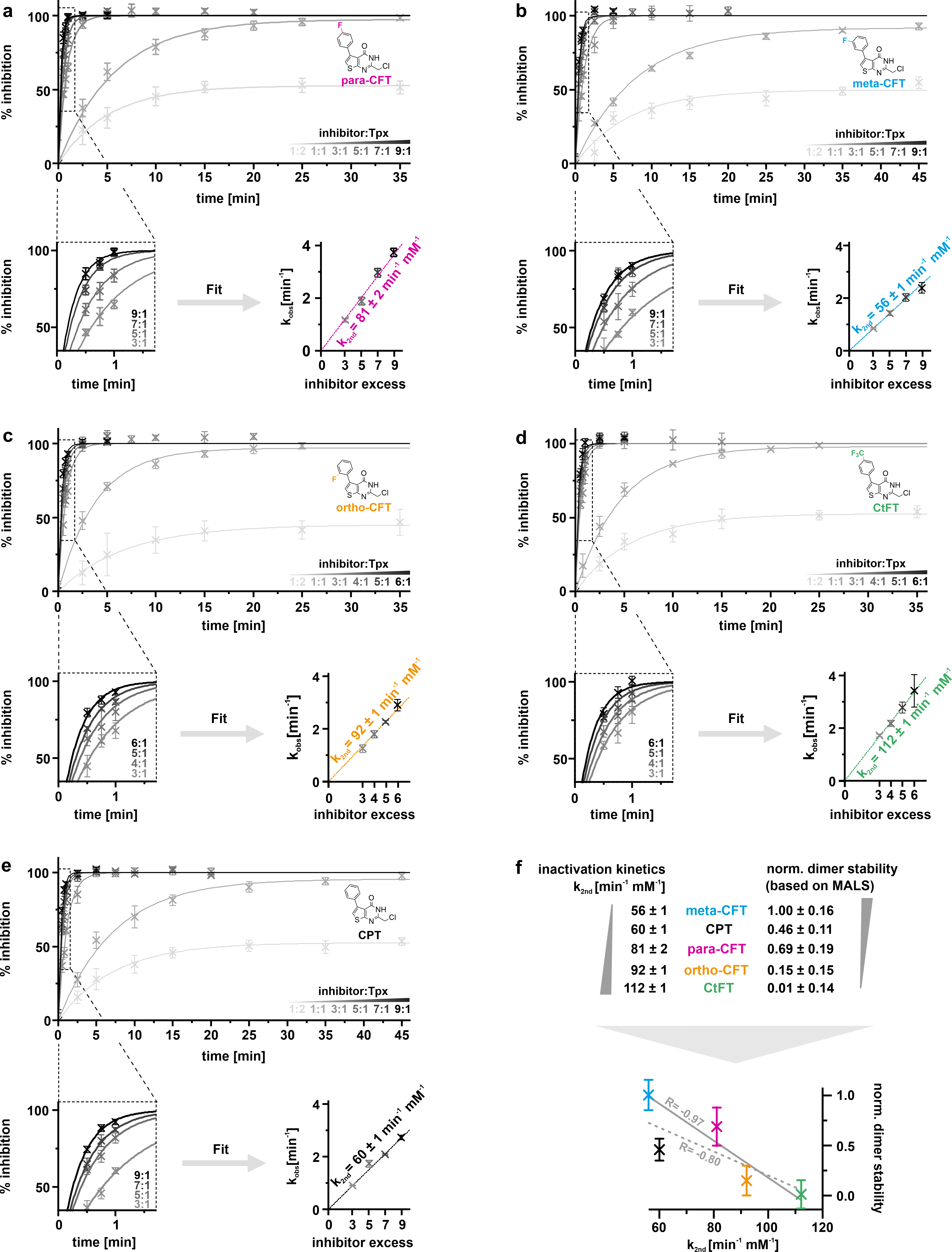

**Supplementary Fig. 3**: **Kinetics of covalent Tpx inhibition are fluorination dependent and inversely correlated with induced dimer affinity. a-e**, Using the reconstituted peroxide clearance cascade, time-resolved inhibition curves of 5 µM Tpx with substoichiometric, equimolar and excess amounts of inhibitors **1-5** (3-, 5-, 7-, and 9-fold excess for slow inactivators (**1**-**2**, **5**); 3-, 4-, 5-, and 6-fold excess for fast inactivators (**3**-**4**)) were recorded at 25 °C. Curves were fitted as a one phase exponential association with f(x)= I_max_*(1-exp(-k_obs_*x)), with I_max_ as the inhibition limit and k_obs_ as the observed experimental inhibition constant (**a**-**e** top). A close-up (bottom left in each panel) shows inhibition shortly after addition of excess amount of inhibitor. Observed reaction rate constants (k_obs_) were plotted against inhibitor concentration, shown here as x-fold inhibitor excess. The slope of a linear regression fit (dashed line) yielded second order rate constants k_2nd_ (bottom right in each panel). Data are shown as mean value ± standard deviation of three technical triplicates. **f**: Correlation of inhibition kinetics (k_2nd_) with dimer affinities (based on SEC-MALS with 100 µM Tpx in complex with inhibitor). Data were normalized to the most stable complex (Tpx/meta-CFT) as assessed by ITC, SEC and SEC-MALS (see **Fig. 2** and main text). Correlation yielded negative Pearson R values (fluorinated inhibitors (**1-4**): solid line; all inhibitors (**1-5**): dashed line), showing that inhibition kinetics are slow for potent Tpx dimerizers and vice versa.

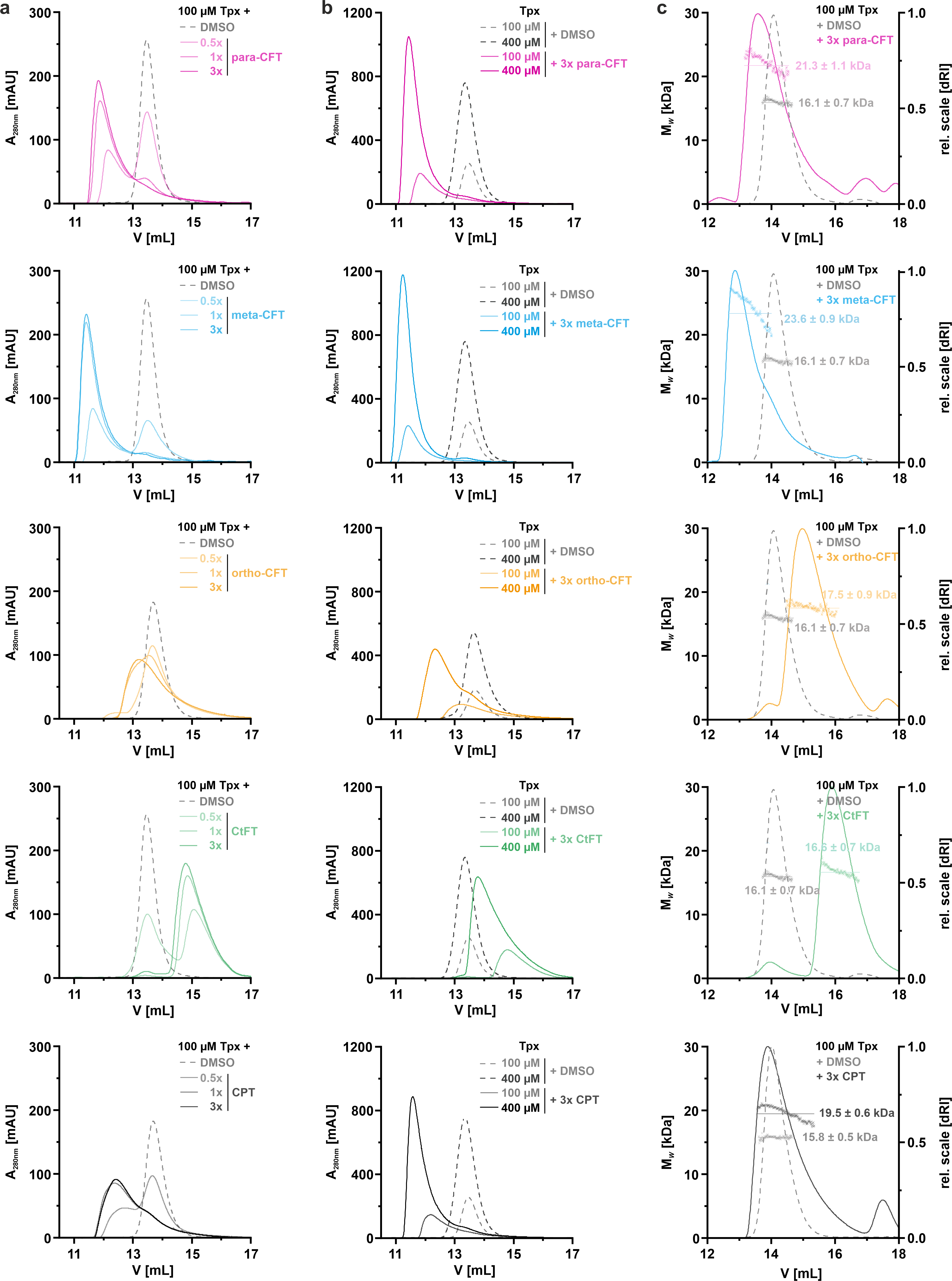
 **Supplementary Fig. 4:** **Analytical SEC and SEC-MALS of Tpx and Tpx/inhibitor complexes**. **a**, Elution profiles of reduced Tpx WT (100µM) with 7.5% DMSO (grey dashed line), and with inhibitors **1**-**5** at substoichiometric (1:0.5), equimolar (1:1) and excess (1:3) molar amounts (solid lines). Inhibitor binding caused smaller Tpx elution volumes except for CtFT (**4**), which does not lead to notable dimerization at 100 µM protein concentration and enhances column interactions (see **Supplementary Fig. 5**). **b**, Concentration dependent elution profiles of 100 µM or 400 µM unmodified Tpx in the reduced state (with 7.5% and 30% DMSO, respectively) compared to Tpx/inhibitor complexes generated by threefold molar excess of inhibitor. In all cases, a higher protein concentration shifted the Tpx elution volume towards the permanent, BM(PEG)_2_-linked dimer (V= 11.3 mL). **c**, SEC-MALS of Tpx and Tpx/inhibitor complexes (100 µM each) with changes in refractive index (*d*RI, right y-axis) and average molar mass (*M*_w_, shown as crosses (left y-axis)). Except for the CtFT complex, higher *M*_w_ were observed for modified than unmodified Tpx.

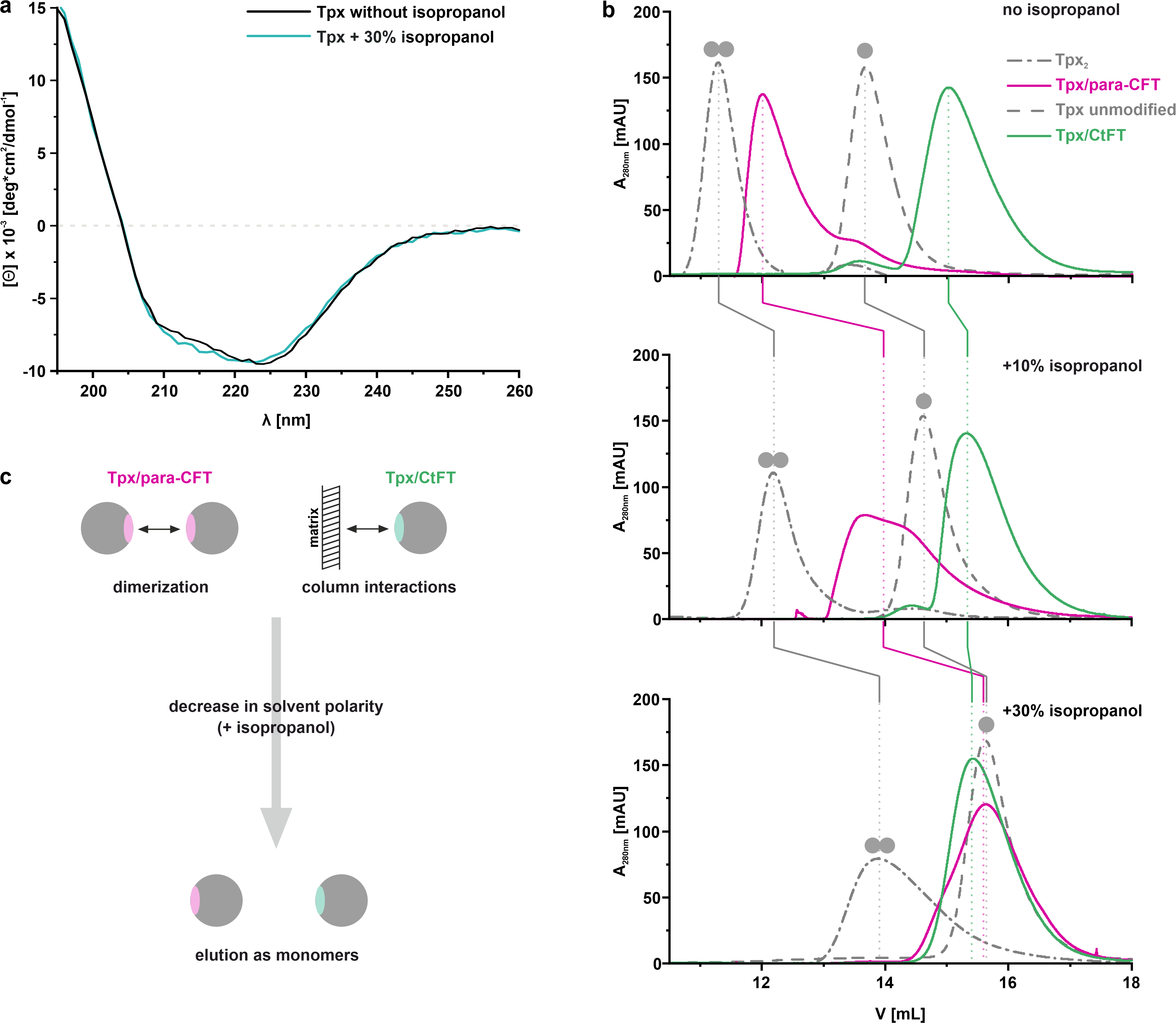

**Supplementary Fig. 5: Delayed SEC elution of the Tpx WT/CtFT complex is due to secondary column interactions. a,** To mimic the length of a SEC run, Tpx was incubated for 30 min at room temperature in SEC buffer without (black) or complemented with 30% v/v isopropanol (petrol). Circular dichroism (CD) spectra show that the structural integrity of Tpx was not affected by isopropanol. **b,** SEC elution profiles of Tpx monomers (Tpx, dashed grey line), BM(PEG)_2_-crosslinked Tpx dimers (Tpx_2_, dot-dashed grey line), Tpx/para-CFT (pink line) and Tpx/CtFT complexes (green line) in SEC buffer without isopropanol (top), with 10% v/v isopropanol (middle), and with 30% v/v isopropanol (bottom). Isopropanol increases the hydrophobicity of the mobile phase and thus weakens hydrophobic interactions of analytes with themselves (e.g. dimerization) as well as with the column matrix. For the monomeric and dimeric Tpx standard, the oligomeric state is indicated by grey spheres above the respective elution peaks. In buffer without isopropanol, Tpx/para-CFT dimers eluted earlier and Tpx/CtFT later than unmodified Tpx. In buffer with 30% v/v isopropanol, unmodified and inhibitor-bound Tpx eluted similarly, suggesting that all proteins are monomeric under these conditions. In contrast, the permanent, crosslinked Tpx dimer, albeit also affected by the solvent polarity, still eluted at smaller elution volumes showing that the column still separated by particle size under these conditions. **c**, Model for secondary analyte interactions that affect the SEC elution volume. At high solvent polarity (no isopropanol), dimer interactions reduce the Tpx/para-CFT elution volume, while for Tpx/CtFT, column interactions lead to the observed larger elution volume. With 30% v/v isopropanol, both protein/inhibitor complexes elute as monomers because attractive dimer and column interactions are weakened. Increased SEC-elution volumes due to hydrophobic interactions with the column have been reported previously.^13,14^

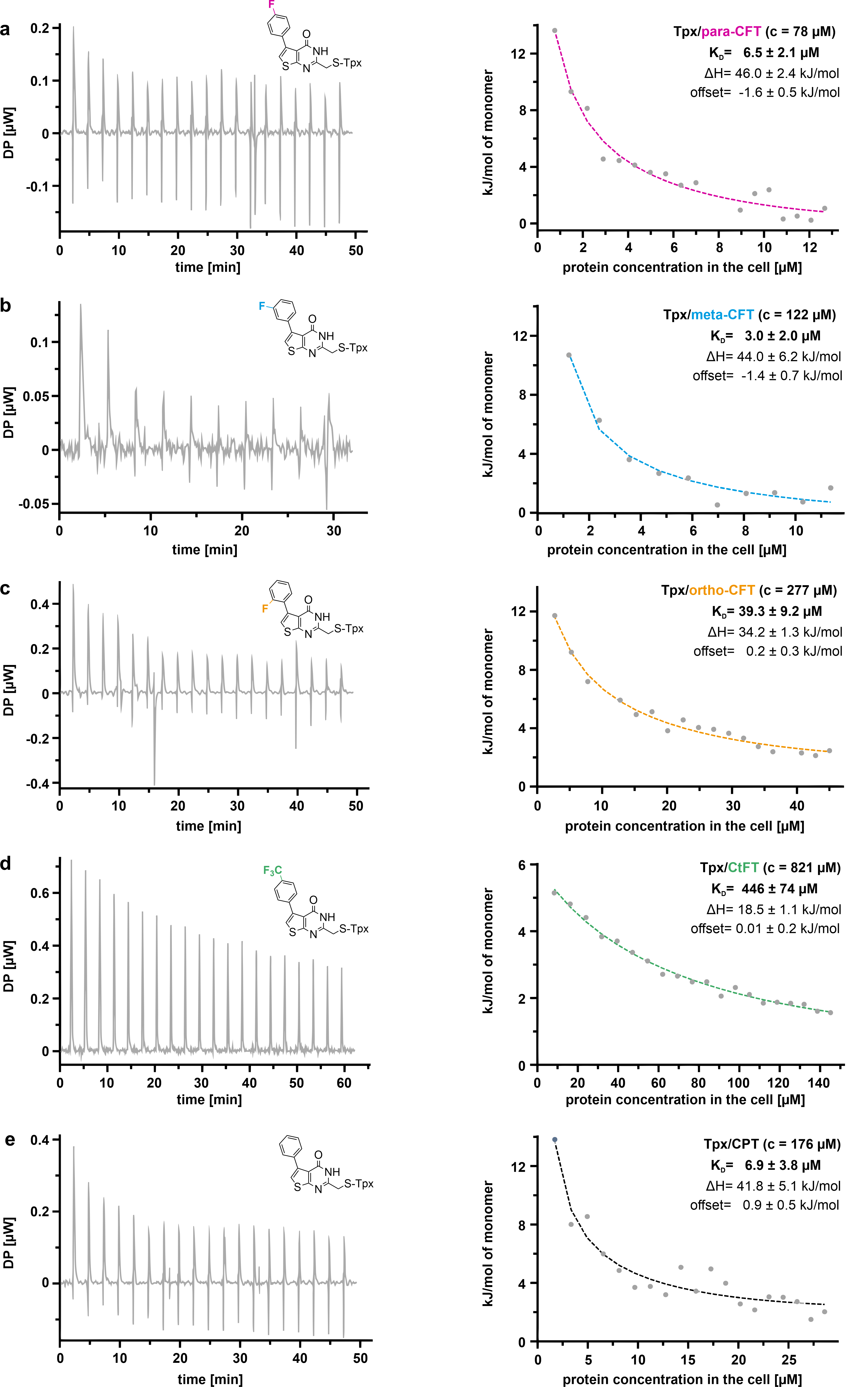

**Supplementary Fig. 6: Dissociation constants (*K*_D_) for dimeric Tpx/inhibitor complexes. a-e,** Dilution isothermal titration calorimetry (ITC) of covalent Tpx/inhibitor complexes. Thermograms after serial injections of the analyte into Tpx buffer are shown in the left panels. Integrals of thermal signals were fitted to monomer equivalents in the cell (right panels) to yield dissociation constants *K*_D_, dissociation enthalpies ΔH, and offsets as fitting parameters. Protein concentration in the syringe is given in brackets. Each measurement was performed in a technical triplicate (for simplicity, only one replicate is shown here for each analyte). Dimer dissociation was endothermic, and integrals decreased with increasing analyte concentration in the measurement cell.

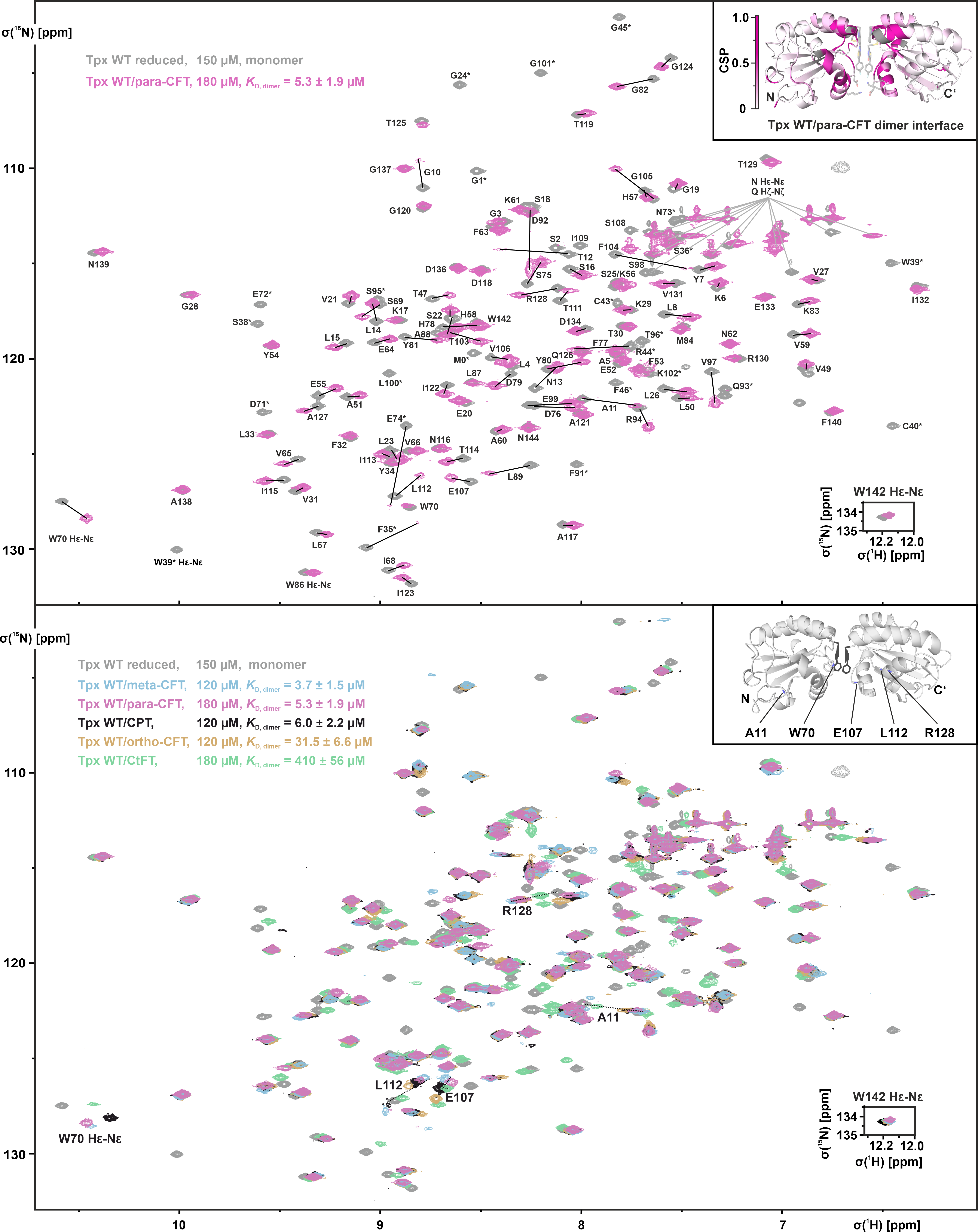

**Supplementary Fig. 7: Tpx WT interactions with inhibitors 1-5 monitored by solution NMR spectroscopy. Top:** 2D ^1^H, ^15^N-HSQC spectra of unmodified, monomeric ^15^N-labeled Tpx WT in the reduced state (150 µM, black) and the dimeric Tpx WT/para-CFT complex (180 µM, pink). The backbone assignments for the para-CFT bound state were transferred from our previous assignment for the reduced Tpx WT^15^, and verified through additional 3D experiments. Signals displaying line broadening upon inhibitor binding (not visible in the spectrum) are marked with an asterisk. The dissociation constant (*K*_D_) of para-CFT (**1**)-induced Tpx dimerization (top left) was obtained from dilution ITC**. Bottom:** ^1^H, ^15^N-HSQC spectra of free and inhibitor bound Tpx WT (concentrations and dimer *K*_D_s shown in the top left). Backbone assignments were transferred from our previous assignment of reduced Tpx WT^15^ and, where necessary, verified through additional 3D experiments for the inhibitor-bound states. Inhibitor binding led to signal broadening of residues constituting the dimer interface, and additional chemical shift perturbations of backbone and side chain resonances of residues throughout the protein. Inhibitor fluorination pattern-dependent chemical shift trajectories for selected amide resonances (A11, W70 side chain, E107, L112 and R128) are indicated by dotted lines. In most cases, the extent of the chemical shift followed the dissociation constants *K*_D_ of the inhibitor-induced dimers, suggesting that Tpx dimerization is the primary cause. Trajectories not following this trend included residues in direct contact with inhibitors **1-5** (e.g. E107 amide backbone and W70 side chain, see **Fig. 3b** as well as main text for details). The dissociation constants (*K*_D_) for inhibitor-induced Tpx dimerization were obtained from dilution ITC.

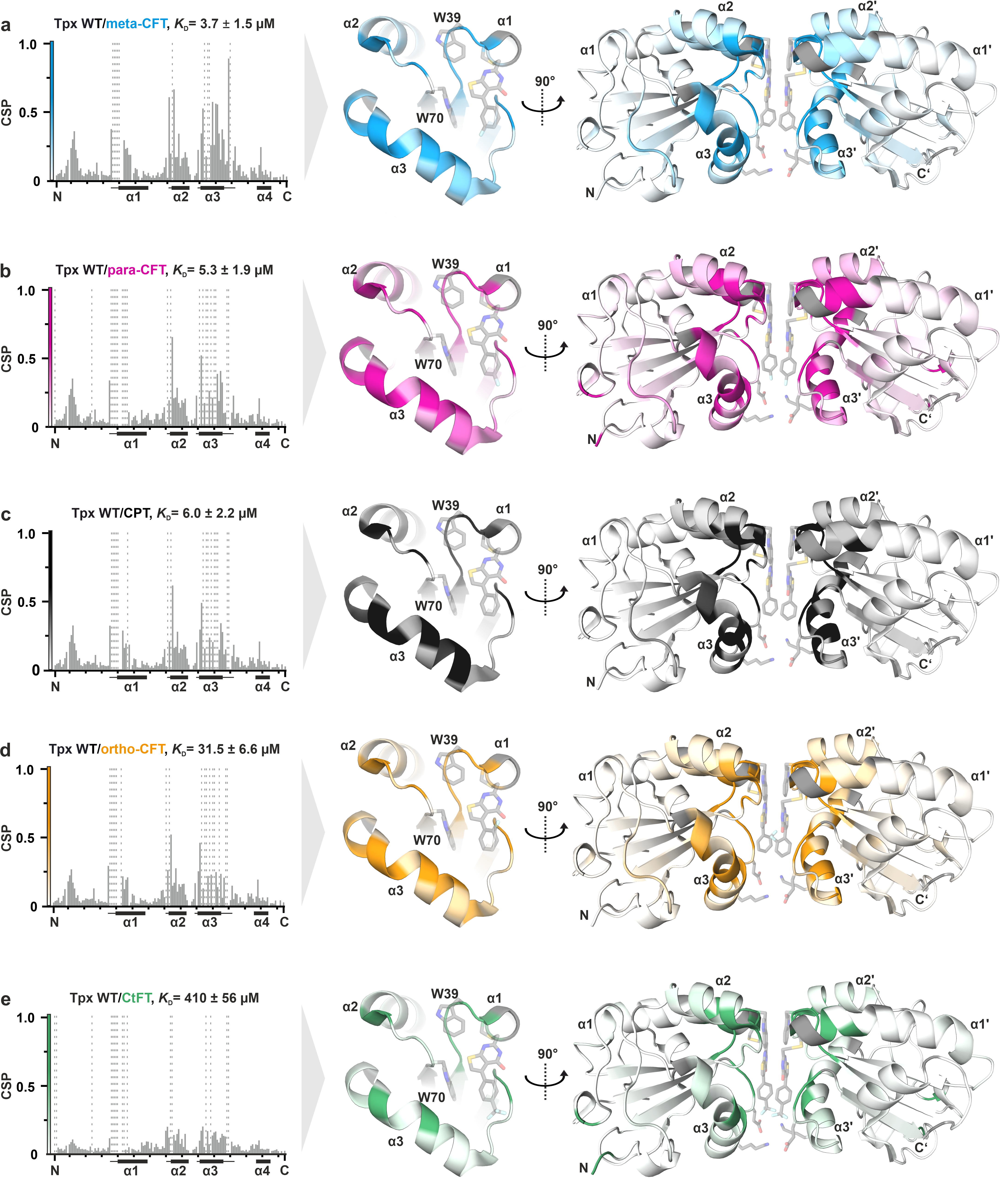

**Supplementary Fig. 8: Inhibitor interactions and chemically induced dimerization of Tpx WT monitored by ^1^H, ^15^N-NMR spectroscopy.** **a-e,** Inhibitor induced chemical shift perturbations (CSP) in ^1^H, ^15^N-HSQC spectra of Tpx WT (see **Supplementary Fig. 7**). The CSP plots (left) start with residue G1 and end with N144 (indicated with “N” and “C”, respectively), ticks indicate increments of ten, starting with G10. The Tpx α-helices are indicated by black cylinders on the x-axis and adjacent loops with high CSP are indicated by lines. Signals displaying line broadening are shown as dashed bars. For each inhibitor, CSP upon inhibitor binding were also mapped onto the structure of Tpx/para-CFT dimers (PDB: 6GXG^1^, Chains A and B), and are shown in color on an open book representation of the Tpx dimer interface (center), and in the context of the induced Tpx dimer (right, copied from **Fig. 3a** for clarity). All Tpx dimer interfaces consist of the respective inhibitor, Tpx α-helices 1-3, and loops containing W39 and W70 (for simplicity, W70 is not shown in the Tpx dimer).

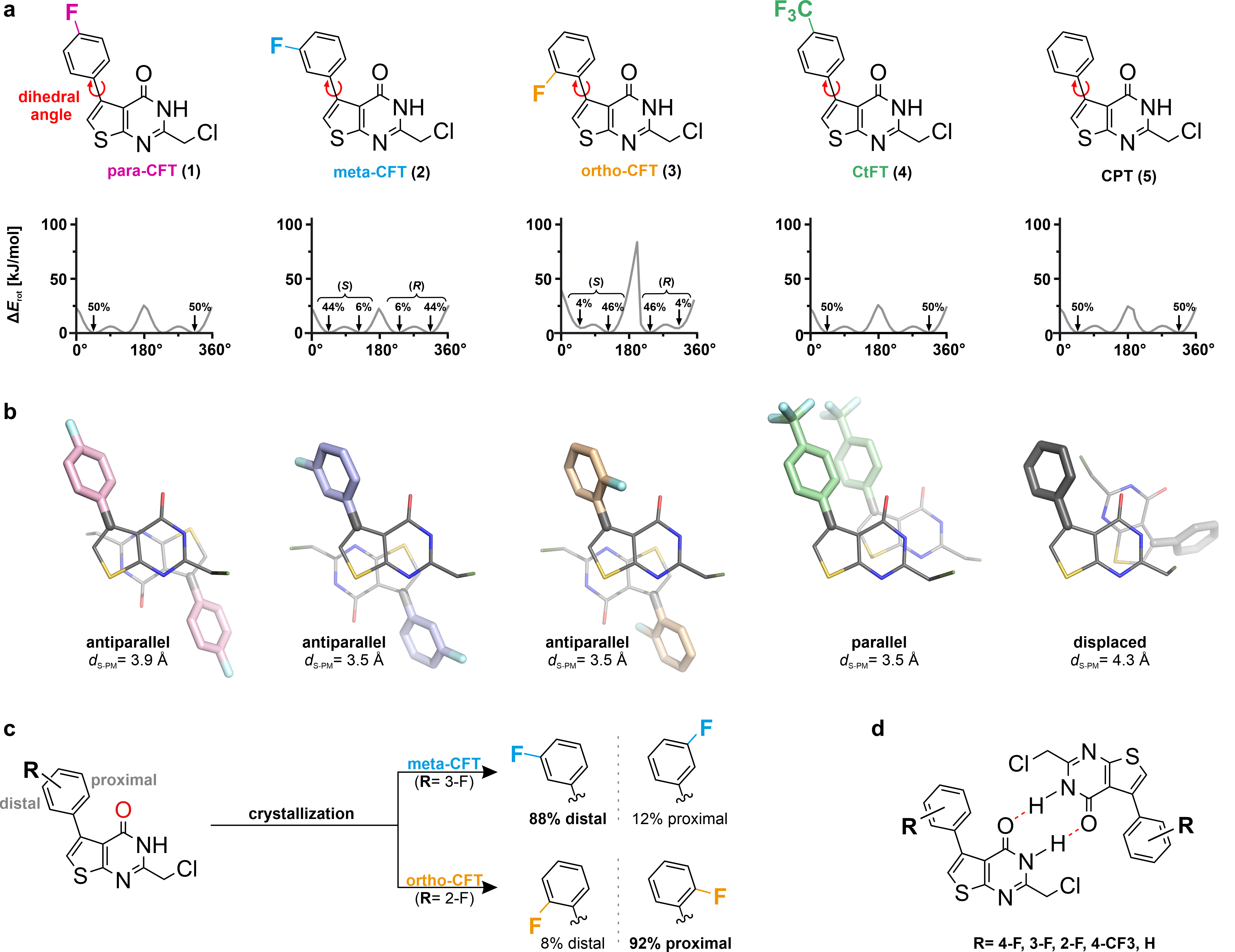

**Supplementary Fig. 9: Density functional theory (DFT) calculations and crystal structures of isolated compounds 1-5.** **a**, Structural formulas and energetic profiles of rotamers of compounds **1-5**. Beginning with the rotamer depicted (= 0°), the dihedral angle between phenyl and thiophene was increased in steps of 10° and the respective rotamer’s energy was calculated using DFT. Ortho-CFT (**3**) shows a rotational barrier of ~80 kJ/mol at 180° due to repulsive forces between the molecule’s fluorine and oxygen. Rotamers seen by X-ray crystallography for the isolated compounds (b, c) are indicated with arrows, the respective populations are noted in %. Axial pro-chirality is indicated above. The dihedral angle observed for isolated para-CFT (47°) agreed with the dihedral angles observed for Tpx-bound para-CFT (47° and 52° for chains A and B, respectively), and was therefore used to set up MD simulations of monomeric and dimeric Tpx/inhibitor complexes. **b,** X-ray crystal structures of isolated compounds **1-5**. The thienopyrimidinone core is shown in grey, and phenyl moieties of para-CFT in pink, meta-CFT in cyan, ortho-CFT in tan, CtFT in green and CPT in grey. Sulfur is shown in yellow, and fluorine in light blue. In the crystal unit cell, inhibitors were found to stack as antiparallel (**1-3**), parallel (**4**) or “displaced” (**5**) dimers. In the antiparallel orientation, the thienopyrimidinone rings are turned by 180° respective to each other. This allows the thiophene rings to stack and, typical for a face-to-face π-stacking interaction^16–18^, adopt a distance of 3.5 to 3.9 Å between the sulfur and the pyrimidinone ring centroid (*d*_S-PM_). Similarly, in the Tpx/para-CFT dimer structure, the *d*_S-PM_ between two parallelly oriented para-CFT molecules is 3.6 Å.^1^ Interestingly, we observed a similar *d*_S-PM_ for isolated CtFT (**4**), even though this compound crystallized in a parallel orientation. This suggests that π-stacking between thiophen-sulfur and pyrimidinone is a reoccurring, energetically favored interaction of these thienopyrimidinone-scaffold molecules. CtFT (**4**) is the only compound with interacting phenyl moieties due to the parallel stacking mode in the crystal structure. The electron-deprived phenyl rings display an ~85° angle with a phenyl/phenyl centroid distance of 4.9 Å, suggestive of stabilization via T-shaped π-interactions.^16–18^ CPT (**5**) molecules show the least aromatic overlap resulting in the “displaced” orientation. **c**, Population of the respective fluorophenyl ring rotamers in meta- and ortho-CFT crystal structures relative to the inhibitor’s oxygen atom. The fluorine in ortho-CFT (**3**) is mainly oriented towards the inhibitor’s carbonyl group (proximal, 92%), which could be facilitated by a fluorine lone pair interaction with the carbonyl carbon (nb/π*‑interaction).^19^ The opposite orientation (distal) is more populated in meta-CFT crystals (88%) which might result from repulsive interactions between the inhibitor’s fluorine and oxygen atoms. **d,** Amide groups in all compounds interacted with a symmetry mate via hydrogen bonds (red dashed lines).

The CCDC deposition numbers of the structures of isolated compounds 1-5 are 1862408, 2421180, 2421181, 2421182, and 2423222.

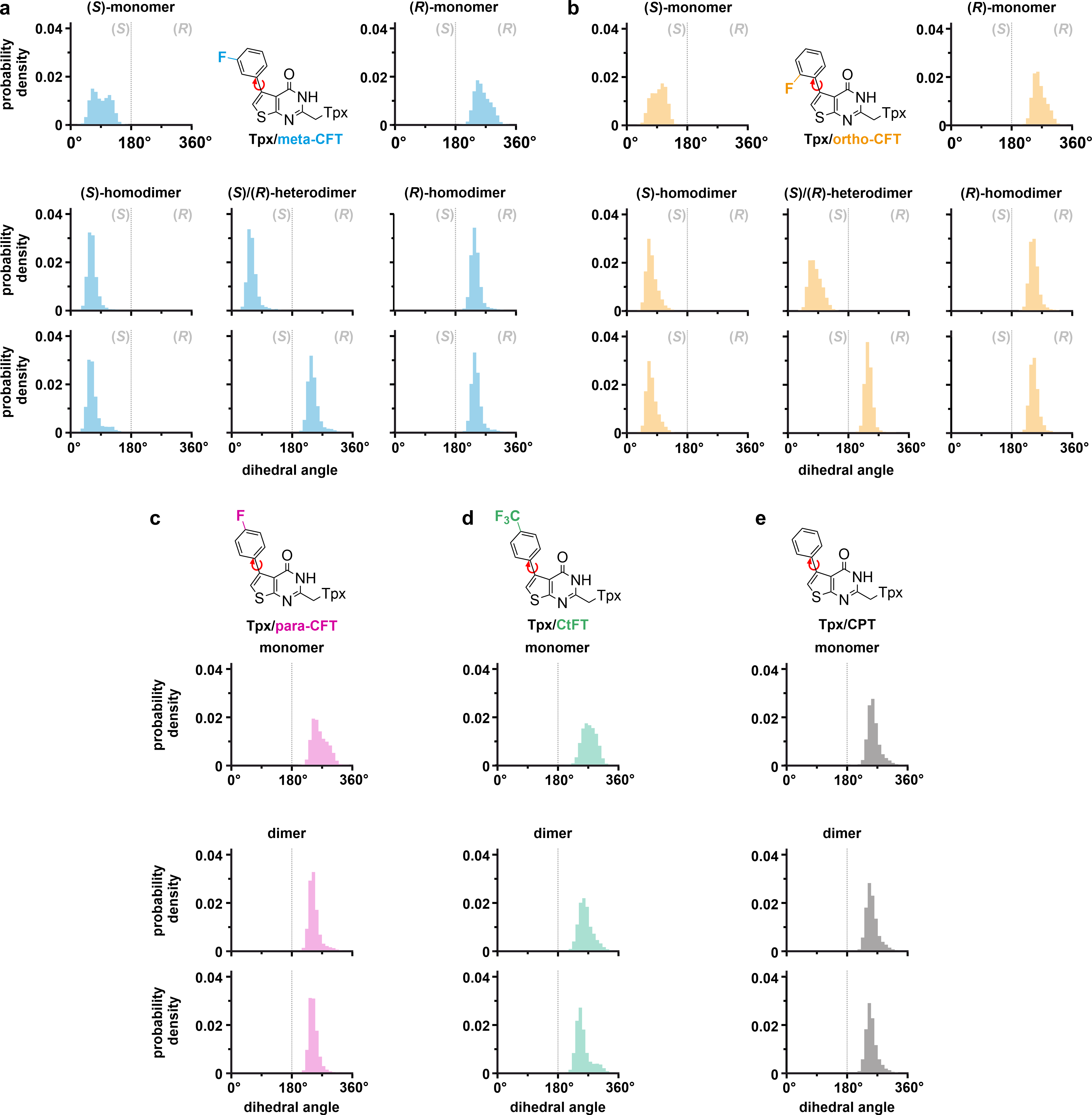

**Supplementary Fig. 10: Phenyl/thiophen dihedral angle distributions of monomeric and dimeric Tpx WT/inhibitor complexes in silico.** Five independent 100 ns molecular dynamics simulations were carried out per atropisomer for **a,** Tpx**/**meta-CFT, **b**, Tpx/ortho-CFT, **c**, Tpx/para-CFT, **d**, Tpx/CtFT, **e**, Tpx/CPT. In all cases, the orientation of protein chain A (monomers) or chain A and chain B (dimers) and the inhibitor molecule of the crystal structure of the Tpx/para-CFT complex (PDB: 6GXG^1^) were used as starting models (see main text for details). The number of frames displaying a specific rotamer over the course of the simulations is shown for each inhibitor. For Tpx/inhibitor complexes with meta- and ortho-CFT (**2**, **3**), the (*S*)- and (*R*)-atropisomers with fluorine pointing towards the binding pocket (“F-in”) or towards the solvent (“F-out”) were simulated. For Tpx dimers, this also included “homodimeric” complexes with both inhibitors in the same, and “heterodimeric” complexes with inhibitors in opposite directions. In all simulations, both in the monomer and dimer, rotation of the phenyl substituent was found to be restrained by the protein, and the accessible dihedral angles were limited to the respective ensemble between the rotational barriers at 0° (or 360°) and 180° (see also **Supplementary Fig. 9**). This suggests that once Tpx/inhibitor complexes with meta- and ortho-CFT (**2**, **3**) are formed, they are chiral. However, the simulations do not allow to gauge which atropisomer is energetically more stable and, even though we did not observe racemization throughout the 100 ns MD simulations, we cannot rule out that (*R*)- and (*S*)-atropisomers are interconverted over a longer time. Of note, most dihedral angle distributions are narrower in the dimer compared to the respective monomeric Tpx/inhibitor complex, indicating that dimerization further restrains rotation of the inhibitor’s phenyl substituent.

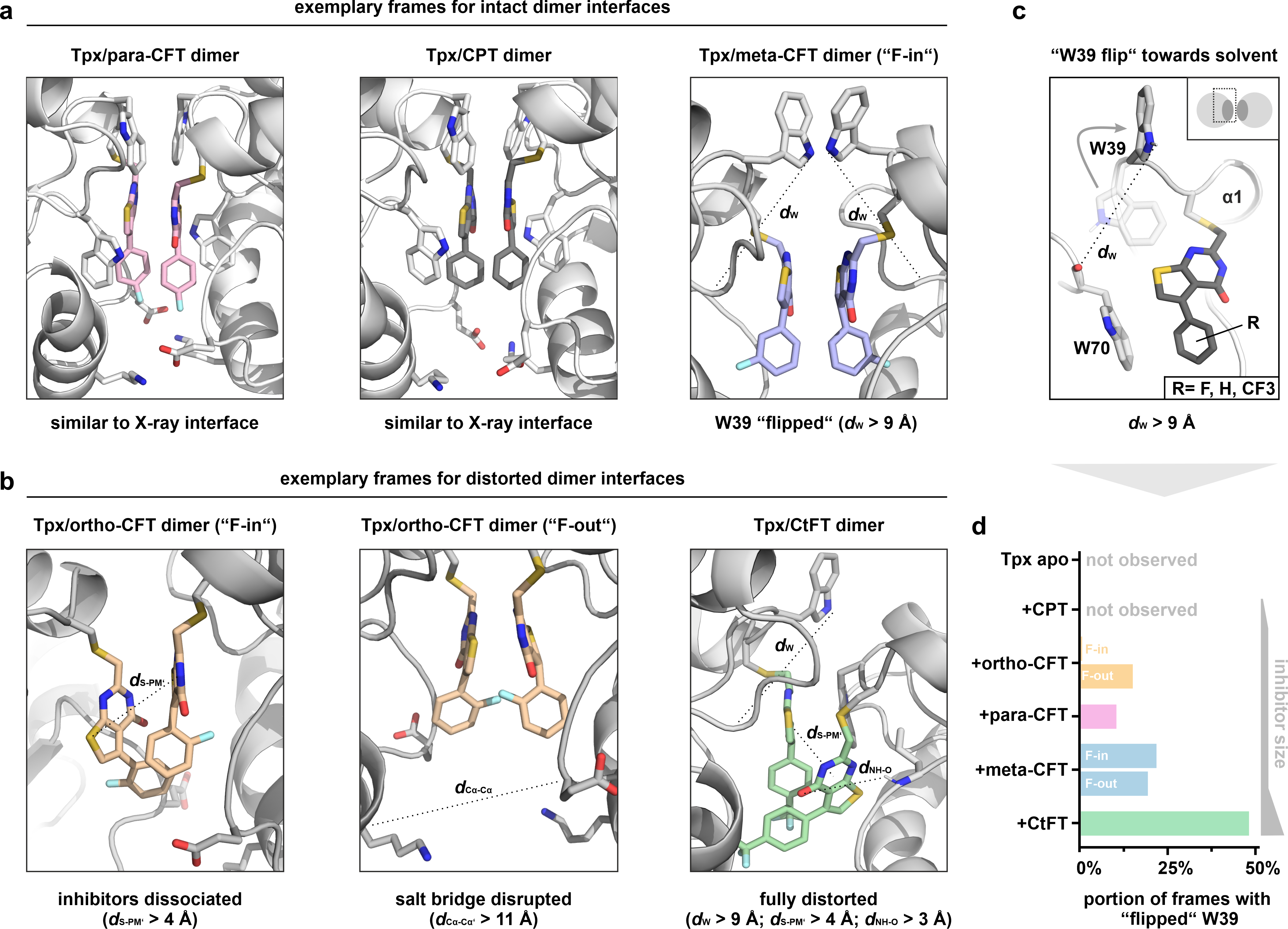

**Supplementary Fig. 11: Exemplary frames from MD simulations of dimeric Tpx/inhibitor complexes showing intact and distorted dimer interfaces.** **a-b,** Exemplary MD-frames of intact (**a**) and distorted (**b**) Tpx dimer interfaces with inhibitors **1**-**5**. Shown are selected interactions within the dimer interface. Exceeding a critical threshold (values indicated below figure) resulted in interface distortion and partial complex dissociation. Loss of contacts deemed relevant for stable dimer formation are highlighted by dashed lines: inhibitor molecule interactions (*d*_S-PM’_ between thiophene sulfur and pyrimidinone centroid), charged residues forming salt bridges (*d*_Cα‑Cα’_ between K102/E107’), the inhibitor and the protein binding pocket (*d*_NH-O,_ between the thienopyrimidinone core and the I109 backbone NH) (see also **Fig. 3c,** main text, **movies 1-6**, and Extended Material and Methods for details). For the strong Tpx dimerizers para-CFT, meta‑CFT and CPT (**1-2, 5**), we did not observe dissociation, and the dimer interfaces, resembling our Tpx/para-CFT dimer crystal structure 6GXG), were stable throughout 100 ns MD simulations. For the weak Tpx dimerizers ortho-CFT and CtFT (**3**, **4**), distortions of the dimer interface were frequently observed (e.g. inhibitors did not stack and/or the salt bridge could not form, quantified in **Fig. 3d**). We also monitored the distance between the two tryptophan residues W39 and W70 within the same protomer (indicated by *d*_W_ representing the distance between the W39 indole NH and the backbone oxygen of W70). Here, in a fraction of frames, the W39 side chain adopted a solvent exposed stance (exemplarily shown for Tpx/meta-CFT dimers in the “F-in”-orientation), that still allowed a dimer-productive interaction. **c**, Illustration of the “W39 flip”, quantified by the distance *d*_W_ (criterion *d*_W_ > 9 Å), which occurred in multiple Tpx dimer MD simulations but did not disrupt Tpx dimers. The “W39 flip” was also observed for corresponding monomers (**Supplementary Fig. 17**). **d**, The number of MD simulation frames where W39 adopted a “flipped”, solvent exposed orientation is shown in a bar graph for each inhibitor. Occurrence of a “W39-flip” correlated with inhibitor size (non-fluorinated < monofluorinated < trifluoromethylated inhibitor) but not Tpx dimer affinity (high portion of frames for both the least efficient (CtFT, **4**, green bar, dimer *K*_D_= 410 ± 56 µM) and the best dimerizer (meta-CFT, **2**, blue bar, dimer *K*_D_= 3.7 ± 1.5 µM)). Interestingly, the capability of ortho-CFT (**3**) to displace the W39 side chain greatly varied depending on whether the fluorine atom pointed towards the binding pocket (“F-in”) or the solvent (“F-out”), and indicated that a fluorine in ortho-position, pointing towards the binding pocket, weakens inhibitor adsorption to the protein (also observed in MD simulations of monomeric Tpx/ortho‑CFT complexes, see **Fig. 5e**, **second panel**, and **Fig. 6b**).

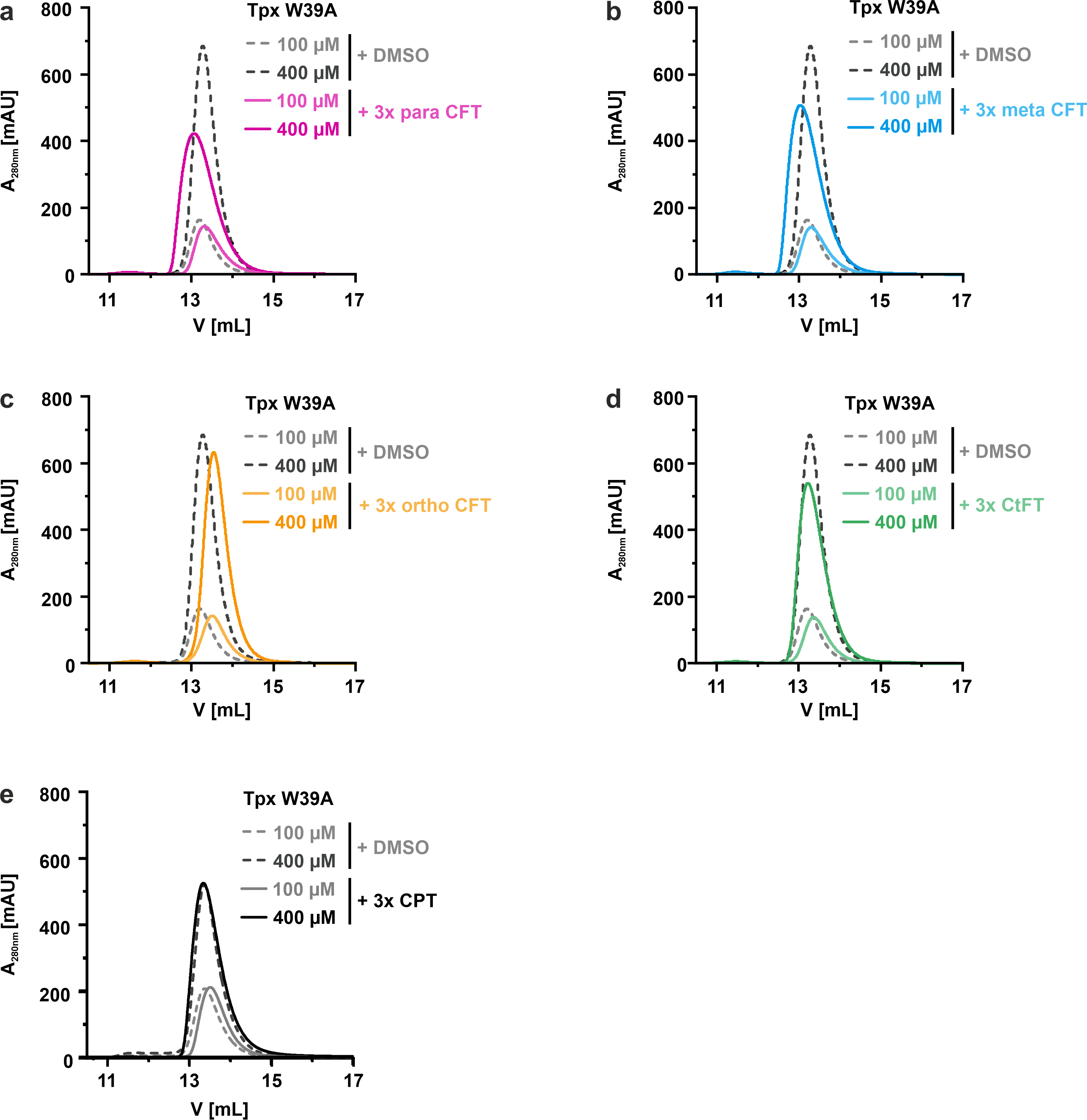

**Supplementary Fig. 12: Tpx W39A remains monomeric upon binding of compounds 1-5. a-e,** Size exclusion chromatography (SEC) of Tpx W39A with three-fold molar excess of compounds **1-5** at 100 µM and 400 µM protein concentration. As a control, the elution profiles of reduced Tpx W39A with 7.5% DMSO (100 µM protein) or 30% DMSO (400 µM protein) in the absence of inhibitor were also recorded (n= 1). In all cases, the elution volumes were highly similar, indicating that the Tpx W39A mutant does not appreciably dimerize under the tested conditions.

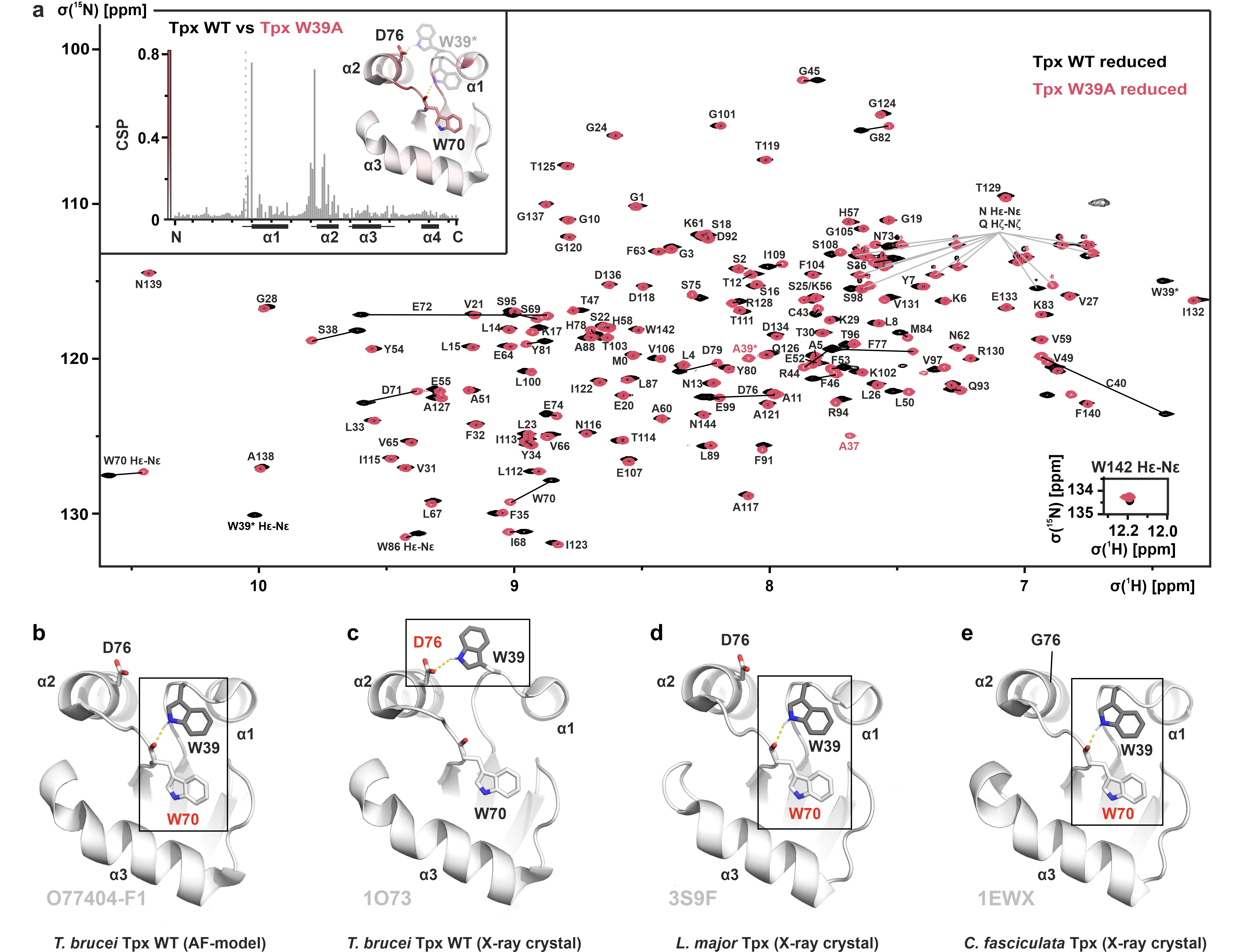

**Supplementary Fig. 13: Backbone NMR assignment of Tpx W39A and W39 side chain flexibility. a,** ^1^H, ^15^N-HSQC spectra of Tpx WT (black) and Tpx W39A (red) in the reduced state, and mutation-induced chemical shift perturbation (CSP) analysis (inset). The backbone assignment for Tpx W39A could be partially obtained by transferring our previous assignment for the Tpx WT^15^, and was verified and completed through additional 3D experiments (BMRB entry 53259). Resonances from the mutated residue 39 are marked with an asterisk. The inset shows the per residue CSP introduced by the W39A mutation which was mapped onto the protein sequence (left) and a structural model of Tpx (right) which is based on both the Alpha Fold model (AF-O77404-F1-model_v4^6,7^) and the *T. brucei* Tpx crystal structure (1O73^10^) to consider the structural ambiguity of residues 38-40 which is particularly pronounced for the W39 side chain orientation (**b** and **c**, respectively). The CSP plot (left) starts with residue G1 and ends with N144 (indicated with “N” and “C”, respectively), ticks indicate increments of ten, starting with G10. The newly appearing signal for residue A37 (which is invisible in the WT spectrum) is indicated with a dashed line. The Tpx α‑helices are indicated by black cylinders on the x-axis and adjacent loops with high CSP are indicated by lines. The tertiary structure model (right) shows W39 in two different conformations where it either forms a hydrogen bond to the W70 backbone (yellow dashed line, see **b**) or the D76 side chain oxygen (see **c**), which is in good agreement with residues that undergo the highest CSP. Heteroatom color scheme: N: blue; O: red. **b-e**, AlphaFold model of *T. brucei* Tpx WT (AF-O77404-F1-model_v4^6,7^), and crystal structures of Tpx WT from *T. brucei* (1O73^10^), *L. major* (3S9F^8^) and *C. fasciculata* (1EWX^9^). Side chains of W39, W70, and respective residues 76 are depicted as sticks (W39 in grey, W70 and residues 76 in white). In the *T. brucei* Tpx WT crystal structure (**c**), W39 adopts a conformation where its side chain is in a hydrogen bond with D76 (*d*= 2.6 Å, yellow dashed line) and fully solvent exposed, i.e. energetically unfavorable for a hydrophobic amino acid in aqueous solution. This clearly differs from the AlphaFold prediction (**b**) where W39 forms a hydrogen bond with the W70 backbone oxygen (*d*= 2.1 Å). As shown above in **Supplementary Fig. 11 c, d**, the W39 side chain also did not adopt a solvent exposed stance throughout a 100 ns MD simulation of the apo state Tpx. Similarly, in crystal structures of orthologous Tpx from *L. major* and *C. fasciculata*, W39 is adsorbed to the protein surface and forms a hydrogen bond with the carbonyl backbone of W70 (*d*_w_= 2.4 Å and 2.0 Å, respectively, yellow dashed lines) regardless of the presence of an aspartate residue at position 76. Taken together, we thus hypothesize that in solution, the W39 side chain of unmodified *T. brucei* Tpx WT is predominantly hydrogen-bonded to the residue W70 backbone and that the solvent-exposed W39 side chain conformation in the available apo state X-ray structure (1O73^10^) reflects only a very small fraction of the conformational ensemble in solution.

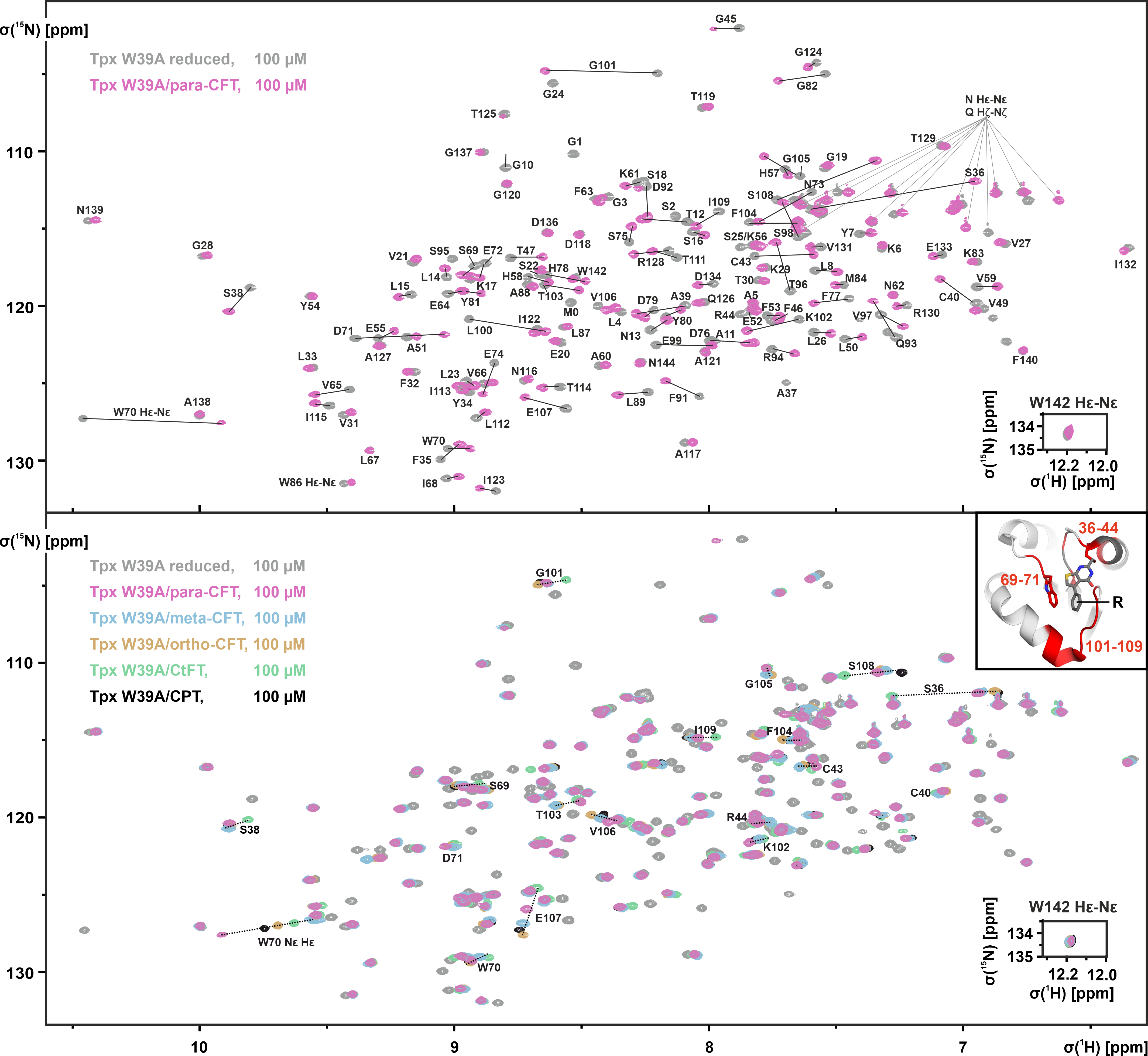

**Supplementary Fig. 14: Chemical shift perturbation of monomeric Tpx W39A upon binding of compounds 1-5. a,** ^1^H, ^15^N-HSQC spectra of Tpx W39A in the reduced (black) and para-CFT-bound state (pink). The backbone assignments for reduced and para-CFT-bound Tpx W39A were transferred from our previous assignment for the reduced Tpx WT^15^ and, where necessary, verified through additional 3D experiments. **b,** ^1^H, ^15^N-HSQC spectra of ^15^N-labeled Tpx W39A in the reduced (grey) and inhibitor-bound states (black, colored). The backbone assignments for Tpx W39A with bound inhibitors (**2-5**) were transferred from our assignments of para-CFT-bound Tpx W39A (see **a**) and, where necessary, verified through additional 3D experiments. Major CSPs are indicated by dotted lines and mostly belong to residues within the Tpx binding site (inset based on the Tpx/para-CFT crystal structure 6GXG^1^, residues affected by inhibitor binding are shown in red). For CSP mapping onto the protein sequence and structure, see **Fig. 4c, d**.

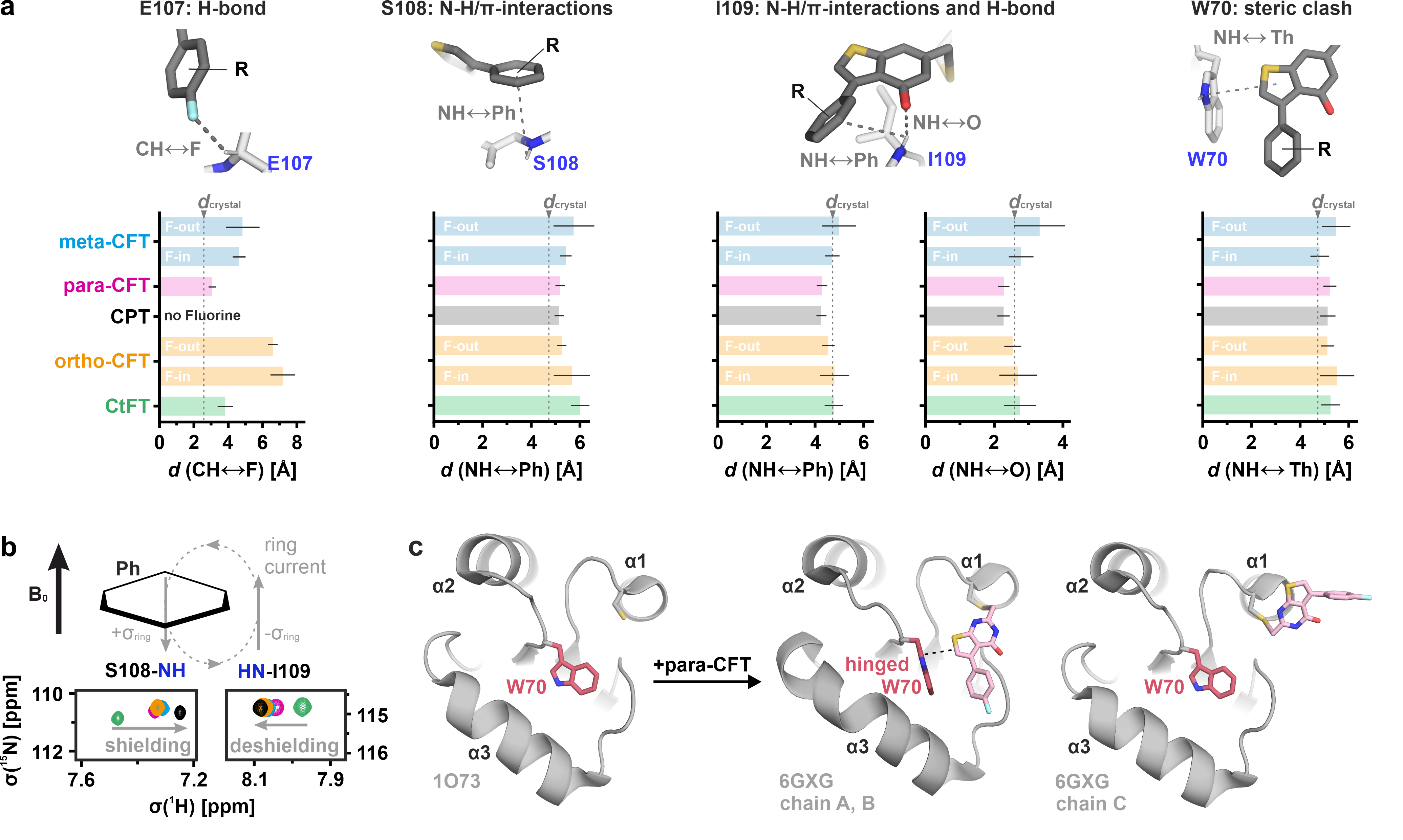

**Supplementary Fig. 15: Structural basis of inhibitor interactions with monomeric Tpx W39A. a**, Analysis of MD simulations (100 ns, 5 replicates) of monomeric Tpx W39A/inhibitor complexes. Simulations were set up using chain A of the Tpx/para-CFT crystal structure (PDB: 6GXG^1^) and replacing W39 with alanine. The mean distances *d* between Tpx W39A/inhibitor contacts, including their standard error (n=5), are shown as bar graphs. For comparison, the corresponding distances observed in the Tpx WT/para-CFT crystal structure (*d*_crystal_) are indicated by a dashed line. For clarity, contacts are visualized on top (copied from **Fig. 4b**, hydrogen bonds and aromatic interactions between Tpx residues and inhibitor moieties (fluorine, phenyl, oxygen, and thiophen) are indicated by dashed lines). Throughout the simulations, all inhibitors (**1-5**) remained in a pose similar to the Tpx/para-CFT crystal structure (*d*_crystal_), with slight deviations for meta-CFT and CtFT (**2**, **4**), showing that the crystal structure can serve as a structural template to model Tpx W39A. As expected, the distance between the respective inhibitor fluorine atom(s) to the C_α_-proton of E107 (i) varied the most due to the different fluorination patterns (*d*_para_<*d*_meta_<*d*_ortho_). Heteroatom color scheme: N: blue; O: red; S: yellow; F: light blue. **b**, The backbone amide resonances of S108 and I109 displayed opposing chemical shift trajectories with inhibitors **1-5**. We propose that an applied magnetic field B_0_ induces an orthogonal phenyl ring current^20^ that shields the backbone NH of S108, which is located underneath the phenyl ring, and deshields the I109 backbone NH next to it. As expected, these effects are least pronounced for the electron deprived phenyl ring carrying a trifluoromethyl group (CtFT **4**, green), most pronounced for the non-fluorinated inhibitor CPT (**5**, black), and nearly identical for the mono-fluorinated inhibitors para-, meta- and ortho-CFT (**1-3**, pink, blue, orange). **c,** Crystallographic data on the Tpx W70 side chain orientation in the apo (1O73)^10^ and para-CFT (**1**)-bound states (6GXG^1^, chain A).^1^ Once the inhibitor adsorbs to the protein surface it induces a ~90° rotation of the W70 side chain, pushing it towards the α3-helix. W70 can thus be considered a “wing gate”^21^, explaining the consequences of inhibitor binding for the chemical environment and the resulting NMR chemical shift of the W70 side chain NH (see **Fig. 4b**, main manuscript). In contrast, mere binding of the inhibitor to C40 does not seem to be sufficient to induce the structural changes in the wing gate, as illustrated by our crystal structure with a solvent exposed, bound inhibitor (6GXG^1^, chain C, “wing gate closed”). Here the orientation of W70 matches that seen in the apo state crystal structure (1O73^10^). In all MD simulations of Tpx/inhibitor complexes, which were set up with the “hinged” W70 pose from the Tpx/para-CFT dimer structure (6GXG^1^, chains A, B, “wing gate open”), the overall W70 orientation did not transition to the apo orientation (seen in crystal structures 1O73^10^ and 6GXG^1^, chain C, “wing gate closed”) (see also **Supplementary Fig. 17**, **movies 1-14**).

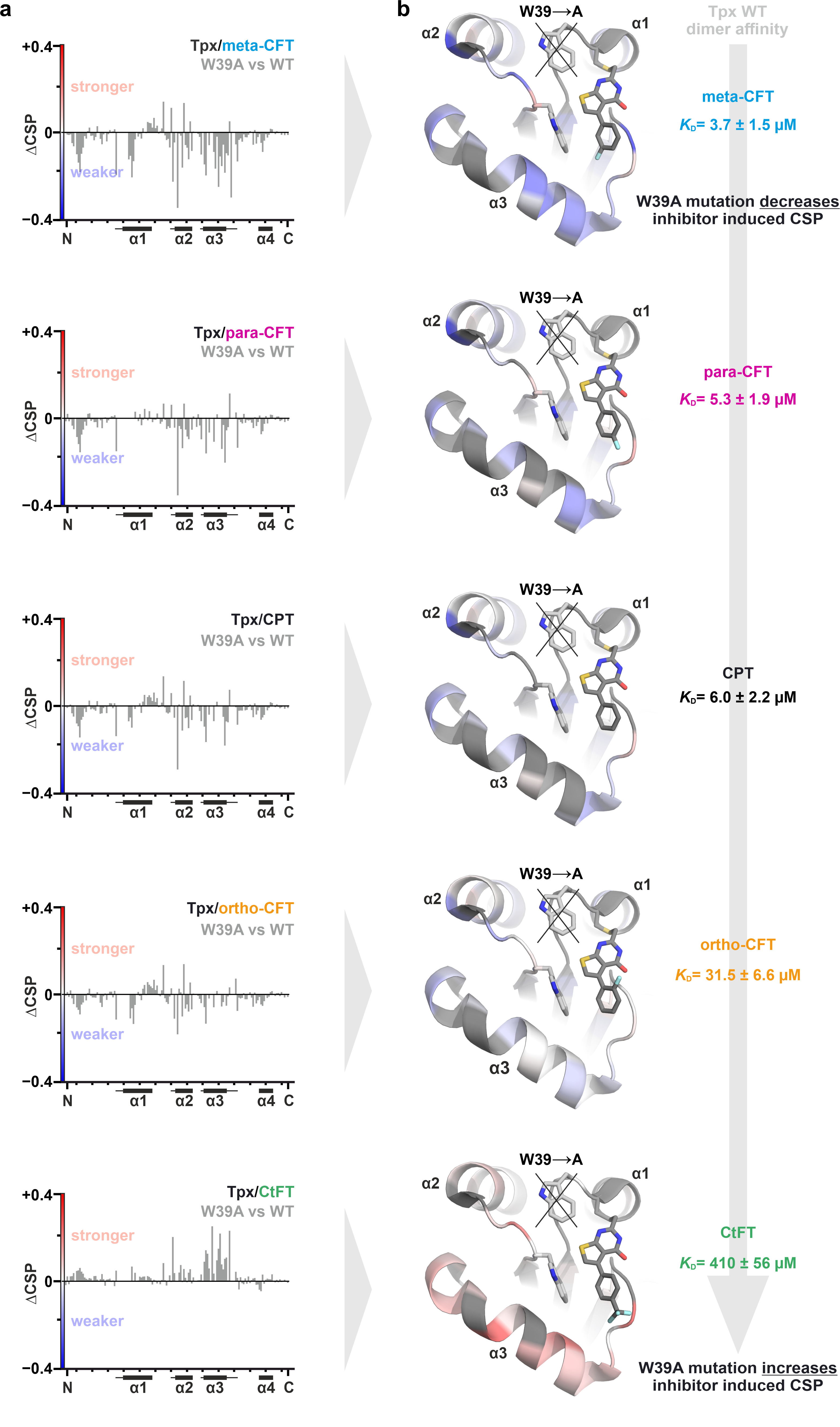

**Supplementary Fig. 16: Comparison of NMR chemical shifts of inhibitor-bound Tpx W39A and Tpx WT. a**, Inhibitor-induced CSP of Tpx WT were subtracted from the inhibitor-induced CSP of Tpx W39A to yield ΔCSP for each inhibitor (**1**-**5**). ΔCSP is shown as bar graphs (left), starting with residue G1 and ending with N144 (indicated with “N” and “C”), ticks indicate increments of ten, starting with G10. Residues without evaluable NMR signals (e.g. due to peak broadening) were arbitrarily set to 0. **b**, ΔCSP mapped onto the inhibitor binding site of Tpx (model based on the Tpx/para-CFT crystal structure 6GXG^1^, chain A). Residues with no ΔCSP are shown in white, and residues without evaluable NMR signals (e.g. due to peak broadening) are depicted in grey. For the strong dimerizer meta-CFT (**2**), ΔCSP is mainly negative (shown in blue) because dimerization, which strongly contributes to the overall CSP, is suppressed in Tpx W39A. Conversely, ΔCSP for the monomeric Tpx WT complex with CtFT (**4**) is positive (shown in red), indicative of a stronger inhibitor interaction with the protein surface once the W39 indole ring is removed via mutation. This suggests a gatekeeping character for Tpx W39 (see main text for details, the figure for CtFT (bottom) is replicated from **Fig. 5a** to allow a side-by-side comparison across inhibitors). Heteroatom color scheme: N: blue; O: red; S: yellow; F: light blue.

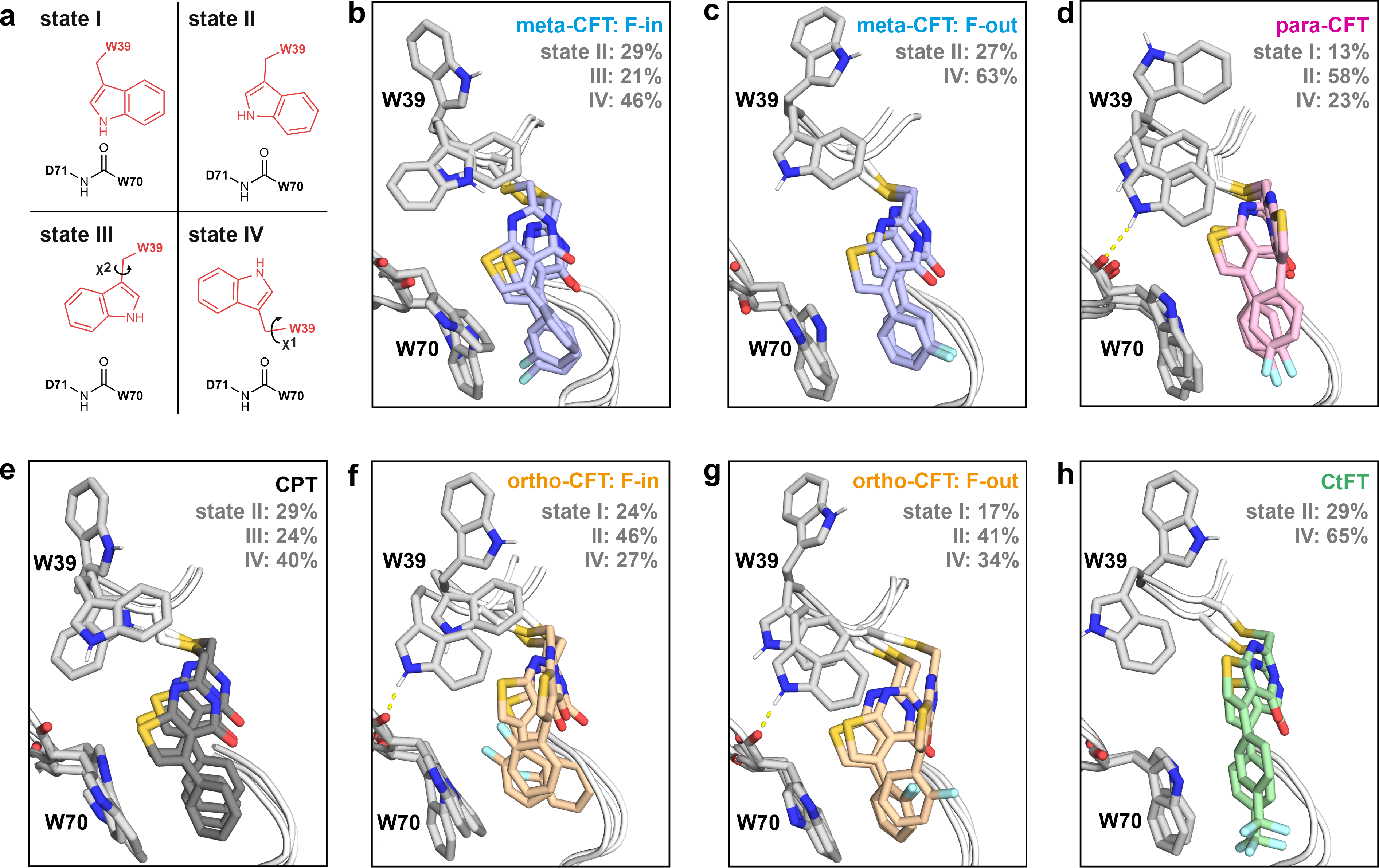

**Supplementary Fig. 17: Dynamics and inhibitor interactions with aromatic surface residues W39 and W70. a,** Schematic depiction of Tpx W39 states (**I-IV**) observed in MD simulations of monomeric Tpx/inhibitor complexes (100 ns, n=5) which were set up using the crystal structure of Tpx WT with bound para-CFT (PDB: 6GXG^1^) as a starting model. State **I** shows W39 in a hydrogen bond with the W70 backbone, which is missing in state **II**. In state **III**, the W39 indole side chain is flipped “sideways”, and in state **IV** “upwards” which leaves the indole ring fully solvent exposed (see also **Supplementary Fig. 13**). For quantification of states **I-IV**, we analyzed the distance between the Tpx W39 indole NH and the backbone oxygen of residue W70 (see **Fig. 5** and extended Material and Methods for details). **b-g:** Snapshots of the Tpx/inhibitor binding interface illustrating W39 states that occurred in at least 10% of all MD simulation frames. Inhibitor fluorination greatly affected the dynamics of the tryptophan side chains and inhibitors in the Tpx binding site, which is also reflected in the relative proportion of states **I-IV** for each inhibitor (shown in the top right of each panel and **Fig. 5e**). In sum, the side chain of W39 is much more dynamic (see also **Supplementary Fig. 13**) than the W70 side chain, which mostly remains in a “hinged” orientation (see **Supplementary Fig. 15**) allowing close inhibitor adsorption to the consensus Tpx binding pocket. Of note, the weak dimerizers ortho-CFT (especially in “F-in” orientation) and CtFT (**3**, **4**) occurred more often in a loosely attached pose (**panels f-h**, **movies 11-13**) than the other inhibitors, which adapted tightly protein-adsorbed poses (**panels b-e**, **movies 8-10, 14**) similar to the inhibitor stance in the Tpx/para-CFT crystal structure (**Supplementary Fig. 15c**). Hydrogen bonds are shown as yellow dashed lines, heteroatom color scheme: N: blue; O: red; S: yellow; F: light blue.

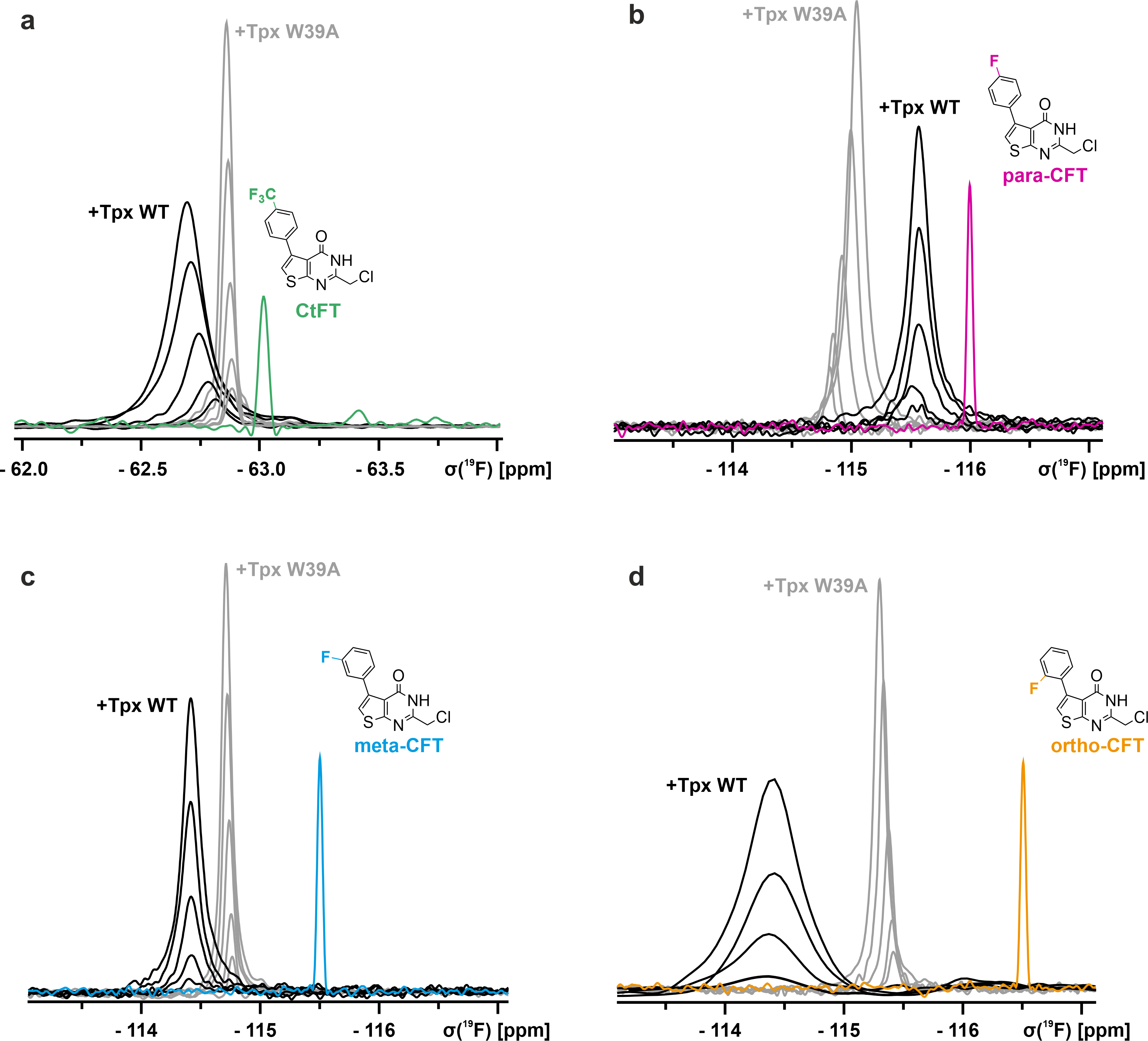

**Supplementary Fig. 18: The ^19^F-NMR signals of free and protein-bound inhibitors provide information about dimer dissociation and bound molecular glue dynamics. a-d,** 1D-^19^F-NMR spectra of free (100 µM, shown in the respective color) and Tpx-bound (750, 500, 250, 100 and 50 µM each; WT: black, W39A: grey) inhibitors **1-4**. ^19^F‑NMR spectra were referenced to TFA and, where DMSO was added, the chemical shift was corrected accordingly (see also **Supplementary Table 4**). The signal of the Tpx WT/ortho-CFT complex at 50 µM is not shown due to poor signal to noise ratio.

**References for Supplementary Information**

1.	Wagner, A. *et al.* Inhibitor-Induced Dimerization of an Essential Oxidoreductase from African Trypanosomes. *Angewandte Chemie International Edition* **58,** 3640–3644; 10.1002/anie.201810470 (2019).

2.	Tormyshev, V. *et al.* Aryl Alkyl Ketones in a One-Pot Gewald Synthesis of 2-Aminothiophenes. *Synlett* **2006,** 2559–2564; 10.1055/s-2006-951484 (2006).

3.	Titchenell, P. M. *et al.* Synthesis and structure-activity relationships of 2-amino-3-carboxy-4-phenylthiophenes as novel atypical protein kinase C inhibitors. *Bioorganic & Medicinal Chemistry Letters* **23,** 3034–3038; 10.1016/j.bmcl.2013.03.019 (2013).

4.	Mahasenan, K. V. *et al.* Exploitation of Conformational Dynamics in Imparting Selective Inhibition for Related Matrix Metalloproteinases. *ACS Medicinal Chemistry Letters* **8,** 654–659; 10.1021/acsmedchemlett.7b00130 (2017).

5.	Adepu, R. *et al.* Facile assembly of two 6-membered fused N-heterocyclic rings: a rapid access to novel small molecules via Cu-mediated reaction. *Chemical communications (Cambridge, England)* **49,** 190–192; 10.1039/c2cc37070k (2013).

6.	Jumper, J. *et al.* Highly accurate protein structure prediction with AlphaFold. *Nature* **596,** 583–589; 10.1038/s41586-021-03819-2 (2021).

7.	Varadi, M. *et al.* AlphaFold Protein Structure Database in 2024: providing structure coverage for over 214 million protein sequences. *Nucleic acids research* **52,** D368-D375; 10.1093/nar/gkad1011 (2024).

8.	Fiorillo, A., Colotti, G., Boffi, A., Baiocco, P. & Ilari, A. The crystal structures of the tryparedoxin-tryparedoxin peroxidase couple unveil the structural determinants of Leishmania detoxification pathway. *PLoS neglected tropical diseases* **6,** e1781; 10.1371/journal.pntd.0001781 (2012).

9.	Hofmann, B. *et al.* Structures of tryparedoxins revealing interaction with trypanothione. *Biological chemistry* **382,** 459–471; 10.1515/BC.2001.056 (2001).

10.	Alphey, M. S. *et al.* Tryparedoxins from Crithidia fasciculata and Trypanosoma brucei: photoreduction of the redox disulfide using synchrotron radiation and evidence for a conformational switch implicated in function. *The Journal of biological chemistry* **278,** 25919–25925; 10.1074/jbc.M301526200 (2003).

11.	Fueller, F., Jehle, B., Putzker, K., Lewis, J. D. & Krauth-Siegel, R. L. High throughput screening against the peroxidase cascade of African trypanosomes identifies antiparasitic compounds that inactivate tryparedoxin. *The Journal of biological chemistry* **287,** 8792–8802; 10.1074/jbc.M111.338285 (2012).

12.	O'Brien, J., Wilson, I., Orton, T. & Pognan, F. Investigation of the Alamar Blue (resazurin) fluorescent dye for the assessment of mammalian cell cytotoxicity. *European Journal of Biochemistry* **267,** 5421–5426; 10.1046/j.1432-1327.2000.01606.x (2000).

13.	Hellberg, U., Ivarsson, J.-P. & Johansson, B.-L. Characteristics of Superdex® prep grade media for gel filtration chromatography of proteins and peptides. *Process Biochemistry* **31,** 163–172; 10.1016/0032-9592(95)00044-5 (1996).

14.	Berger, A. A., Völler, J.-S., Budisa, N. & Koksch, B. Deciphering the Fluorine Code-The Many Hats Fluorine Wears in a Protein Environment. *Accounts of Chemical Research* **50,** 2093–2103; 10.1021/acs.accounts.7b00226 (2017).

15.	Wagner, A., Diehl, E., Krauth-Siegel, R. L. & Hellmich, U. A. Backbone NMR assignments of tryparedoxin, the central protein in the hydroperoxide detoxification cascade of African trypanosomes, in the oxidized and reduced form. *Biomol NMR Assign* **11,** 193–196; 10.1007/s12104-017-9746-7 (2017).

16.	Meyer, E. A., Castellano, R. K. & Diederich, F. Interactions with aromatic rings in chemical and biological recognition. *Angewandte Chemie (International ed. in English)* **42,** 1210–1250; 10.1002/anie.200390319 (2003).

17.	Zhao, Y. *et al.* Conformational Preferences of π-π Stacking Between Ligand and Protein, Analysis Derived from Crystal Structure Data Geometric Preference of π-π Interaction. *Interdiscip Sci Comput Life Sci* **7,** 211–220; 10.1007/s12539-015-0263-z (2015).

18.	Chen, X., Wang, G., Zeng, X., Li, W. & Zhou, M. π–π Stacking and Structural Configurations in Aromatic Thiophene and Fluorobenzene Dimers Revealed by Rotational Spectroscopy. *J. Am. Chem. Soc.*; 10.1021/jacs.5c02401 (2025).

19.	Harry, S. A. *et al.* The Close Interaction of a C-F Bond with an Amide Carbonyl: Crystallographic and Spectroscopic Characterization. *Angewandte Chemie International Edition* **61,** e202207966; 10.1002/anie.202207966 (2022).

20.	Baranac-Stojanović, M. New insight into the anisotropic effects in solution-state NMR spectroscopy. *RSC Adv.* **4,** 308–321; 10.1039/C3RA45512B (2014).

21.	Gora, A., Brezovsky, J. & Damborsky, J. Gates of enzymes. *Chemical reviews* **113,** 5871–5923; 10.1021/cr300384w (2013).
